# Honeybee colony performance affected by crop diversity and farmland structure: a modelling framework

**DOI:** 10.1101/2019.12.17.880054

**Authors:** Juliane Horn, Matthias A. Becher, Karin Johst, Peter J. Kennedy, Juliet L. Osborne, Viktoriia Radchuk, Volker Grimm

**Affiliations:** Helmholtz Centre for Environmental Research – UFZ, Permoserstr. 15, 04318 Leipzig, Germany; Environment and Sustainability Institute, University of Exeter, Penryn Campus, Penryn, Cornwall, TR10 9FE, UK; Leibniz Institute for Zoo and Wildlife Research (IZW) in the Forschungsverbund Berlin e.V., Alfred-Kowalke-Straße 17, 10315 Berlin, Germany; Plant Ecology and Nature Conservation, University of Potsdam, Am Mühlenberg 3, 14476 Potsdam, Germany

**Keywords:** *Apis mellifera*, colony viability, cropping system, decline, forage availability, forage gaps, honeybees, modelling, BEEHAVE, landscape generator, crop diversity

## Abstract

Forage availability has been suggested as one driver of the observed decline in honeybees. However, little is known about the effects of its spatiotemporal variation on colony success. We present a modelling framework for assessing honeybee colony viability in cropping systems. Based on two real farmland structures, we developed a landscape generator to design cropping systems varying in crop species identity, diversity, and relative abundance. The landscape scenarios generated were evaluated using the existing honeybee colony model BEEHAVE, which links foraging to in-hive dynamics. We thereby explored how different cropping systems determine spatiotemporal forage availability and, in turn, honeybee colony viability (e.g., time to extinction, *TTE*) and resilience (indicated by, e.g. brood mortality). To assess overall colony viability, we developed metrics, *P*_H_ and *P*_P,_ which quantified how much nectar and pollen provided by a cropping system per year was converted into a colony’s adult worker population. Both crop species identity and diversity determined the temporal continuity in nectar and pollen supply and thus colony viability. Overall farmland structure and relative crop abundance were less important, but details mattered. For monocultures and for four-crop species systems composed of cereals, oilseed rape, maize and sunflower, *P*_H_ and *P*_P_ were below the viability threshold. Such cropping systems showed frequent, badly timed, and prolonged forage gaps leading to detrimental cascading effects on life stages and in-hive work force, which critically reduced colony resilience. Four-crop systems composed of rye-grass-dandelion pasture, trefoil-grass pasture, sunflower and phacelia ensured continuous nectar and pollen supply resulting in *TTE* > 5 years, and *P*_H_ (269.5 kg) and *P*_P_ (108 kg) being above viability thresholds for five years. Overall, trefoil-grass pasture, oilseed rape, buckwheat and phacelia improved the temporal continuity in forage supply and colony’s viability. Our results are hypothetical as they are obtained from simplified landscape settings, but they nevertheless match empirical observations, in particular the viability threshold. Our framework can be used to assess the effects of cropping systems on honeybee viability and to develop land-use strategies that help maintain pollination services by avoiding prolonged and badly timed forage gaps.

## Introduction

Honeybees (*Apis mellifera L.*) are a key pollinator of insect-pollinated crops and wild plants, with overall insect pollination services being estimated to exceed 153 billion USD in agricultural systems (Gallai et al. 2009). Thus, the ongoing substantial loss of managed honeybee colonies in Europe and USA (e.g. Lee et al. 2015) is of particular concern. Keeping up with the rising demand for insect-pollinated food production seems to be at high risk (Aizen et al. 2008).

As one important stressor, substantial losses of suitable habitats resulting from agricultural intensification and lack of floral resources as alternatives to crops have been assumed to crucially affect honeybee health and hive losses (Naug 2009, Vanbergen et al. 2013, Clermont et al. 2015). Ongoing agricultural intensification over decades has led to simplified annual cropping patterns, predominance of monocultures, increasing silage production, and scarcity of habitats rich in alternative floral resources such as hay meadows, hedgerows, field margins, and grasslands (e.g. Kleijn et al. 2006). Consequently, in many regions after a short period with ample amounts of nectar and pollen provided by mass-flowering crops, there is a forage dearth in early spring and summer (Decourtye et al. 2010, Couvillon et al. 2014).

Although previous field studies using nectar measurements and pollen traps show seasonal patterns in nectar and pollen collection by the bees (e.g. Requier et al. 2015), seasonal dynamics in nectar and pollen provided by intensively managed farmlands and their effects on the honeybee colonies are poorly known. This limits our understanding of how the composition and configuration of agricultural landscapes drives the continuity of food availability in space and time and in turn affects honeybee colony dynamics and survival. Nevertheless, apiculturists have long realized that landscape context is a critical factor for colony success (Sponsler & Johnson 2015).

Temporary shortages in sufficient flowering resources, which are a typical phenomenon in spring and summer in many European agricultural landscapes, can strongly affect colony success (Decourtye et al. 2010). If such food shortages occur in a sensitive phase of colony development, i.e. when the colony is close to achieving maximum brood rearing and adult population size, the colony’s resilience and survival capability are strongly impaired via cascading effects on life stages and tasks (Horn et al. 2016).

Systematically exploring in field experiments how landscape configuration and composition affect colony resilience and viability is not feasible (Henry et al. 2017). We therefore performed corresponding simulation experiments by using a two-step modelling framework. First, we developed a landscape generator (NePoFarm) that generates and calculates, on a daily basis, the nectar and pollen supply of cropping systems scenarios which varied in landscape structure, and the crop species identity, diversity and relative abundance. For this, we had to compile data on the phenology and the quantities of nectar and pollen provided by the major crop species. Second, we fed the resulting spatiotemporal data on nectar and pollen supply into the simulation model BEEHAVE (Becher et al. 2014) to assess honeybee colony performance under the different cropping systems scenarios.

BEEHAVE is the first honeybee model that directly links in-hive dynamics to foraging in realistic landscapes via energy gains (Becher et al. 2013). BEEHAVE is spatially implicit, i.e. the location of nectar and pollen resources and the distance flown by the bees matter, but it represents each field by only one location, its center of gravity. Moreover, BEEHAVE allows the incorporation of spatiotemporal dynamics in nectar and pollen availability by explicitly implementing the flowering periods, the amounts of nectar and pollen and sugar concentration of certain crop species. These dynamics in floral resources are directly linked to the colony development via nectar and pollen intake and consumption. It was considered realistic enough to be recommended as a basis for developing tools for regulatory pesticide risk assessment (EFSA 2015, 2016). BEEHAVE was also validated with data from field studies (Agatz et al. 2019).

The major aims of this study are to present our modelling framework for assessing honeybee colony performance for various cropping systems in a systematic way and to report our first generic findings. The framework will help to improve our understanding of how cropping systems could provide sufficient and sufficiently continuous nectar and pollen supply that meet the colony’s requirements for ensuring viability and how temporary gaps in forage supply affect colony resilience. To introduce the framework and its potential, we assumed hypothetical but still realistic patterns of crop composition and distribution. Semi-natural habitats provide important floral resources even in intensively managed agricultural landscapes. As data about their spatial distribution, species composition and nectar and pollen rewards are sparse we represented them in our modelling framework in an aggregated way.

In this study we apply our modelling framework to the most important European cultivated crops, asking the following questions: (1) What is the minimum diversity of cultivated crops required to meet a colony’s food requirements for continuous food supply and thereby enable viability? (2) Does this minimum depend on the identity of the cultivated crops involved and their relative abundance? (3) How does farmland structure, i.e. spatial configuration and the size distribution of agricultural fields, affect colony viability?

## Methods

Due to the complexity of the question addressed and the tools used, considerable effort went into collecting data for parameterization and for analyzing and testing the model. Most details of the corresponding information are presented in the Supplementary Information (SI) but are relevant for using our modelling framework for different settings and questions. Therefore we provide an overview of the contents of the SI in Table 1.

**Table 1.** Overview of the content of the Supplementary Information (SI).

|  |  |  |  |
| --- | --- | --- | --- |
| S1 | Model parameters, equations and assumptions | Initial parameter settings, weather conditions suitable for foraging, most important model equations | 2 tables<br>1 figure |
| S2 | Farmland maps | How the two farmland maps we prepared to implement different cropping system scenarios<br>S2-1: R script “NePoFarm”<br>S2-2: Input file for NePoFarm | 1 table |
| S3 | Crop species and their nectar and pollen data | Source for selecting and parameterizing crop species (flowering periods, nectar and pollen parameters) as well as for the composition and abundances of the cropping systems considered | 4 tables<br>12 figures |
| S4 | Landscape generator NePoFarm to create scenarios as input files for the simulation model BEEHAVE | Description of NePoFarm with links to corresponding R script and parameter file | 4 tables<br>1 figures |
| S5 | $P_H / P_P$: Indices for comparing forage-induced stress at the colony level | Parameters and equations for calculating these indices | 1 table |
| S6 | Ranking farmland factors by mimicking a local sensitivity analysis procedure | Description of sensitivity analyses and statistical procedures to evaluate their results | - |
| S7 | Simulations - Landscape settings | Sensitivity analyses of nectar and pollen parameters, the distance and size of flowering fields, field size of monocultures, and of semi-natural habitats | 10 tables<br>18 figures |
| S8 | Current limitations | Current limitations of our modelling framework which are due to uncertainty and data and model assumptions. | - |

### The BEEHAVE model in short

The honeybee model BEEHAVE integrates in-hive colony dynamics, in-hive varroa mite population dynamics, mite-mediated disease transmission, and foraging for nectar and pollen in heterogeneous landscapes. It was designed to explore how various stressors and their interactions affect the structure and dynamics of a single honeybee colony (Becher et al. 2014). BEEHAVE is implemented in the freely available software platform NetLogo (Wilensky 1999). The model, its detailed description following the ODD (Overview, Design concepts, Details) protocol (Grimm et al. 2006, 2010), and a user manual are freely available (www.beehave-model.net). Important model assumptions and equations are listed in Tables S1 and S2.

BEEHAVE comprises three modules. The colony module is based on age cohorts and describes, on a daily basis, in-hive colony structure and dynamics driven by the queen’s egg-laying rate, weather, and forage input. The bees’ developmental stage and disease status, available nursing bees and the colony’s honey and pollen stores determine the mortality rates of brood and adult bees. The mite module describes the dynamics of a virus-transmitting varroa mite population within the hive. The foraging module is individual-based and represents the bees’ foraging behavior; it is executed once per day and operates on a time scale of minutes. Weather conditions affect the daily time allocated for nectar and pollen collection. Landscape features, including changes in spatiotemporal availability of nectar and pollen can be updated every day.

In-hive dynamics and foraging are linked via energy and protein budgets: foragers, in-hive bees, and brood require certain amounts of energy and protein provided by nectar and pollen, respectively. These requirements are satisfied with incoming forage and are linked to the production and consumption of nectar and pollen stores. A work force which is too small to care for brood can lead to reduced brood production. The distance of flowering patches to a colony and their nutritional reward determine the energetic efficiency of foraging, which is communicated within the colony via a representation of the waggle dance. Details on the foraging module are provided in Table S2.

### Initial settings

Following BEEHAVE’s default settings, all simulations started on 1^st^ January with an initial colony size of 10,000 worker bees and 25 kg honey, which is the amount of honey needed to let a colony of this size survive until spring. As this study focuses on impacts of farmland structure and crop composition, we considered neither virus-transmitting varroa mite infestation nor its management. Previous simulations found that with an untreated varroa infestation, 50 % of model colonies died after four years even under beneficial foraging conditions (continuous forage supply and 500 m flight distance). Increasing flight distances to the forage patch accelerated colony failure, but the increasing prevalence of the virus in the colony over time in declining bee population had a stronger effect (Becher et al. 2014). Effects of pesticide exposure have been studied elsewhere (Rumkee et al. 2015, Thorbek et al. 2017, Schmolke et al. 2019), and the effects of combined stressors are demonstrated by Henry et al. (2017).

Weather conditions define the maximal daily foraging period in the BEEHAVE model. We chose the annual “Rothamsted (2011)” weather option already provided in the BEEHAVE model (based on a data set from Rothamsted, UK, from 2011) for all scenarios and for each of the simulated years, as it offers favorable foraging conditions from early spring till late autumn without prolonged periods of bad weather to avoid interference with forage gaps (Fig. S1).

As phenological data on species-specific nectar and pollen availability are scarce, we assumed crop species-specific nectar and pollen contents per flower and sugar concentration to be constant over the corresponding flowering period. To represent early spring foraging on days when weather conditions are suitable for foraging flights, we placed a forage patch providing low daily amounts of nectar (1 l) and pollen (0.5 kg) from January until end-March at 1,000 m from the hive, which allows the colony to survive the early spring period (see also Horn et al. 2016). This forage patch represents early flowering plants such as snowdrop (*Galanthus nivalis*), crocus (*Crocus spec*.) and willow trees (*Salix spec*.). The available nectar and pollen amounts at this forage patch can be completely depleted on a given day, so the time a forager needs to collect a full nectar or pollen load (handling time) increases with the degree of forage depletion at this patch.

### Farmland factors for generating cropping system scenarios

We investigated various aspects of cropping systems in terms of farmland structure, crop species identity, and crop diversity represented by number and relative abundance of different crop species (Figs. 1 and 2). These farmland factors are used to generate different scenarios reflecting different cropping systems. We made simplified assumptions about crop rotation and the distribution of semi-natural habitats in agricultural systems as data about floral phenology, pollen and nectar quantities and qualities of plant species of semi-natural habitats are scarce (see Baude et al. 2016).

**Figure 1.**
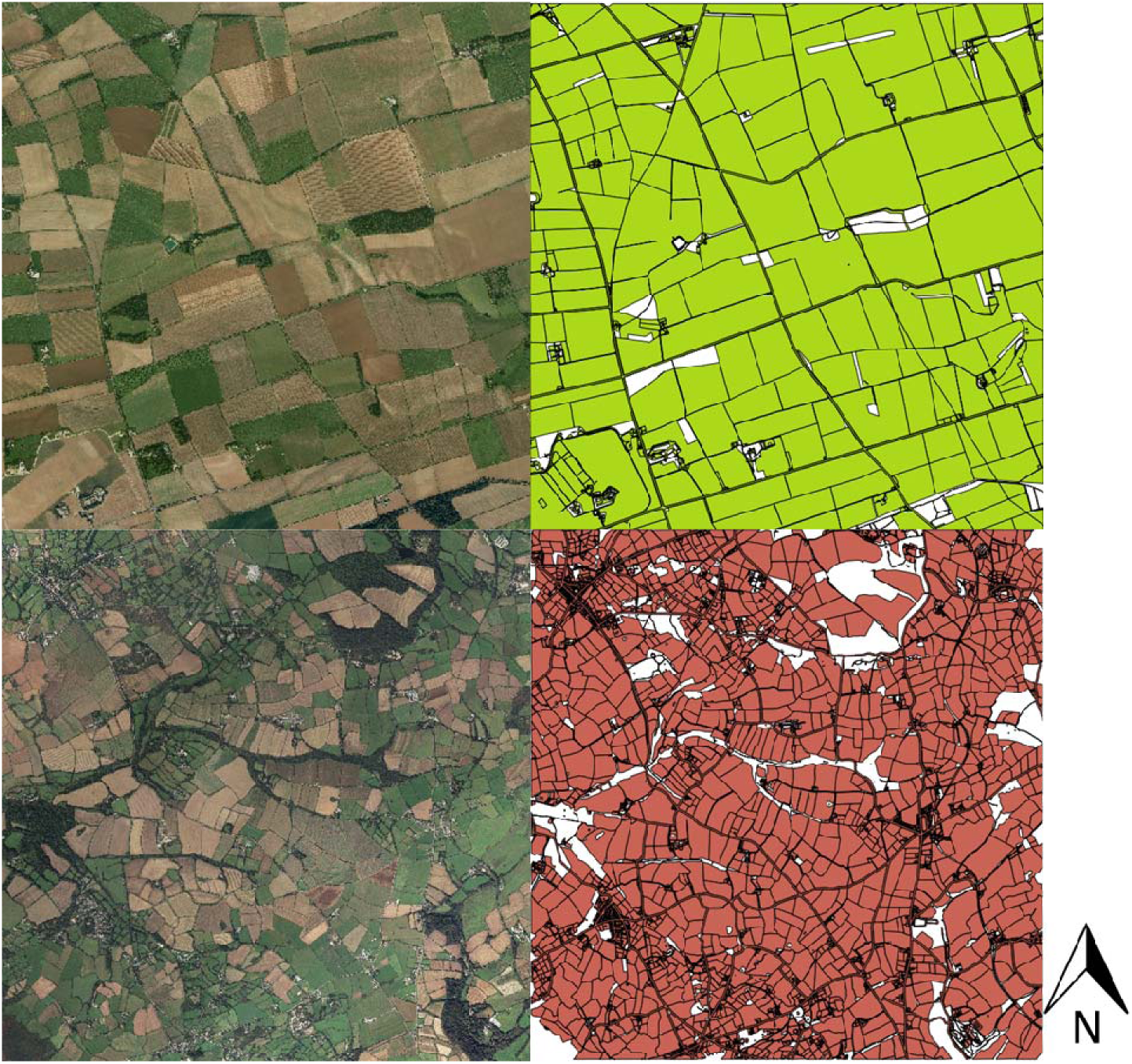
Aerial (left) and raster (right) images of 5 x 5 km maps of the two farmland structures: a simple-structured farmland located in Lincolnshire – Nocton Heath (above) and a complex-structured farmland located in Cornwall – Trenwheal (below) (vector data were taken from OS VectorMap Local^TM^ 2014). These farmland structures are used as templates to generate various scenarios of cropping systems differing in their crop composition and landscape configuration. White and black areas represent buildings, roads, small forested areas, hedgerows and were not considered.

**Figure 2.**
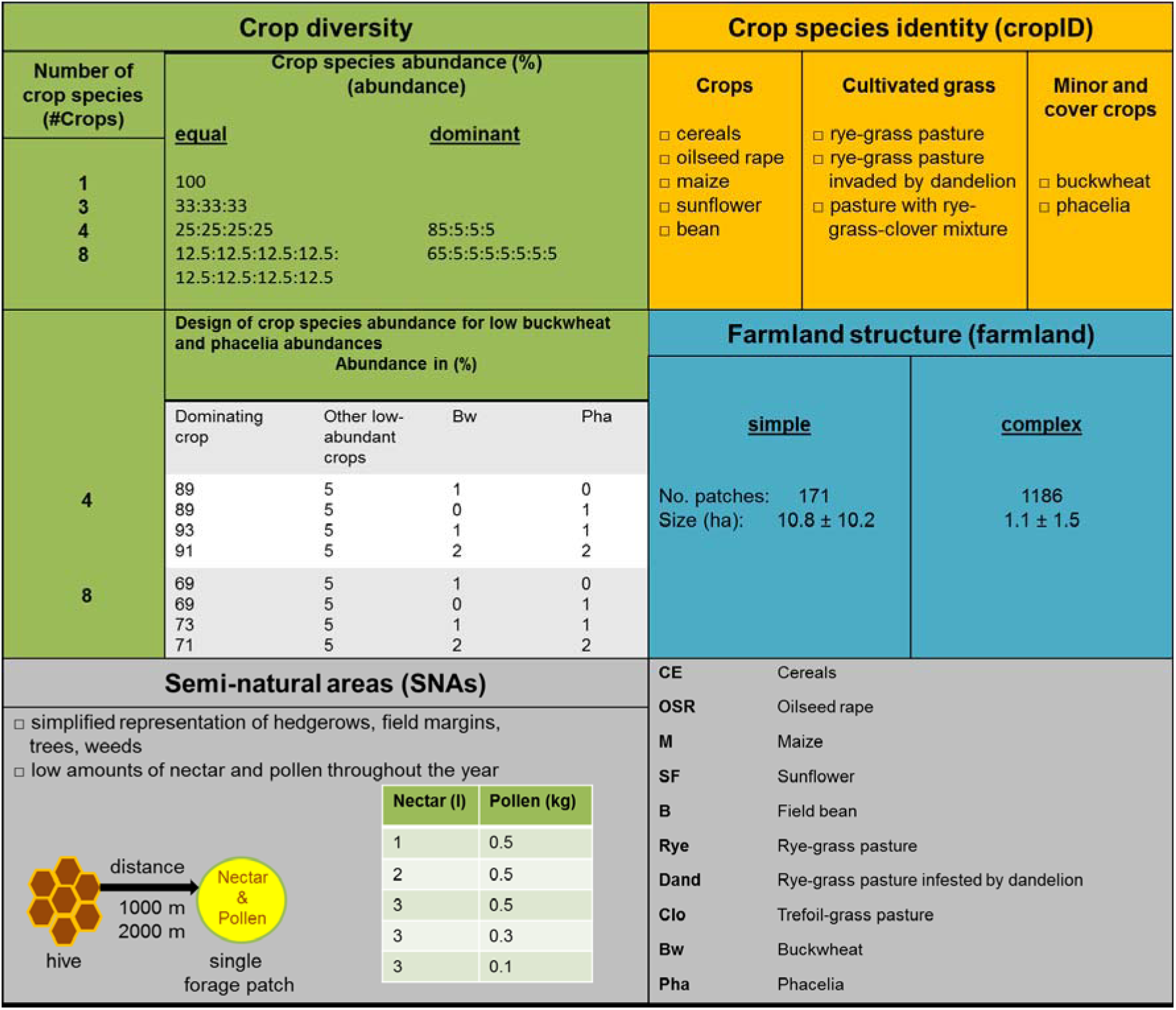
Design of cropping system scenarios. Our design consists of the following variables: number of crop species (*#Crops)*, crop species abundance (*abundance,* in %), crop species identity (*cropID*) and farmland structure (farmland; simple versus complex), and representation of forage providing semi-natural areas (SNAs). Representations of SNAs are simplified in terms of one single patch of one ha size providing nectar amount from 1 up to 3 l and pollen from 100 to 500 g on a daily basis. This single semi-natural habitat patch providing continuous forage over the whole year was located 1000 m or 2000 m away from the hive. Abbreviations of crop species used in the text and the following figures and tables are given.

#### Farmland structure

We used 5 km x 5 km maps of two farmlands differing in their spatial structure (hereafter called ‘farmland structure’) as templates for our semi-realistic landscape settings (Fig. 1). As one landscape we chose an agricultural region with many small agricultural holdings owning small crop fields as common for example in Western Germany and Southwestern England. As second landscape structure we selected an agricultural region with few agricultural holdings owning large fields as common for example in Eastern England and Eastern Germany. The first landscape was a complex-structured farmland characterized by small field sizes (< 1 ha), while the second one was a simple-structured farmland characterized by large field sizes (> 4 ha). The maps were taken from freely available vector data including shape files of area and road characters (OS VectorMap Local^TM^ 2014) for the Trenwheal area in Cornwall (small fields < 1 ha) as a complex-structured farmland and for the Nocton Heath area located in Lincolnshire (large fields > 4 ha) as a simple-structured farmland (Table 2).

**Table 2.**
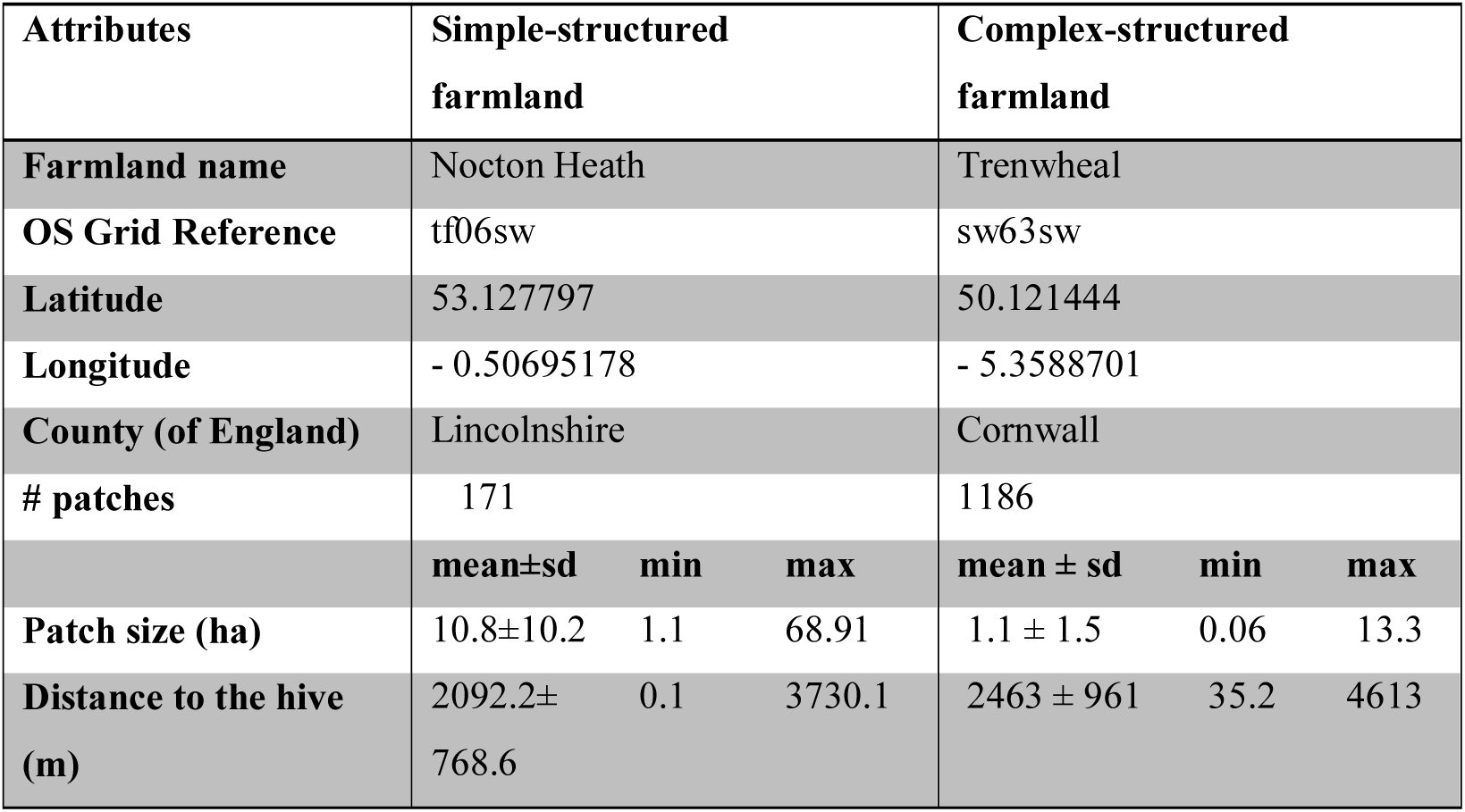
Attributes of both farmland structures.

| Attributes | Simple-structured farmland |  |  | Complex-structured farmland |  |  |
| --- | --- | --- | --- | --- | --- | --- |
| **Farmland name** | Nocton Heath |  |  | Trenwheal |  |  |
| **OS Grid Reference** | tf06sw |  |  | sw63sw |  |  |
| **Latitude** | 53.127797 |  |  | 50.121444 |  |  |
| **Longitude** | - 0.50695178 |  |  | - 5.3588701 |  |  |
| **County (of England)** | Lincolnshire |  |  | Cornwall |  |  |
| **# patches** | 171 |  |  | 1186 |  |  |
|  | **mean±sd** | **min** | **max** | **mean ± sd** | **min** | **max** |
| **Patch size (ha)** | 10.8±10.2 | 1.1 | 68.91 | 1.1 ± 1.5 | 0.06 | 13.3 |
| **Distance to the hive (m)** | 2092.2±768.6 | 0.1 | 3730.1 | 2463 ± 961 | 35.2 | 4613 |

We used R version 3.1.2 (R Development Core Team 2014) to combine all available shape files and to extract all agricultural field boundaries. To rasterize maps we used QGIS version 2.6 (QGIS Development Team 2014). Methodological details are given in the Supplement (S2).

BEEHAVE does not directly use GIS maps but requires, as input, a list of the size, distance and availability of forage for all field patches. To identify the size and location of each field patch and its distance to the hive for both GIS farmland maps, we used the software tool BEESCOUT (Becher et al. 2016). For the simple-structured farmland, 171 field patches were identified with an average size of 10.82 ± 10.21 ha, whereas the complex-structured one contained 1,186 field patches with an average size of 1.09 ± 1.54 ha (Table 2). We placed the hive of the simulated colony always at the same central location (x-coordinate: 157, y-coordinate: 125; map dimensions: 210 × 210 grid cells) for both maps (Table S3).

#### Crop species identity and species-specific nectar and pollen data

We opted to not use the crop species that were actually present in the two landscapes because that would have yielded only two points in the multi-dimensional space of possible crop compositions. Instead, we distinguished ten crop species (Fig. 2), consisting of common European agricultural crops, forage crops as common in intensive pastures, and important European minor and cover crops (Eurostat 2015, FAO 2015). Their use allowed us to explore a wide range of hypothetical cropping system scenarios, which, more generally, reflects real farmlands. Therefore, such a choice renders our analysis a higher level of generality. Details on crop species most common in European agriculture are provided in Tables S4-S5 and in Figures S2-S13.

As agricultural crop species we considered cereals (wheat, rye, barley and oats were pooled together, providing no resources to bees by themselves), oilseed rape (*Brassica napus* L.), maize (*Zea mays* L., providing only low-nutritional pollen to bees), sunflower (*Helianthuus annuus* L.), and field bean (*Vicia faba* L.) as these are the most important crops in European agriculture (FAO 2015, Eurostat 2015). We further considered three different pasture types common in many temperate parts of Europe for grazing, forage harvesting, silage and hay production (EFG 2007). Most pastures are solely sown with rye-grasses (e.g. *Lolium perenne*), whereas other grasses such as *Dactylis glomerata* are rarely sown (EFG 2007, Eurostat 2015). Especially, in permanent pastures, perennial grasses (mainly *Lolium perenne*) combined with white clover (*Trifolium repens*) are preferred (EFG 2007). White clover in a trefoil-grass pasture will ensure a high forage quality improving both energy and protein content for livestock (Hall 1993). For the third pasture type, we included long-lived perennial dandelion (*Taraxacum officinale*) as a valuable forage plant for livestock, as this commonly infests sown rye-grass pastures from nearby roads and field margins (Gibson 1997). Cattles graze on dandelion as readily as on grasses and herbicide control is usually unnecessary for moderate abundances (Bergen et al. 1990).

Further, we implemented important European minor and cover crops. Phacelia (*Phacelia tanacetifolia*) and buckwheat (*Fagopyrum esculentum*) are rapidly growing flowering cover crops in annual cropping systems grown to improve soil quality and are recommended as bee pasture (e.g. Bjorkman and Shail 2010, Decourtye et al. 2010, Baets et al. 2011, Lee-Mader et al. 2014). Buckwheat is also an important minor crop grown for grain-like seeds (Michalova 2001), whereby major producers are France, Ukraine and Poland (FAO 2015, Eurostat 2015).

To translate the identity of crop species (hereafter referred to as ‘*cropID*’) on a given patch to units of nectar and pollen, we extracted floral data (flowering period, flower density, nectar and pollen volume, sugar concentration) from existing literature. Existing data about the amount of nectar and pollen provided by different crop species are highly variable due to high intra-specific variation, and abiotic (e.g. soil nutrition, water content) and biotic factors (e.g. pests, diseases) affecting productivity of floral rewards (e.g. Burke and Irwin 2009). We thus used average values, which are most likely to be linked with the applied beneficial weather conditions throughout the year (“Rothamsted 2011”), where flower production is not limited by bad weather conditions. Furthermore, we assumed constant forage provisioning during the flowering period. Overviews of floral data extracted from literature and equations are given in Table S6 and S7.

We used the flowering period (defined by the start and end days within a year), the amount of nectar and pollen provided per day, and the sugar concentration of each crop species as input data for the honeybee model BEEHAVE. The resulting data of nectar and pollen availability are highly variable but still in the range of nectar productivity for arable land according to Baude et al. (2016). For example, for winter-sown oilseed rape, we defined the flowering period from day 114 (24^th^ April) up to day 144 (24^th^ May) within which it provided 0.001 l/m^2^ nectar at a sugar concentration of 1.7 mol/l (∼ 58%) and 0.349 g/m^2^ pollen per day (Table 3; Fig. 3). Sensitivity analysis of nectar and pollen quantities and qualities of high- and low-rewarding crop species are available in the Supporting information (Table S14; Figs S15–S17).

**Figure 3.**
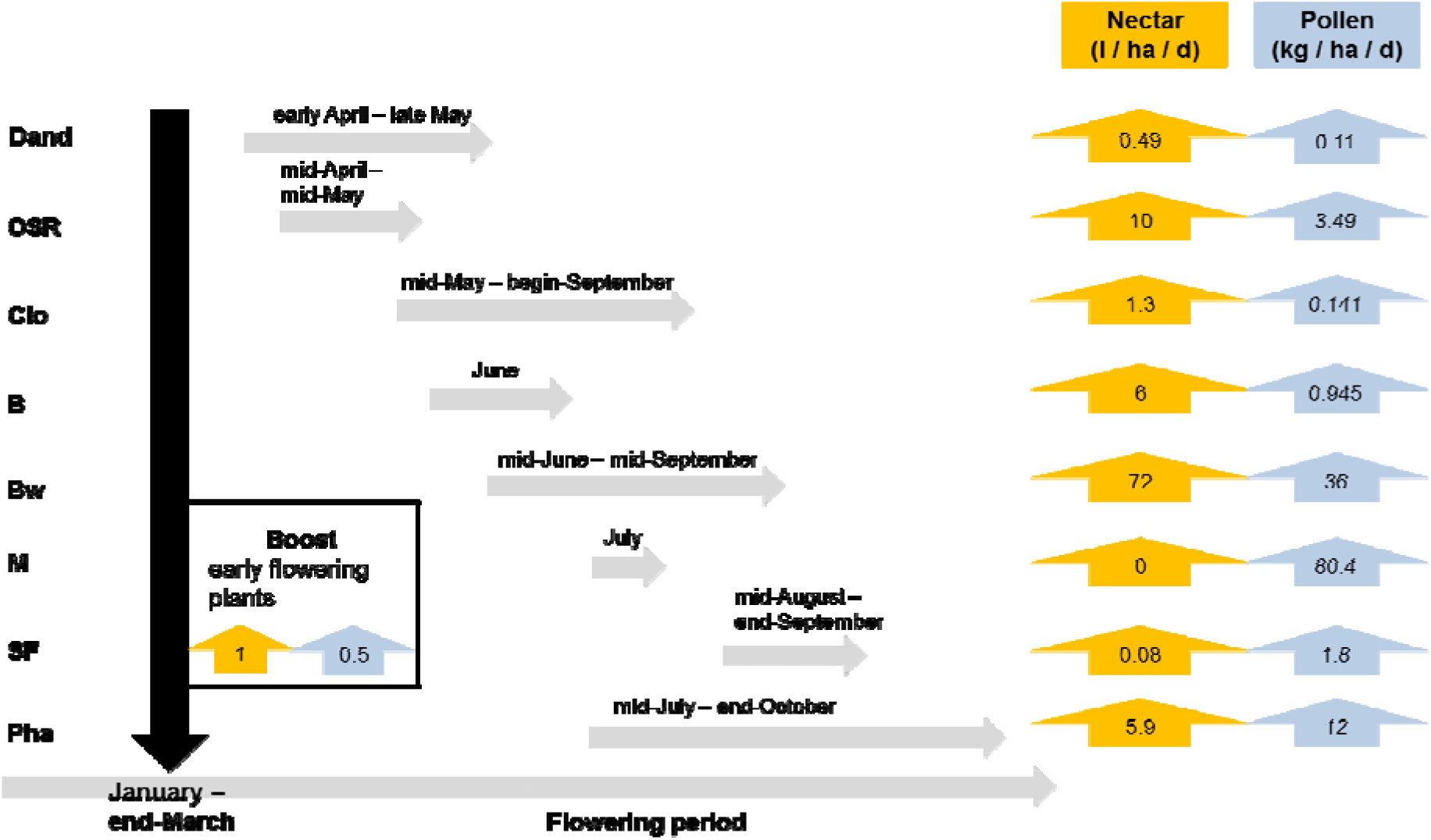
Seasonal flowering phenology of crop species and their average daily nectar and pollen amounts per hectare produced over their specific blooming period. Oilseed rape (OSR) provides on average 10 liter nectar and 3.49 kg pollen per day on one hectare over its blooming period from mid-April to mid-May.

**Table 3.** Nectar and pollen availability of crop species. : Flowering period of each crop species is given by its start and end date of blooming within a year. The amount of nectar and pollen per m^2^ of each crop species is calculated by its mean number of open flower units per m^2^ and its average daily amount of nectar and pollen produced per flower unit (i.e. single floret or flower head). Sugar concentration is given in mol/l. Abbreviations of each crop species is given (these are used in Table 4 and in the Figures 3 and 5). Literature data and equations for calculations are provided in the Supporting information S3 (Tables S6 and S7).

| Crop ID | Flowering Period (days) |  | Nectar (l / m<sup>2</sup>) | Pollen (g / m<sup>2</sup>) | Sugar concentration (mol / l) |
| --- | --- | --- | --- | --- | --- |
|  | Start | End |  |  |  |
| **Oilseed rape**<br>(*Brassica napus*) | 114 | 144 | 0.001 | 0.349 | 1.7 |
| **Maize**<br>(*Zea mays*) | 197 | 210 | 0 | 8.036 | 0 |
| **Sunflower**<br>(*Helianthus annuus*) | 237 | 264 | 0.000008 | 0.18 | 0.7 |
| **Field bean**<br>(*Vicia faba*) | 153 | 182 | 0.0006 | 0.0945 | 1.46 |
| **Cereals** | 1 | 365 | 0 | 0 | 0 |
| **Dandelion**<br>(*Taraxacum officinale*) | 92 | 169 | 0.000049 | 0.011 | 1.19 |
| **White clover**<br>(*Trifolium repens*) | 140 | 242 | 0.00013 | 0.0141 | 1.08 |
| **Rye-grass**<br>(*Lolium perenne*) | 1 | 365 | 0 | 0 | 0 |
| **Buckwheat**<br>(*Fagopyrum esculentum*) | 172 | 260 | 0.0072 | 3.6 | 1.17 |
| **Phacelia**<br>(*Phacelia tanacetifolia*) | 196 | 255 | 0.00059 | 1.2 | 1.05 |
|  | 265 | 301 | 0.000443 | 0.9 |  |

Farmlands do, of course, provide certain amounts of nectar and pollen in semi-natural habitats. Since very limited data are available on the amount and spatiotemporal distribution of these semi-natural resources (but see Baude et al. 2016 and Becher et al. 2018), we ignored them in our overall analyses. However, we estimated how much this simplification affects the results by an in-depth mechanistic analysis of two scenarios contrasting in crop diversity (a 4 crop and an 8 crop species combination both resulting in rapid colony extinction). As shown in Fig. 2, we tested different scenarios by introducing a semi-natural habitat corresponding to 1 ha in size. We varied the daily nectar amount from 1 up to 3 l and the daily pollen supply from 100 to 500 g in this patch. We applied two flight distances of 1000 m (intermediate distance as shown for hypothetical landscapes) and 2000 m (“stressful distance” as shown for artificial landscapes; see Tab. S15 and Figs. S18 - S19; also Horn et al. 2016). Nevertheless, our nectar amounts for the semi-natural habitat are in the range of the estimated yearly nectar productivity per hectare of land-use classes e.g. improved grassland or broad-leafed woodland reported by Baude et al (2016).

#### Crop diversity: number and relative abundance of crop species

We characterized crop diversity by the number of crop species and their relative abundance in the arable land. The number of crop species (hereafter called ‘*#Crops*’) considered were: one crop (monocultures), and combinations of three, four and eight crop species (Fig. 2).

The relative abundance of crop species (hereafter called ‘*abundance’*) refers to the proportion of the arable land in the farmland that is occupied by the crop species considered. For monocultures, *abundance* was set to 100 %. For three crop species, abundance of each crop was set to 33.3 %. For four and eight crop combinations, we contrasted *equal* and *dominant* distributions of abundances. For example in the case of four crop species, *equal* meant 25 % *abundance* for each crop species, whereas *dominant* meant that one crop species occupies most of farmland area, i.e. 85 %, and each of the other three 5 %. For *dominant* abundances, each of the crops considered got an equal turn in being the dominant species (Fig. 2).

Additionally, we ran simulations with a low-abundance treatment of 1 % or 2 % for buckwheat and/or phacelia in farmland instead of 5 %, because these minor and cover crops are more likely to be implemented at very low abundances at the landscape-scale in real agricultural systems. For such simulations, the abundance of the dominant crop species e.g. in a four crops combination was set to 89 % up to 93 % depending on the low-abundance treatment (Fig. 2).

### Landscape generator NePoFarm

The input file required by BEEHAVE comprises a list containing the respective data on patch type (i.e. *cropID*), size, location, detection probability, patch size-dependent nectar and pollen amounts and sugar concentrations for each of the farmland patches for each day of the year. To produce input files for different cropping systems in an automated way, we developed a landscape generator, referred to as NePoFarm, implemented in R version 3.1.2 (R Development Core Team 2014), to generate daily nectar and pollen availability lists throughout a year. Each crop species is assigned to available farmland patches of the semi-realistic landscape templates based on a probability related to its abundance, and the amount of nectar and pollen provided during its flowering period is then calculated based on the assigned patch size. NePoFarm, its documentation, the R-code and the landscape template are available in the Supplement (S4).

### Experimental design and analyses

We ran simulations for five years or until the colony became extinct (see below) and used 30 replicates to take into account the stochasticity represented in the model (Becher et al. 2014). Our preliminary sensitivity test indicated that 30 replicates per scenario were sufficient to reduce variation in the outcomes (Figs. S20 and S21) and did not increase the significance levels by increasing the number of simulations.

We explored colony viability for the combinations of farmland structure, crop species identity and crop diversity listed in Fig. 2. For each of the five simulation years for each cropping system scenario, we used the same input file generated by NePoFarm. Thus, we did not implement crop rotation, for several reasons: crop rotation in a certain farmland area usually only changes the spatial arrangement of certain crops but not their relative proportion and thus not necessarily overall forage availability; some forage crops of arable land, temporary and permanent grassland (trefoil-grass mixtures) are cultivated for several years. Finally, some crops e.g. maize and grasses are cultivated in repeated succession every year (3 to 5 years) for biomass production for usage in biogas plants (e.g. De Vliegher et al. 2012).

### Indices for quantifying colony viability

#### Colony viability

We quantified honeybee colony viability in terms of the survival probability as the percentage of surviving colonies within five simulation years (hereafter called ‘*SurvProb*’; defined as the percentage of replicates in which a colony survived 5 years) and in terms of the mean time to extinction (hereafter called ‘*TTE*’; this metric was used only when the colony survived in none of the 30 replicates). Colonies smaller than 4,000 adult worker bees on December 31, and colonies where the number of adult bees went down to zero within the year (e.g. due to depletion of honey stores), were considered extinct or unable to survive without intervention by beekeepers (Becher et al. 2014, Rumkee et al. 2015, Horn et al. 2016).

#### P_H_/P_P_: indices for comparing forage-induced stress at the colony level

To compare the effects of forage availability determined by the spatiotemporal nectar and pollen supply of the different scenarios at the colony level, we calculated new indices: *P_H_* (honey-to-worker-productivity), and *P_P_* (pollen-to-worker-productivity). These indices are a first step to quantify the response of honeybee colony development and long-term viability to the nectar and pollen availability due to the species composition of certain cropping systems. They evaluate how much nectar and pollen (kilograms) provided by a certain cropping system in a given landscape could be converted into the production and maintenance of the colony’s work force in terms of worker bees over five years or over the time to extinction (*TTE*, see above). Our indices include several parameters about daily pollen and nectar consumption of a single worker bee over its complete larval and adult life span. These are incorporated into the model to calculate the number of worker bees generated as hatched larvae to adults over *TTE* (*N*_Workers_). To account for different life spans of different bee categories (e.g. short living summer bees, long-living winter bees) we averaged the life span over all these bee categories according to Rortais et al. (2005). In these two indices, we ignored the production of queens and drones and their corresponding food requirements, as they do not directly contribute to the colony’s work force. All parameters, equations and cited literature for calculation of these indices are given in Table S13.

Then, to evaluate a farmland’s ability to continuously provide sufficient amount of nectar and pollen to meet the colony requirements, we compared these indices with empirical estimates of an average colony’s annual food requirements. According to the literature a viable honeybee colony produces 100,000 up to 200,000 workers per year and thus requires an annual supply of 48 - 80 kg honey and 20 - 55 kg pollen (e.g. Seeley 1985, 1995, Keller et al. 2005). We applied the lower bounds as the minimum threshold for colony viability according to Seeley (1995) for comparing *P_H_* and *P_P_* derived from our simulations with the empirical estimates of viable colonies. Accordingly, *P_H_*and *P_P_* values from our simulations have to exceed 48 kg honey and 20 kg pollen, respectively, per year, summed up to a minimum conversion of 240 kg honey and 100 kg pollen over the 5 years simulation time, to achieve long-term colony viability.

To qualitatively rank farmland factors by their importance for colony viability in terms of *TTE*, *N*_Workers_, *P*_P_ and *P*_H_, we applied a procedure mimicking a local sensitivity analysis by varying only one factor at a time while the others were fixed at their nominal values (Saltelli et al. 2000). Details about this procedure are given in the Supplement (S6).

### Effects of forage supply of different cropping systems on colony dynamics

Scenarios composed of 4 and 8 crop species, which showed contrasting impacts on colony viability and *P_H_* and *P_P_*, were selected to explore in more detail the effects of spatiotemporal forage supply on colony dynamics and resilience. We selected scenarios where (1) the colonies died quickly within the first year (hereafter referred to as ‘*bad forage supply’*) and (2) the colonies survived over the simulation time of five years (hereafter referred to as ‘*continuous forage supply’*). To understand how these scenarios affected colony dynamics, we analyzed the frequency, timing and duration of forage gaps, i.e. periods in which honeybees cannot find any nectar and pollen in the landscape, and plotted their temporal nectar and pollen availability. Additionally, we wanted to understand whether in these selected scenarios the honeybee colony is able to satisfy its forage demand to maintain, after a disturbance (in this case foraging gaps), the basic functionality of the colony in terms of colony’s structure and dynamics, i.e. the life stages and tasks. To this end, the simulations of these selected cropping system scenarios were used to retrieve data on the source and strength of brood mortality, force of worker bees, number of flight trips, and on the colony’s honey stores. Next, we used monthly averages of these variables to compare the selected scenarios.

Specifically, we focused, following Horn et al. (2016), on mortality effects caused by a lack of pollen (quantified by the proportion of larvae that died in a model step, a day, because of lack of protein from pollen, referred to as ‘% *larval losses: pollen lack’*), reduction in the queen’s egg-laying rate (daily number of eggs by which the potential egg-laying rate was reduced, referred to as ‘*# eggs not laid*’), the number of larvae and worker bees, which determines the brood to nurse bee ratio (daily number of all larval stages and all adult workers, referred to as ‘# *larvae*’ and ‘# *workers*’), *flight trips* (daily number of flight trips to search for forage resources and to forage for nectar and pollen), and stores (daily amount of honey stored in kg, referred to as ‘*honey* (kg)’).

## Results

### Indices for quantifying colony performance

#### Colony viability: TTE and SurvProb

In total we screened 500 cropping system scenarios resulting in 60 with *SurvProb* > 0, i.e. at least one colony out of the 30 replicates was still alive after 5 simulation years (Fig. 4). In total we ran 15,000 simulations, in which colonies survived in 10.3 % of all the scenarios with eight crop species and in 4.3 % in the scenarios with four crop species. For all monocultures, the colony went extinct within the first year (Table 4).

**Figure 4.**
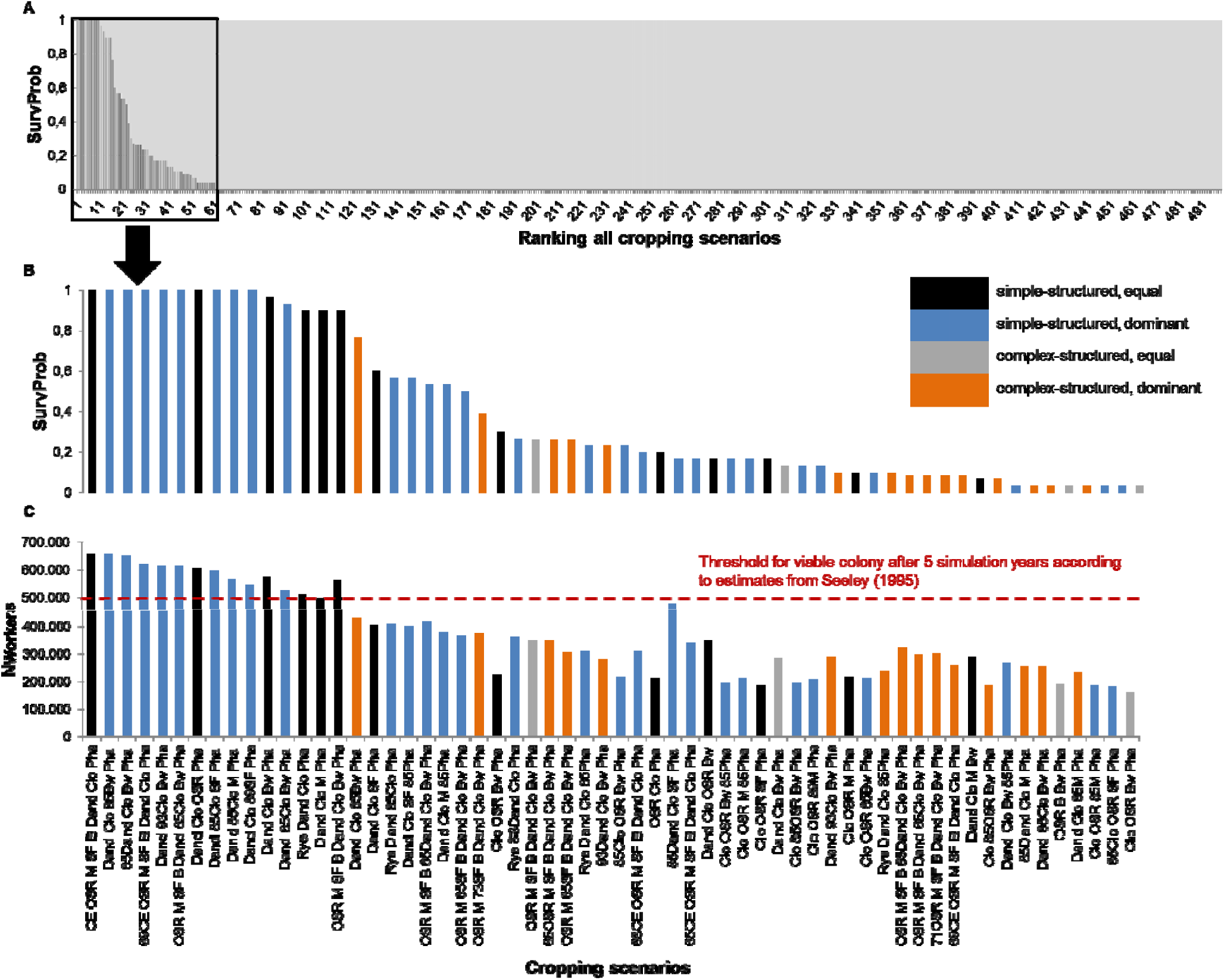
Ranking cropping system scenarios: (A) *SurvProb* (survival probability) of all screened scenarios over the five simulation years. (B) *SurvProb* of 60 out of 498 scenarios. The x-axis indicates the crop composition of the scenarios with colony in at least one, out of 30 replicates, surviving for five years. (C) *N_Workers_* (number of workers produced until the colony collapse or the end simulation time) for the scenarios with at least one surviving colony over the simulation time of five years. The threshold of produced workers for a viable colony after five years of simulation is marked by the red dotted line. Abbreviations of the crop species are given in Fig. 2.

**Table 4.**
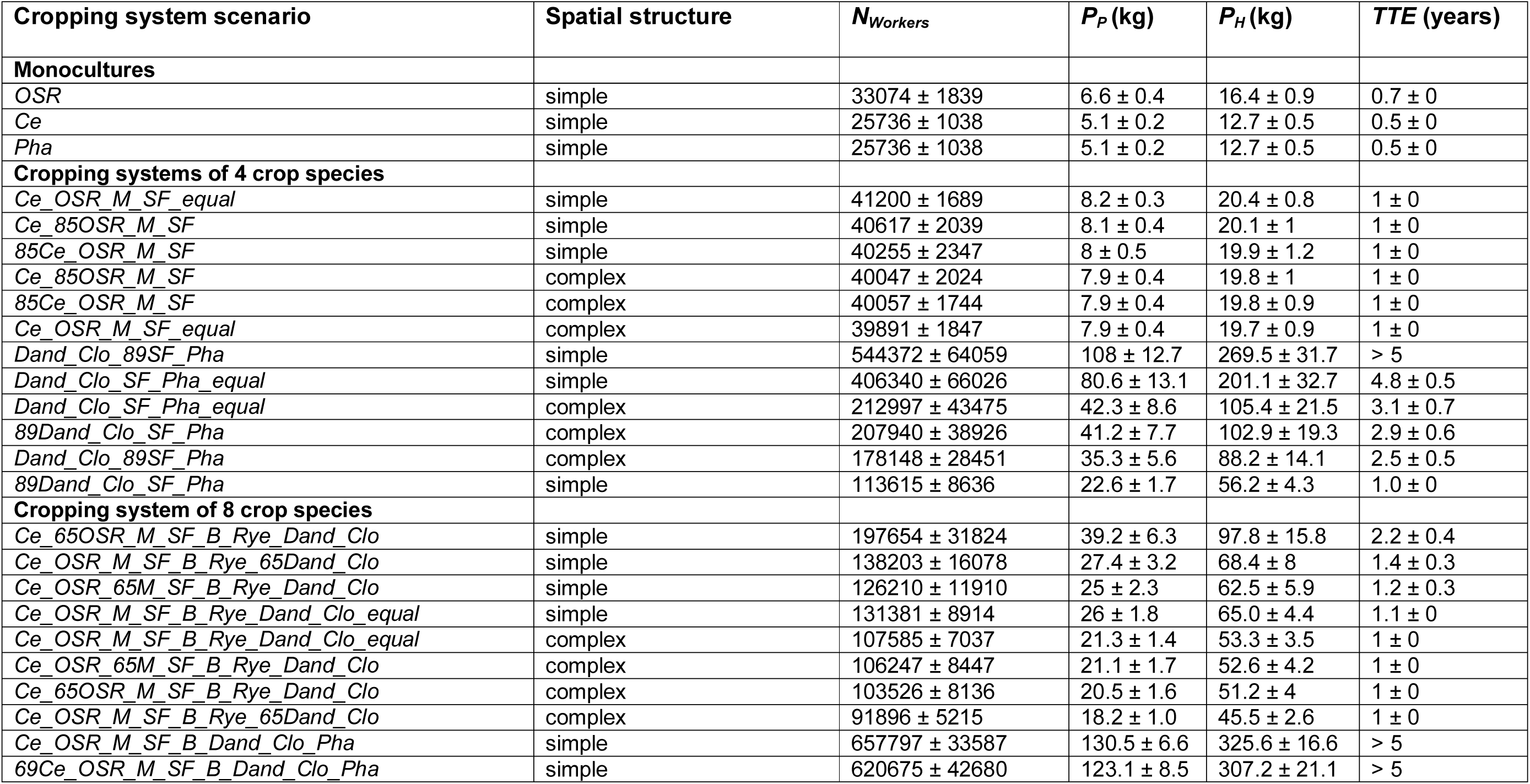
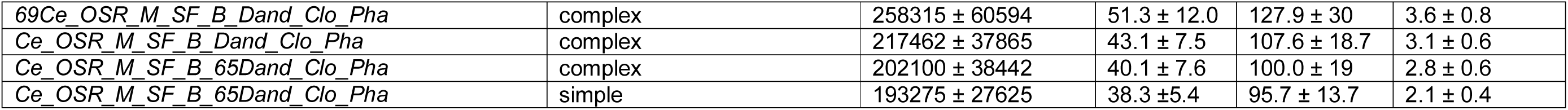
N_Workers_, *P_P_* and *P_H_* over 5 simulation years or until the colony became extinct, and *TTE* for selected cropping system scenarios. Abundances are assigned as follows: equal (all crops have the same abundance on total agricultural area (TAA)), 85 or 89 (in a 4 crop composition the dominant crop occupies 85% or 89% of TAA), 65 or 69 (in an 8 crop composition the dominant crop occupies 65% or 69% of TAA).

For only ten scenarios *SurvProb* was equal to 1: seven of those scenarios contained four crop species and the other three eight. For scenarios composed of three crop species only one out of 26 scenarios achieved *SurvProb* equal to 0.2. For 4 or 8 crop species scenarios, *TTE* ranged from 0.6 to 5.0 or was larger than 5 years (all replicated colonies survived at least the 5 years simulation time) as shown in Table 4.

#### P_H_/P_P_ indices

*N_Workers_*, the adult worker population produced by the colony over the entire simulation time (five years) or until extinction, which is included in the indices *P_P_* and *P_H_*, was 39,249 ± 2,423 for monocultures, 210,601 ± 48,976 for three crop combinations, varied between 23,053 ± 871 and 656,131 ± 65,879 for four crop combinations (depending on the different equality/dominance scenarios) and between 22,990 ± 860 and 657,797 ± 33,587 for eight crop combinations (mean ± sd; n = 30 per scenario). Only for scenarios with *SurvProb* > 0.9 did *N_Workers_*exceed the threshold of 500,000 adult workers bees (100,000 bees per year summed up over 5 years) over the entire simulation time, reflecting a viable colony according to empirical estimates by Seeley (1995) as shown in Fig. 4.

Fig. 5 shows the achieved *P*_H_ and *P*_P_ values for all scenarios and compares them with minimum thresholds of a viable honeybee colony derived from empirical estimates. For all monocultures, the colony converted up to 19.4 kg ± 1.2 kg honey and up to 6.6 kg ± 0.4 kg pollen (mean ± sd; n = 30 replicates), which are below the minimum threshold and the colony became extinct within the first year (Table 4). For the three crop system scenarios *P*_H_ and *P*_P_ ranged from 13.1 ± 0.7 kg to 104.2 ± 24.2 kg and 5.2 ± 0.3 to 41.8 ± 9.7 kg. For the four crop systems, converted honey ranged from 11.4 kg ± 0.4 to 324.8 kg ± 32.6 kg and pollen ranged from 4.6 kg ± 0.2 to 130.2 kg ± 13.1 kg, while for eight crops from 4.6 ± 0.2 kg and 11.4 ± 0.4 kg up to 130.5 ± 6.7 kg and 325.6 ± 16.6 kg, respectively. For some scenarios containing four or eight crop species PH and PP values slightly exceeded the minimum thresholds in the first, up to the third, year (Figs 5A, B), but colonies were then not able to convert sufficient nectar and pollen into a viable adult worker population thereafter and became extinct within spring or summer of the second or third year (SurvProb = 0).

**Figure 5.**
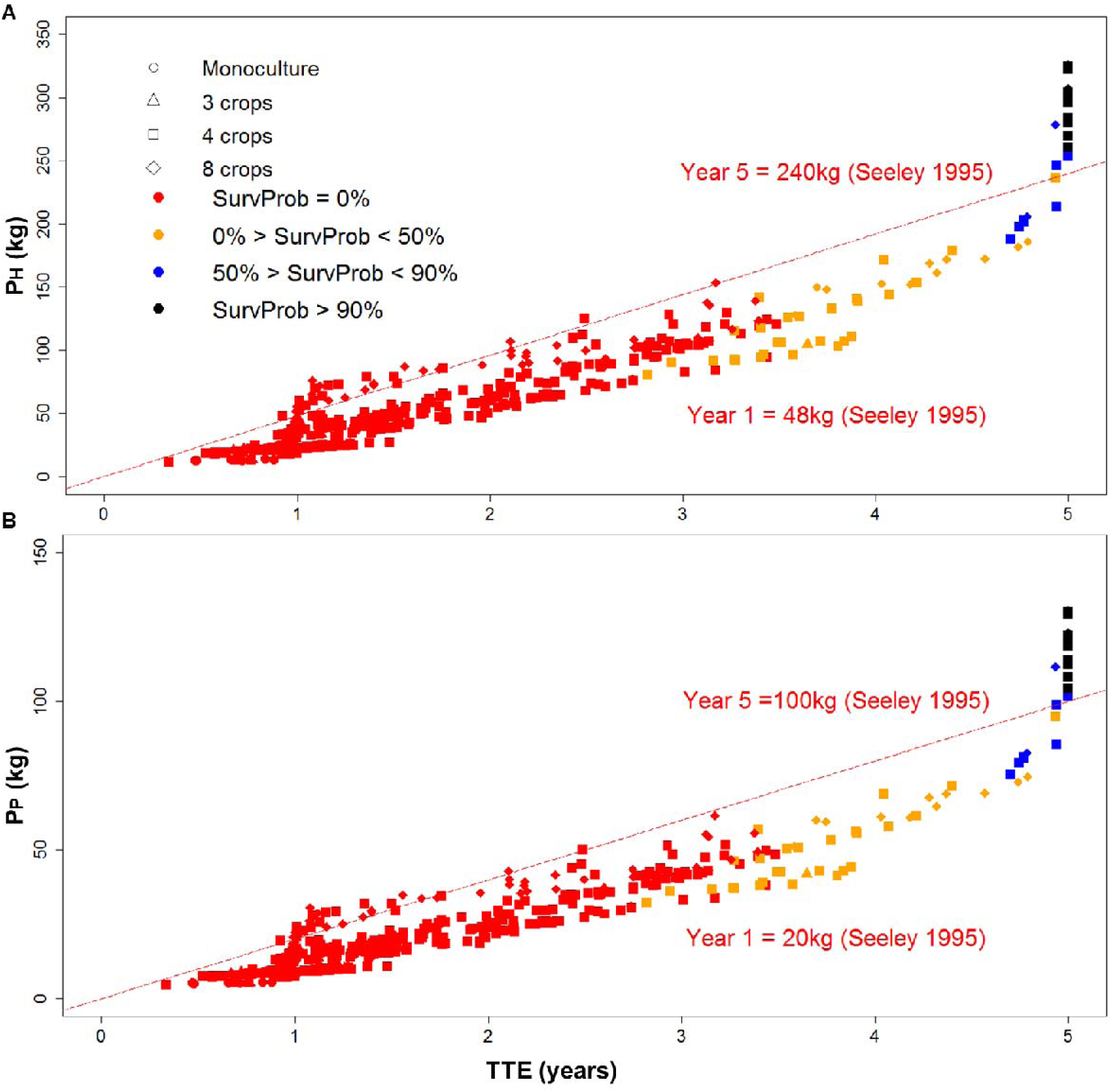
(A) P_H_ and (B) P_P,_ the amount of honey and pollen that the colony was able to convert into production and maintenance of an adult worker bee population in a given agricultural landscape, over simulation time of 5 years or until the time to extinction for all scenarios are plotted. Red dashed lines illustrate empirical estimates of the minimum colony requirements of honey and pollen per year to produce a viable worker population according to Seeley (1995) (at least 48 kg honey and 20 kg pollen per year to produce a sufficient force of 100,000 up to 200,000 worker bees per year). To survive over 5 years simulation time a viable colony needs to convert at least 240 kg honey and 100 kg pollen. *#Crops* is illustrated by symbols and *SurvProb* by colour palette.

Considering all simulations, increasing *#Crops* led to an increase in *TTE* and *P_H_*, while differences in *TTE* between both farmland structures were small (Fig. 6). Our sensitivity analysis indicates that *#Crops* explains most of the variation in *TTE* and P*_H_* (R_adjusted_^2^: 0.119 and 0.3379, respectively), followed by *cropID* (R_adjusted_^2^: 0.1076 and 0.1363), *abundance* (R_adjusted_^2^: 0.02078 and 0.02349), and *farmland structure* (R_adjusted_^2^: 0.003716 and 0.009257). Successful cropping system scenarios with SurvProb > 0 and highest *TTE*, *P_P_* and *P_H_* frequently included trefoil-grass pastures, phacelia, rye-grass pastures infested by dandelion, oilseed rape and buckwheat as crops (Table 4). Moreover, 68.3 % of scenarios implemented in simple-structured landscapes with few, big patches achieved *SurvProb* > 0, whereas only 31.7 % of scenarios implemented in landscapes with many small patches resulted in colony survival (Table 4).

**Figure 6.**
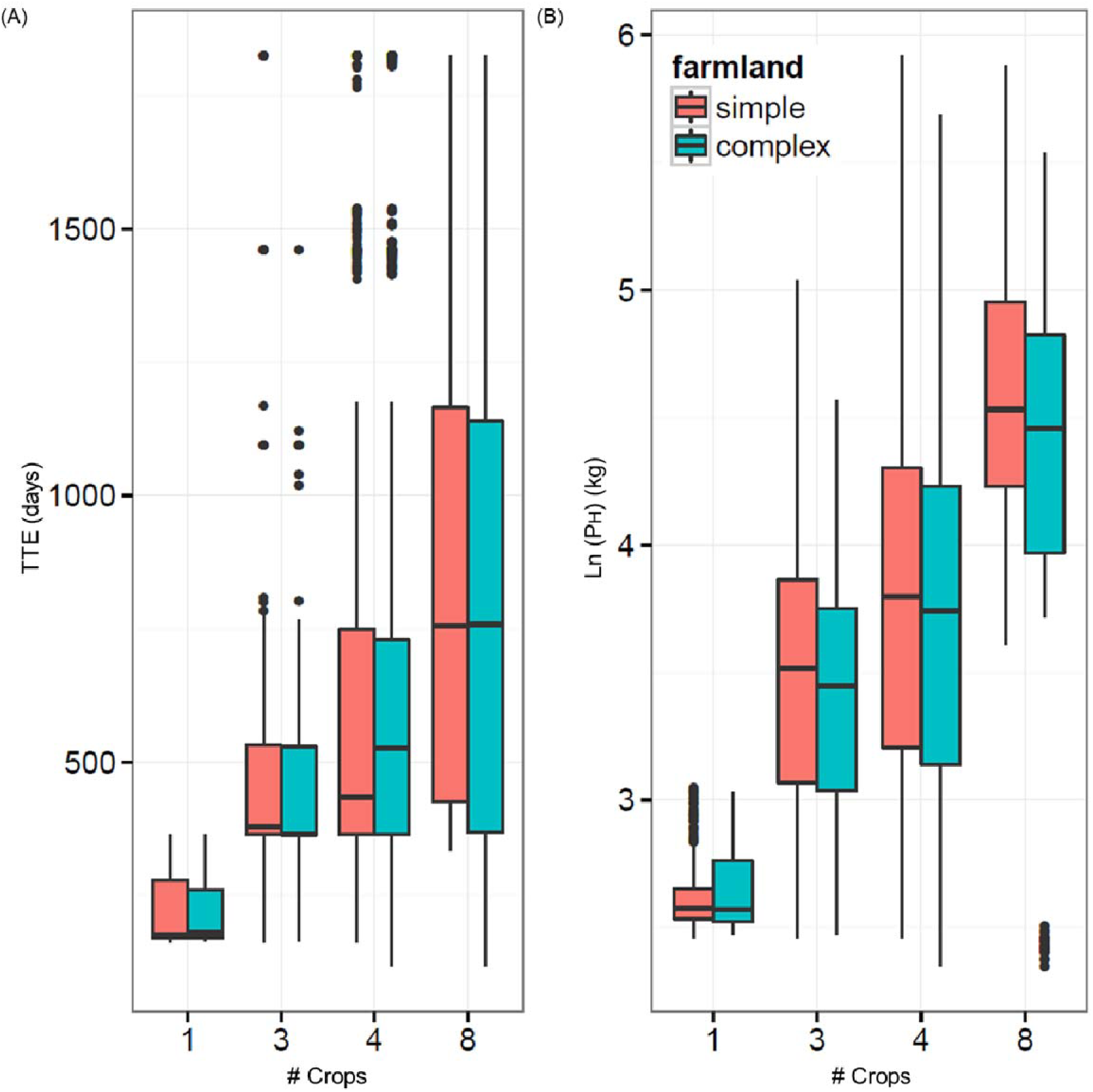
Relation between increasing *#Crops* and (A) *TTE* (days) and (B) *P_H_* (log-transformed, kg) for both farmland structures (simple-structured with few large fields and complex-structured landscape with many small-sized fields).

### Effects of forage supply on colony dynamics

#### Cropping system scenarios composed of 4 crop species

*4 crop species - discontinuous forage supply* (Fig. 7 and Fig. 9): For scenarios involving the four crop species cereals, oilseed rape, maize and sunflower, for all combinations of abundances and farmland structures, *P_H_*, *P_P_* and *N_Workers_* were below the annual minimum threshold of a viable honeybee colony. Thus, the colony became extinct within the first year (Table 4). As illustrated in Fig. 7A, after the early spring flowering there was a gap in nectar and pollen supply (hereafter referred to as ‘forage gap’) of three weeks from end-March onwards before oilseed rape began to bloom in late April. Between mass-flowering of oilseed rape till the 24^th^ May and the start of maize flowering from mid-July to end-July, there is a second and very long forage gap of two months. Then, after the two weeks of pollen provision by maize, from end-July onwards no nectar and pollen was available for one month before sunflower began blooming in late August (third forage gap). From 22^nd^ September onwards, nectar and pollen is no longer provided as sunflower ceased blooming (fourth late-seasonal forage gap).

**Figure 7.**
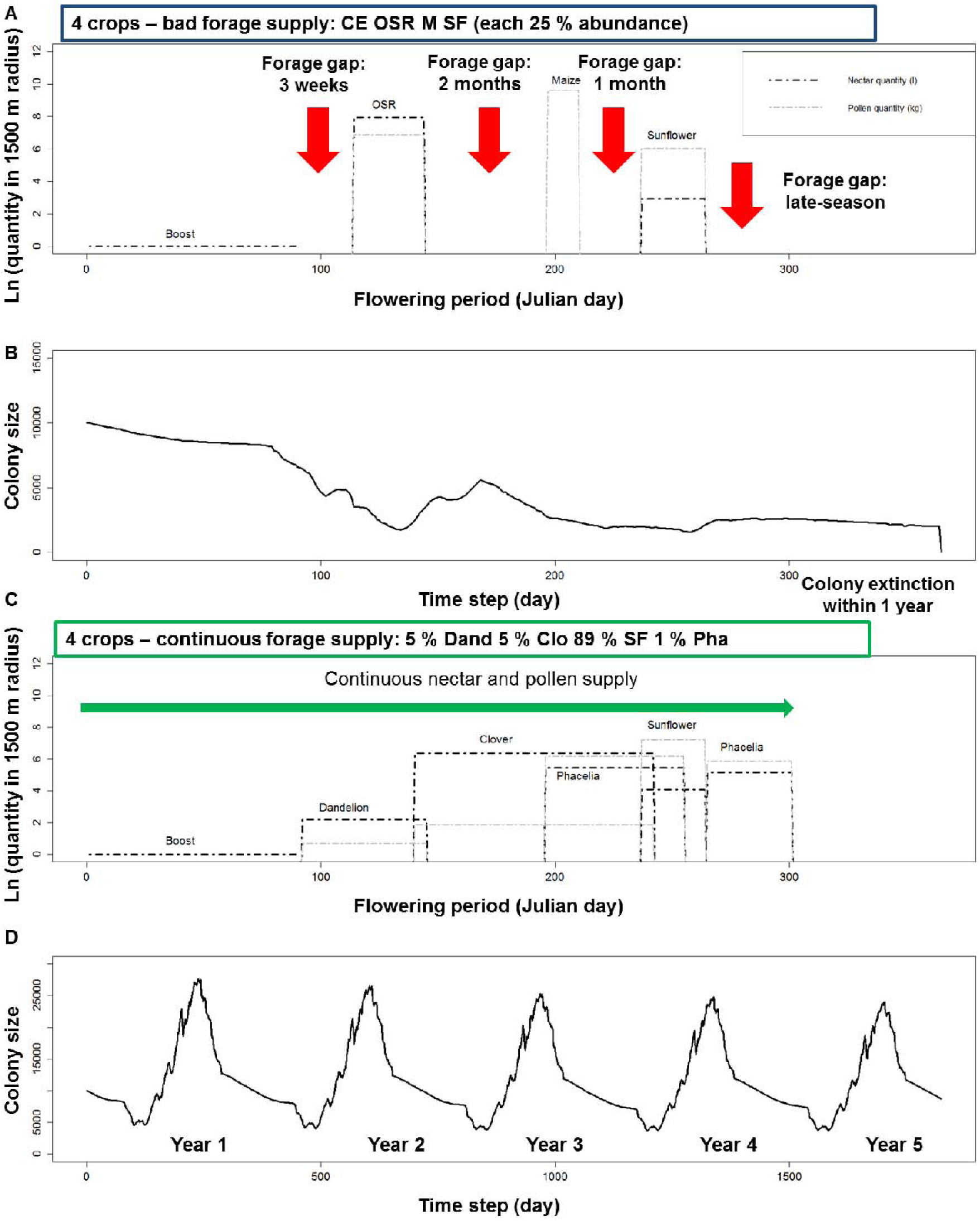
For cropping systems of four crop species shown are: (A, C) Seasonal nectar and pollen availability within 1500 m foraging distance (both log-transformed; nectar is given in l (black dotted lines; pollen is given in kg (grey dotted line)) and (B, D) honeybee colony size (number of workers) development over time on a daily basis. Frequency, timing and duration of temporal gaps in nectar and pollen supply (forage gaps) are indicated by red arrows, whereas green arrows indicate continuous nectar and pollen supply over time without forage gaps. (A) and (B) demonstrate the results for a bad forage supply and (C) and (D) for continuous forage supply.

**Figure 8.**
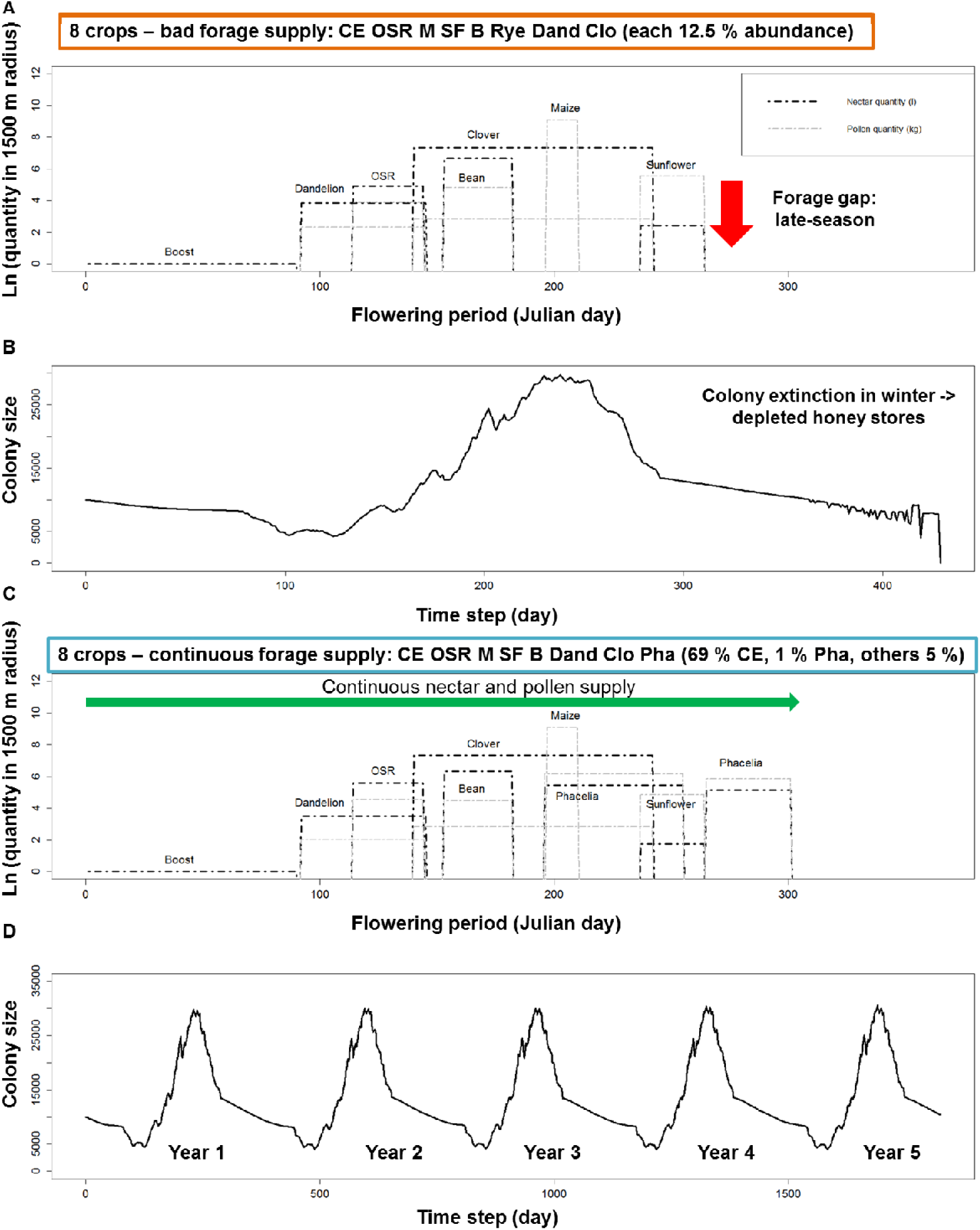
For cropping systems of eight crop species shown are: (A, C) Seasonal nectar and pollen availability within 1500 m foraging distance (both log-transformed; nectar is given in l (black dotted lines; pollen is given in kg (grey dotted line)) and (B, D) honeybee colony size (number of workers) development over time on a daily basis. Frequency, timing and duration of temporal gaps in nectar and pollen supply (forage gaps) are indicated by red arrows, whereas green arrows indicate continuous nectar and pollen supply over time without forage gaps. (A) and (B) demonstrate the results for a bad forage supply and (C) and (D) for continuous forage supply.

**Figure 9.**
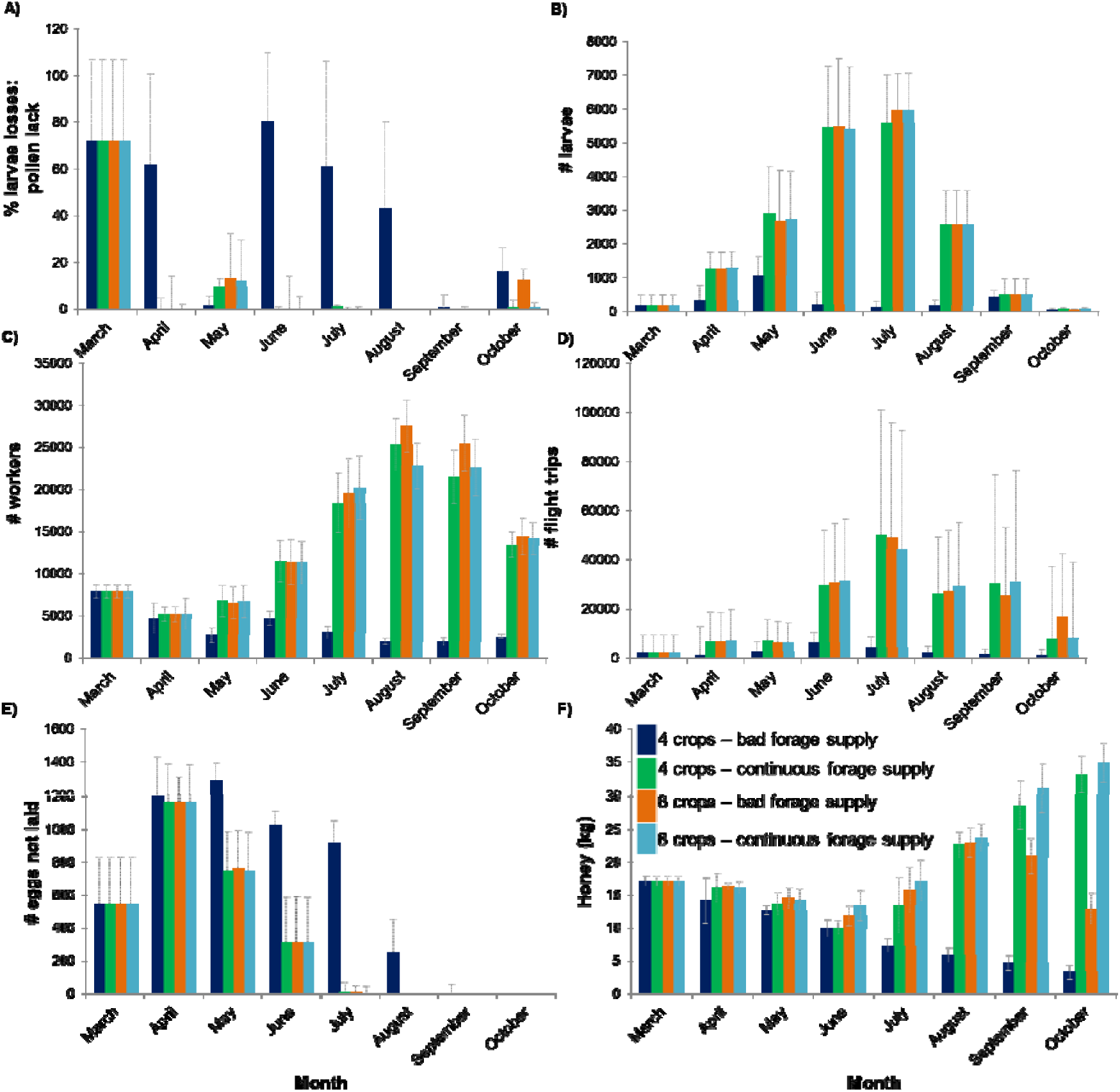
Selected cropping system scenarios composed of 4 and 8 crop species (simple-structured) - coloured bars match scenarios illustrated in Figure 7 and 8: Effects of frequency, timing and duration of temporary gaps in nectar and pollen supply (‘forage gaps’) on colony’s underlying mechanisms are shown: monthly averages of (A) mortalities of larvae caused by a lack of pollen (proportion of larvae that died in a day due to lack of protein from pollen; ‘% larval losses: pollen lack’), (B) the total number of larvae and (C) worker bees (daily number of larval stages and adult workers; ‘# larvae’ and ‘# workers’), (D) flight trips (daily number of searching and foraging trips), (E) reduction in the queen’s egg-laying rate (daily number of eggs by which the potential egg-laying rate was reduced; ‘# eggs not laid’), and (F) colony’s honey stores (daily amount of honey stored in kg; ‘Honey (kg)’). Means and standard errors are shown for 30 replicates per scenario.

During the first forage gap the colony faced high daily larval losses in April up to 61.8% due to lack of protein, which led to reduced daily production of later larval stages, and workers in terms of in-hive bees, and foragers in May (Fig. 9A-C). For the mass-flowering period of oilseed rape in May, plenty of nectar and pollen was available; thus the daily larval losses due to protein lack were only 1.3% (Fig. 9A). During the second forage gap, after six days without pollen income, the pollen stores were depleted on the 1^st^ June. Although no nectar and pollen was available in the landscape, the small number of worker bees performed on average 6273 ± 4148 and 4287 ± 4549 flight trips per day in June and July to search for nectar and pollen resources (Fig. 9D), leading to increased forager mortality. Simultaneously, average daily larval losses were 80.5% in June and 61.2% in July, the reduction in the egg-laying rate of the queen was high, and few larvae survived (Fig. 9A, B, E). During the third forage gap, the colony is faced with high larval losses (43.3% ± 36.8%) resulting in low numbers of larvae, a very small peak adult worker population, a reduction in the queen’s egg-laying rate, and low honey stores (Figs 7B; 9A-F). After the third forage gap, sunflower fields provided 18.7 l nectar and 420 kg pollen per day within 1,500 m foraging radius (233.25 ha), but the reduced worker force was unable to perform sufficient foraging trips, and honey stores were almost depleted (4.6 kg ± 1.1) (Fig. 9C, D, F). Consequently, the colony size and its honey stores kept decreasing (Fig. 7B, Fig. 9C, F). Taken together, colony size rapidly decreased during the year until the colony became extinct at the end of the first year (Fig. 7B).

For the scenarios in which we represented floral resources in semi-natural areas, colony survival strongly depended on the distance of that semi-natural habitat to the hive, which represents the energetic efficiency of exploiting it. If such floral resources provided at least 3 l nectar and 300 g pollen per day throughout the year were located 1,000 m away from the hive, up to 62 % of the colonies were able to survive for five years, but N*_Workers_* and P*_H_* and P*_P_* were below the viability thresholds. Increasing the distance of the additional semi-natural floral resources to 2,000 m led to colony extinction in the second year at the latest (Fig. 10).

**Figure 10.**
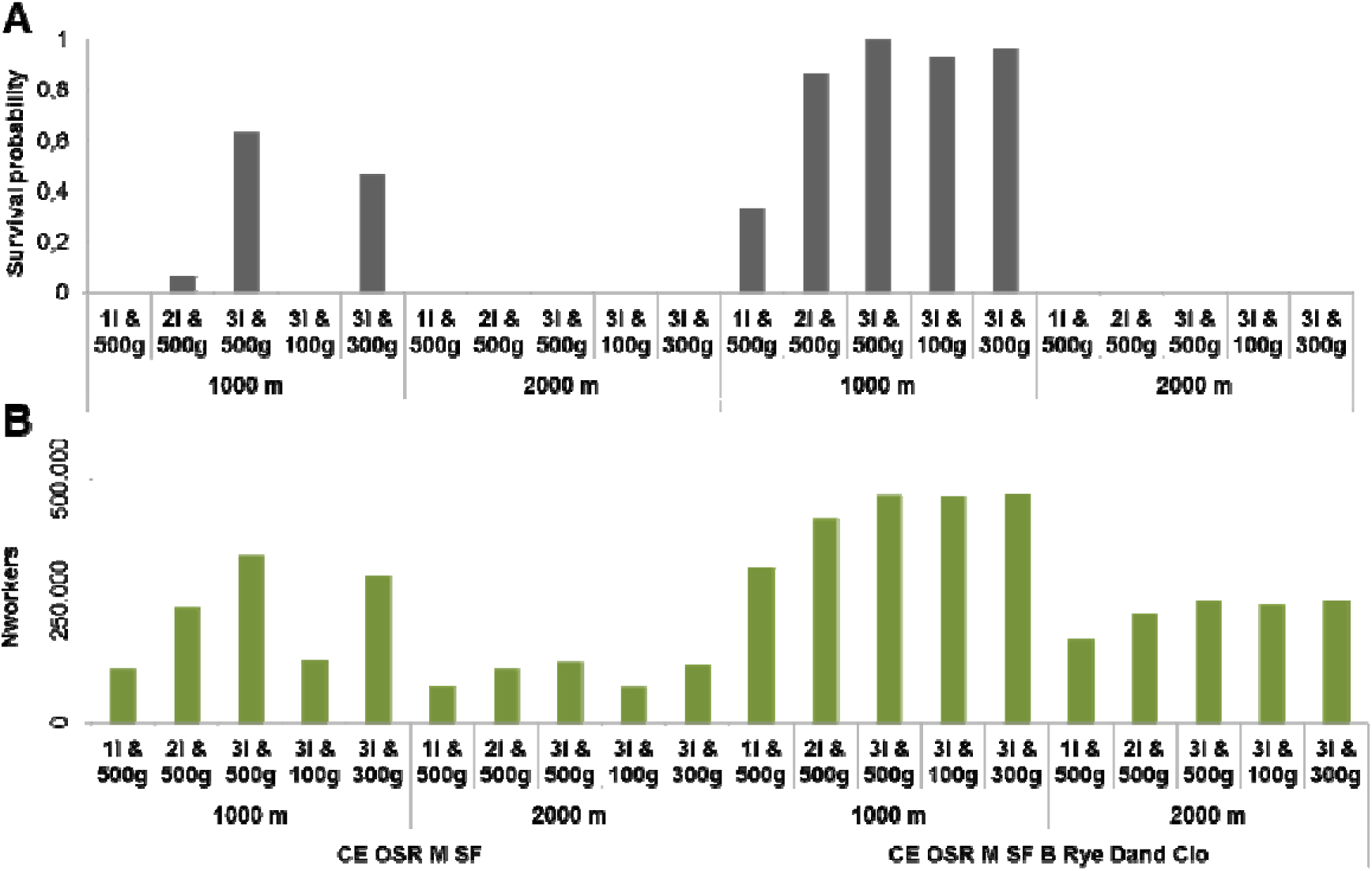
Effects of semi-natural habitats for two exemplary cropping systems of low diversity (4 crops – CE OSR M SF) and high diversity (8 crops – CE OSR M SF B Rye Dand Clo) on (A) survival probability after five simulation years, and (B) colony productivity measured as the number of worker bees produced over five years or until the colony became extinct (N_Workers_). Semi-natural habitats are represented as single forage patches of 1 ha size located 1000 m or 2000 m away from the honeybee hive and providing constant nectar and pollen rewards over the whole year. The colonies go extinct within one (low diversity example) or two years (high diversity example).

*4 crop species – continuous forage supply* (marked by green bars in Figs 7 and 9): Scenarios composed of the four crop species: sunflower, rye-grass pasture infested by dandelion, trefoil-grass pasture and phacelia provided continuous nectar and pollen supply until end-October (Fig. 7C). For abundances of 89 % sunflower, 5 % of each rye-grass pasture infested by dandelion and trefoil-grass pasture, and 1 % phacelia, highest values for *P_H_* and *P_P_* exceeding minimum thresholds were achieved: on average 269.5 kg ± 31.7 kg honey and 108 kg ± 12.7 kg pollen were converted into 544,372 ± 64,059 worker bees over the five years simulation time (Fig. 4 and Fig. 5; Table 4). Here, the abundance of phacelia on the total agricultural area is small, but the patch providing late-seasonal nectar and pollen supply was placed, by chance, directly adjacent to the hive (0.1 m distance). In scenarios where rye-grass dandelion or trefoil-grass pastures dominated this cropping system (89%), the few phacelia patches were, by chance, much further away from the hive (> 1805.6 m), resulting in much lower values for *P_H_*, *P_P_* and *N*_Workers_ (Table 4). Continuous forage supply, and especially high abundance of nectar and pollen from end-August until late October close to the hive (89% sunflower; phacelia: adjacent to the hive) as shown in Fig. 7C led to low larval losses from April to August, high numbers of larval stages, a high number of worker bees, many flight trips to forage nectar and pollen, low reduction in the queen’s egg-laying rate, and a high storage of honey in October (Fig. 9A-F).

#### Cropping system scenarios composed of 8 crop species

*8 crop species – bad forage supply* (marked by orange bars in Fig. 4 and 9): Cropping systems composed of the eight crop species cereals, oilseed rape, maize, sunflower, bean, rye-grass pasture, rye-grass pasture infested by dandelion and trefoil-grass pasture (pasture sown with mixture of rye-grass and white clover) resulted in *P_H_*and *P_P_* up to 97.8 kg ± 15.8 kg honey and 39.2 kg ± 6.3 kg pollen converted into 197,654 ± 31,824 workers until the colony went extinct within the second year (Table 4). Independent of abundance, dominant crop species and farmland structure in all those scenarios, the cropping system was not able to continuously provide sufficient nectar and pollen to ensure colony viability over the annual cycle (Figs 4A, B). The cropping system provided continuous forage supply from early spring to mid-September, but after sunflower ceased blooming, there was a late-seasonal forage gap from mid-September onwards (Fig. 8). Thus, from mid-September onwards, there were many searching trips in late September and October, but due to the gap in nectar supply, the honey stores decreased to 12.9 kg honey in October (Fig. 9D, F).

Again, we ran simulations with additional floral resources of semi-natural habitats. If this additional food source provides at least 2 l nectar and 300 g pollen per day, and is located at either 1,000 m and 2,000 m distance, it is sufficient to compensate for the late-seasonal gap and to ensure viability of the modelled honeybee colony. However, colony size and colony honey stores on 31^st^ December were low (1000 m: 10,841 workers and 18.9 kg honey at the end of the first year; 5,816 workers and 10.3 kg honey after five years), indicating these colonies would not cope well with further stressors such as diseases, pesticide effects, or bad weather.

*8 crop species – continuous forage supply* (Fig. 8 and Fig. 9): For scenarios including cereals, oilseed rape, maize, sunflower, bean, rye-grass pasture infested by dandelion, trefoil-grass pasture and phacelia as crop species providing continuous forage supply, 107.6 kg - 325.6 kg honey and 38.3 kg - 130.5 kg pollen were converted into the production and maintenance of 217,462 - 657,797 worker bees (Table 4), which covers the range from poor to good colony performance and viability. The cropping systems occurring in complex-structured agricultural landscapes with 12.5% abundance of all those crop species did not allow the colony to convert sufficient honey and pollen to produce a sufficient worker population over time. Moreover, if rye-grass pasture infested by dandelion dominated (65%) and phacelia abundance was 5%, the colonies were unable to continuously derive enough nectar and pollen from the agricultural systems to produce enough workers, and consequently became extinct within three years in both simple- and complex-structured landscapes (Table 4). In these cases, late-season forage-providing phacelia patches were located between 1347.2 m and 3545.7 m from the hive and did not enable colonies to remain viable. On the other hand, even for scenarios where cereals, which do not provide forage at all, dominated the cropping system (69% abundance) and phacelia occurred at very small abundance (1%) in a simple-structured landscape, the colony converted 307.2 kg ± 21.1 kg honey and 123.1 kg ± 8.5 kg pollen into the production of 620,675 ± 42,680 worker bees within five years (Table 4; Fig. 4), because late-flowering phacelia was located adjacent to the hive and provided plenty of nectar and pollen till late-October (Fig. 8C). Here again, good colony performance was reflected in low larval losses, as pollen was not lacking between April and August, high numbers of larvae and workers, many foraging trips, low reduction in the queen’s egg-laying rate, and high storage of honey in October (Fig. 9A-F).

## Discussion

Forage availability in agricultural landscapes is not just a matter of how much nectar and pollen there is but also of where the forage resources are and how their phenology ensures temporal continuity in nectar and pollen supply. We therefore presented a modelling framework to analyses spatiotemporal nectar and pollen supply maps of different cropping system scenarios generated by NePoFarm with the existing model BEEHAVE.

Our results demonstrate that number of crop species and their identity determine the frequency, timing and duration of temporary gaps in nectar and pollen supply (“forage gaps”) and thereby strongly affect colony performance. Frequent and prolonged forage gaps and in particular late-seasonal forage gaps lasting several weeks showed detrimental impacts on colony viability via cascading effects on colony’s life stages and tasks. In contrast, relative crop abundance and farmland structure were less important.

As a metric for colony performance in a given landscape and cropping system, we used *SurvProb* and in particular the conversion of honey and pollen into the production and maintenance of a viable adult worker population – *P*_H_, *P*_P_ and N_Workers_. We compared these metrics with empirical estimates of colony’s honey and pollen requirements as reported by Seeley (1985, 1995; see also Keller et al. 2005), which were used to estimate the minimum number of worker bees that are needed to be produced over the year to ensure colony survival. The minimum values for *P*_H,_ *P*_P_ and N_Workers_ for viability obtained in our simulations are in good agreement with the values estimated by Seeley (1995) as shown in Table 4 and Figures 4-5. It should be noted that we achieved this agreement without any calibration. We take this as a strong indicator that despite the uncertainty in parameters for BEEHAVE, the modelled honeybee colony dynamics are likely to be structurally realistic enough and offers the opportunity to evaluate cropping regimes and mitigation measurements. Still, validating our quantification of the nectar and pollen provided by a given agricultural landscape requires data and experiments which are not available. Like with models used for Population Viability Analysis in conservation biology, our model predictions are thus relative, not absolute. Assuming that potential biases in our parameterization have similar effects on cropping systems compared, we still can rank those systems in terms of their suitability for honeybees.

### Crop species number and identity determine temporary gaps in forage supply

Our in-depth analyses of scenarios with 4 and 8 crop species of low versus high viability showed that there is no simple and general relationship between crop diversity and honeybee viability. Rather, crop composition and in particular the presence and spatial arrangement (see next paragraph) of particularly rewarding floral resources such as phacelia can be decisive. The reason for the strong impact of number and identity of crop species is that they determine the temporal continuity of nectar and pollen supply and thus the frequency, timing and duration of forage gaps during the colony’s development cycle.

For monocultures, before and after a short-term period of massive nectar and pollen supply, long-lasting forage gaps led to rapid colony extinction within one year (Table 4). Similarly, colonies located in cropping systems with 3 crops were deemed not viable. For some cropping systems with 4 crops viability occurred, but was still low. In particular for systems composed of the 4 crop species cereals, oilseed rape, maize and sunflower; which are typical crops in European agriculture (Table S4, Figs S2-S7), colony survival was not possible (Table 4) because these resulted in four forage gaps, over several weeks to two months, in April, June to July, August, and end-September (Fig. 7A). These forage gaps have detrimental cascading effects on the colony and lead to its rapid extinction within one year. The forage gaps under this cropping system hit the colony during sensitive phases. In BEEHAVE, the maximum potential egg-laying rate of the queen is from mid-June to mid-July. Thus, the number of newly emerging brood stages developing into new generations of adult worker bees for nursing and foraging tasks is highest in July and August, when the colony is achieving its peak colony size. However, sufficiently high numbers of worker bees over the full annual cycle are necessary to ensure sufficient pollen stores to feed the larvae with protein-rich jelly. This is required to achieve sufficiently high peak colony sizes and thereby ensure sufficient honey stores to survive over winter (Horn et al. 2016). In BEEHAVE, in accordance with field observations (Blaschon et al. 1999), foragers try to maintain pollen stores that last for about seven days. After depletion of pollen stores the larval stages start to die due to decreasing protein content of the jelly. Consequently, flight activity of foragers searching for urgently needed nectar and pollen resources in the surrounding landscape increases. The first gap of three weeks in April (Fig. 7A) caused high larval mortality due to pollen shortage resulting in lower larval numbers and, consequently, reduced numbers of worker bees in May (Fig. 9). This weakens the colony already in spring. The work force of worker bees for nursing and foraging is thus already reduced before the next prolonged forage gap follows from June to mid-July (Fig. 7A). Here again, after the pollen stores become depleted, the ratio of foragers to in-hive bees and flight activity increased. However, as no pollen and nectar is offered within this gap period and as the colony’s brood and work force have already been weakened in the April gap, mortality of larvae due to lack of pollen was high and the queen’s egg-laying rate due to lack in available nurse bees was markedly reduced. These losses in this sensitive phase of colony development reduced the number of newly emerging worker bees for foraging and nursing tasks (Fig. 9). The third gap in August (Fig. 7A) worsened the situation. Due to reduced numbers of workers, few foraging trips are undertaken especially in August and September. Thus, honey stores are already very low in August and then the late-seasonal forage gap (Fig. 7A) deteriorates the situation further (Fig. 9D, F). As a result, honey stores are not sufficient for overwintering and the colony collapses within the first winter.

As our modelled cropping systems were simplified in terms of abundance of floral resources of semi-natural habitats, we conducted additional simulations with representations of additional floral resources which provide continuous but low amounts of nectar and pollen. Whether or not these additional resources are able to buffer the negative effects of forage gaps depends on how energetically efficient it is for the bees to use them. If costs for exploiting these additional resources are low, because semi-natural floral resources are nearby, viability could be increased, although not necessarily to the same level as for cropping systems without forage gaps. The effect of semi-natural resources is similar to the effect of small but highly rewarding and well-timed patches of flowering phacelia, which are only efficient if close enough to the hive. These results indicate that semi-natural habitats buffer the effects of forage gaps in agricultural systems, but whilst data remains sparse on the quality of resources provided and how efficiently they can be exploited by the bees, it remains unknown as to how well they will buffer the effects of forage gaps in real agricultural landscapes with low crop diversity.

Modelled farmland systems composed of 4 or 8 crop species that covered early-, mid- and late-seasonal flowering such as oilseed rape or dandelion, and clover or sunflower, and buckwheat or phacelia, even if their overall abundance was low and other common crops such as cereals that do not provide any nectar and pollen dominate the cropping system, were able to offer continuous forage supply (Table 4). These cropping systems therefore ensure forage availability especially during sensitive phases of colony development and ensure long-term colony viability (Fig. 8). Continuous nectar and pollen supply without temporary gaps can also be achieved by other flowering crop species than those presented in this study. However, in real landscapes it might not be sufficient to rely on a limited number of crop species, particularly to ensure pollen nutrition. Pollen varies in terms of availability and concentration of necessary amino acids among plant species, but BEEHAVE neglects such differences in pollen qualities. Dandelion, for example, is important for honeybees in early spring and during times of dearth, but its pollen is lacking in some necessary amino acids resulting in lowered brood rearing success when honeybees rely on dandelion alone (Loper & Cohen 1987). If dandelion (infesting rye-grass pasture) is the only forage resource in spring, the scope of the model to capture all effects of this reduced pollen quality on brood rearing is limited; which might have led to an overestimation of honeybee colony performance. However, bees tend to collect abundant pollen irrespective of its nutritional value in times when high-quality pollen is rarely available (Höcherl et al. 2012).

Our interpretation of our results might in principle be limited by having ignored the sampling effect (De Laender et al. 2016): higher diversity scenarios have a higher probability of containing the most nectar and pollen rewarding crop species. However, in our case we are confident we can exclude this effect. The highest rewarding crop species, oilseed rape, buckwheat and phacelia, occur in 33, 32 and 58 scenarios with *SurvProb* > 0, respectively, but they were also present in 271 (oilseed rape), 198 (buckwheat) and 254 (phacelia) scenarios without any surviving colony (*SurvProb* = 0). This, together with similar data for other crop species, makes us believe that our simulations are sufficiently balanced to buffer against sampling effects because the presence of high-rewarding crops (oilseed rape or buckwheat or phacelia or all three) did not guarantee colony persistence and also for the highest diversity level of eight crop species survival probability is relatively low.

### Effects of spatial arrangement and relative abundance of crops

Overall, both relative crop abundance and farmland structure showed the lowest impact on colony viability. However, details can matter. According to our results, presence of mid- and late-flowering species such as phacelia at low abundance (2% or 1%) can be sufficient to ensure required honey stores for colony survival even in intensively used agricultural systems. For such late-seasonal floral resources to be effective, they must be located close to the hive. If the distance of these late-flowering resources increases to > 1500 m from the hive, the foraging efficiency decreases and foragers are not able to gain a sufficient energy surplus if relying on this late-flowering resource alone.

Thus, in addition to temporal aspects of crop species leading to the presence or absence of forage gaps, spatial aspects also matter: distances of fields with critical crops preventing forage gaps (Table 3) should not exceed honeybees’ regular foraging distances as reported by Steffan-Dewenter and Kuhn (2003) and Couvillon et al. (2014). For example, even if the abundance of phacelia was 5% in cropping system of 8 crops, but the fields were 1347 - 3546 m away from the hive, the colony was not able to satisfy, in the long run, its late-seasonal demand for nectar to provide honey stores for overwintering (Tables S15-S23 and Figs. S20-S33 present analyses of stylized landscape where each crop species has just one field, which varies in size and distance to the hive).

One might argue that in many agricultural landscapes, field sizes larger than our maximum field size, 69 ha, exist. For the single colony represented by BEEHAVE, this does not matter because the forage provided by fields larger than 10 ha cannot be completely consumed by this colony. This figure might change if intra- and interspecific competition is considered, but it seems unlikely that changes in the maximum field size considered by us will affect our overall findings.

Crop relative abundance was not important in our simulations, but this might be due to the fact that we did not consider competition with other honeybees and wild pollinators. Mass-flowering crop fields providing high amounts of nectar and pollen during their blooming period at the landscape scale (e.g. one hectare of was not important in our simulations oilseed rape provides 300 l nectar and 105 kg pollen during its flowering period of 30 days) cannot be depleted by the bees of a single colony as the colony’s population of foraging bees is not able to collect all offered nectar and pollen within this short-term period. Still, even for monocultures of the highest rewarding crop, oilseed rape, maximum patch size (69 ha) and minimum distance (0.1 m) to the hive, colonies collapsed within the first year. This indicates that the long-lasting gap in forage supply from mid-May onwards after the blooming period of oilseed rape cannot be compensated even by large fields and shortest flight distance of the worker bees to the forage source (additional simulations with simplified, stylized landscapes with three or four crop species, are presented in the supplement S7 and provide further mechanistic insight into the consequences of forage gaps.)

Still, even in the case of temporal continuity in the nectar and pollen supply, the spatial arrangement of the rewarding flowers can matter as it determines the energy expenditure by the flying bees. If the energy costs for foraging trips to these flowers exceed the energy gain, the colony in terms of its force of foragers is weakened over time. Longer foraging distances increase forager mortality and reduce production of a new worker generation due to lowered nectar and pollen intake. For example, for a cropping system scenario composed of the eight crops dominated by rye-grass dandelion (65 % of the agricultural area and all other crops occupy 5 %) the colony collapsed in the third year at latest. Here, none of the five late-seasonal rewarding phacelia patches is located in a flight distance shorter than 1000 m.

### Effects of nectar and pollen supply in reality

Recent studies addressing bee performance in different landscape contexts have not directly considered nectar and pollen availability over time but used proportion of semi-natural habitats in agricultural landscapes (Steffan-Dewenter and Kuhn 2003, Sponsler and Johnson 2015) as a proxy for forage availability to assess farmland quality. How well these proxies work could be assessed with the modelling approach presented here, but goes beyond the scope of our study.

Nonetheless, we demonstrated that we can explain the effects of crop species identity and diversity on colony performance by determining frequency, timing and duration of temporary forage gaps and revealing their impacts on underlying cascading effects on life stages and work force of the modelled colony (Fig. 8). Some of these mechanisms, e.g. reduced number of nursing bees resulting in high larval mortality, are also pointed out by previous modelling studies of honeybee colony dynamics (Torres et al. 2015, Rumkee et al. 2015). Outcomes of this modelling approach reflected some patterns observed in reality, but in general we offer hypotheses regarding colony’s response to different cropping systems which can be tested with experiments.

In most temperate zones, semi-natural habitats are patchily distributed and sometimes rare in intensively managed agricultural systems. Forage dearth periods occur mainly in summer, but nectar and pollen availability to the bees may vary widely among locations and land-use types. However, forage supply from weeds and wild plants during gaps heavily depends on their diversity in agricultural landscapes. Our finding that landscape structure is less important than crop diversity and identity is confirmed by field studies which suggest that honeybee foraging is not that much affected by landscape structure as it is by seasonal patterns, whereby especially in summer (June and July) overall forage availability in the landscape is low and is most challenging (Steffan-Dewenter and Kuhn 2003, Couvillion et al. 2014).

If gaps in forage availability occur at a small landscape scale, then the colony may be buffered from strong effects on colony structure and dynamics due to the long-foraging range of honeybees (Beekman and Ratnieks 2000). However, during sensitive phases of colony development long-range foraging may weaken colonies, because larger foraging distances means higher mortality risk for foraging bees. Therefore, the spatiotemporal dynamics of nectar and pollen in the landscape are important to provide a sufficient energy surplus to the colony (Lonsdorf et al. 2009). Hicks et al. (2016) pointed out that continuous forage availability throughout the year is important for insect pollinator richness and abundances.

Of course frequently occurring forage gaps in agricultural landscapes may lead beekeepers to provide sugar solutions and pollen substitutes, and to move hives to temporary forage-rich floral resources within the landscape in order to avoid malnutrition and colony collapse. But, recent studies suggest that decline in honeybee health and hive numbers is due to long non-foraging periods caused by intensive agriculture that leads to poor floral diversity and monocultures at large landscape scales (Brown and Paxton 2009), and insufficient and untimely feeding (Brodschneider et al. 2010).

Our results indicate measures for avoiding forage gaps, especially by implementation of mid- and late-flowering crop species for boosting honeybee colony success in intensive agricultural systems. For instance, re-introduction of clover in legume-rich grass swards instead of growing ‘green deserts’ typical for European forage cropping systems, or growing phacelia and buckwheat (Decourtye et al. 2010, Sponsler and Johnson 2015), and increasing the abundance and diversity of forage-rich, semi-natural floral resources is needed to overcome temporary gaps in nectar and pollen availability in agricultural landscapes.

### Limitations and next steps

There are a number of limitations in the modelling framework presented here (see also Supplement S8). In our simulations we considered landscapes that are simplified in their crop composition, rotation and abundance of floral resources, and in the representation of semi-natural habitats. Data about flowering phenology and nectar and pollen rewards for different crop species and plant species of semi-natural habitats are scarce and hard to distil from literature and vegetation data bases (Baude et al. 2016). We therefore focused on a selection of the most important crops, pasture types, and minor and cover crops and we assumed constant crop species-specific nectar and pollen content over the blooming period. Due to the high intraspecific variation in reported data, we used values which are likely to be linked with good weather throughout the year, because the weather data we used (“Rothamsted (2011)”) represent largely beneficial conditions. Nevertheless, applied nectar data for crop cultivation and our simplified representations of semi-natural habitats are in the range of those reported by Baude et al. (2016).

Our results show that even agricultural landscapes of higher diversity may not fulfill the continuous nectar and pollen requirements of the colony over the whole season. Presence of the gaps in forage supply, even if they occur at the end of the foraging season, induce certain levels of stress to the colony but it can take two or more years to accumulate and to completely unfold its consequences to colony’s brood stages and work force. This is an important observation from the model to improve bee monitoring for risk assessment.

Furthermore, our simulation study pointed out the important aspect of land use patterns, which determines the continuity in forage quantity and quality according to colony’s nectar and pollen needs at certain times of development cycle, for risk assessment. In order to include this important driver for bee losses into population models we need the data for the most important land cover types about floral resources and cover of crops, semi-natural habitats, and weeds in high spatial resolution e.g. from remote sensing data on land cover types and plant covers from vegetation data bases or experts. Nectar and pollen data can be derived from quantitative surveys of crops, weed species and plants from meadows, forests and rural habitats. Given that such surveys are labor-intensive, pollen and nectar volume can be predicted from statistical models by using flower morphology and flower counts.

Moreover, information about farming practices on the landscape level and regional weather data and forecasts are required in order to predict the floral rewards under several climate conditions. Landscape-level risk assessment depends on robust information about exposure to pesticides and bee pathogens. Recent modeling studies highlighted that honeybee colonies might be able to tolerate more stress in forage-rich landscapes and low forage availability makes them more vulnerable to exposure and diseases (Thorbek et al. 2017, Horn et al. 2016, Becher et al. 2014). Parameterization of land use pattern-depended forage intake is therefore an important task to model predictions for risk assessment and management. Agatz et al. (2019) make specific recommendations for collecting data that would allow for further, more detailed validation and, hence, use of BEEHAVE.

BEEHAVE neglects differences in pollen qualities (e.g. amino acids) among different species. In reality honeybees rely on a high diversity of floral resources to meet their pollen requirements especially in spring. Higher pollen diversity enhances immunity to diseases and tolerance to pesticides (Di Pasquale et al. 2013) and ensures the health and sustainability of the colony. This cannot currently be reflected by the model.

A further important limitation is that we do not consider competition with other honeybee colonies and wild pollinators. Implementing realistic densities and distributions of colonies or apiaries will considerably increase the complexity of an already complex modelling framework, but this challenge needs to be tackled. Regarding competition with wild pollinators, currently available data, except for common bumblebee and solitary bee species, are scarce and need to be reflected via aggregated and conservative assumptions.

In our analysis, we deliberately ignored other stressors such as pathogens, certain beekeeping practices, and pesticide effects. It is therefore important to keep in mind that the predictions of colony performance presented here are relative, not absolute. Certain cropping systems reduce the frequency, timing and duration of temporary gaps in nectar and pollen supply affecting colony’s life stages and work force and enable higher viability and better performance than others, and this enables the colonies to cope better with additional stress.

## Conclusions

Awareness of the importance of the spatiotemporal continuity of floral resources for many insect pollinators has long been recognized (Van Bergen et al. 2013, Sponsler and Johnson 2015, Hicks et al. 2016, Alaux et al. 2017, Dozales et al. 2019, Guzman et al. 2019), but evidence of landscape context effects on honeybee colony performance and their underlying mechanisms are scant (but see Sponsler and Johnson 2015). Thus, we believe that our modelling framework improves our knowledge of how agricultural systems affect spatiotemporal forage availability and thus the performance of a single honeybee colony. Our most important general outcome is the ability to predict the extent to which forage gaps reveal detrimental cascading effects on colony dynamics and put honeybee colonies at risk, by using the availability of nectar and pollen over time.

Agricultural policies should adopt a way of assessing cropping systems and develop incentives or regulations that minimize forage gaps. These measures should address both cropping systems, and their diversity and composition, and the amount and spatial distribution of semi-natural habitats. For representative farmland structures and weather conditions in different eco-regions (EFSA 2016), the effects of policies and farmland structure and dynamics could be first screened by generating “forage provisioning plots” (Fig. 7A and 8A) and, if in doubt, assessed in more detail by using BEEHAVE.

Our results suggest that increasing crop number increases a colony’s viability, but the identity of crop species still determines the frequency, timing and duration of temporary gaps in nectar and pollen supply and thus the strength of cascading effects on the structure and dynamics of a honeybee colony. If the floral diversity of the cropping system does not ensure a continuous supply with nectar and pollen, especially during sensitive phases of colony development, colonies are negatively affected in terms of a reduction in worker bees for nursing and foraging tasks.

Overall, our framework can help to understand, quantify and rank the effects of different agricultural production systems on honeybee health, to predict colony responses to future anthropogenic land-use changes, and to test land-use management strategies and policies on maintenance of honeybees and other pollinators for assurance of pollination services in agricultural systems.

## Supporting information

Input file for NePoFarm

R script NePoFarm (rename to .R)

## Acknowledgements

This work was supported by grants from the German Academic Exchange Service (DAAD) and the Helmholtz Interdisciplinary GRADuate School for Environmental Research (HIGRADE). JO, MB and PK are supported by a BBSRC UK grant BB/K014463/1. We thank Rosalind Shaw for methodological support in QGIS and R applications to deal with data vector data. We thank B. Woodcock and one anonymous reviewer for their insightful and constructive comments.

