## Supplement for "Honeybee colony performance affected by crop diversity and farmland structure: a modelling framework"

Inhalt

### S1 Model parameters, equations and assumptions

**Initial settings**

The model parameters in BEEHAVE are shown in Table S1. Most of the BEEHAVE default settings were used: all simulations started on the 1^st^ January with an initial colony size of 10,000 worker bees, no infestation with virus-infected varroa mites and no beekeeping practices were included (see Appendix S5 in Becher *et al.* 2014).

To analyse effects of farmland structure, crop identity and crop diversity, data of forage availability in terms of nectar and pollen for various farmland scenarios were imported via input files containing information about size, location, crop species identity, amount of provided nectar and pollen, sugar concentration and detection probabilities throughout the year by setting INPUT_FILE TRUE. Weather defines the daily foraging period (Figure S1). We use real weather data of Rothamsted, UK (2011) representing beneficial annual weather conditions (Figure S1).

**Table S1:** Most important initial parameters. Bold parameter values mark deviations from the BEEHAVE default setting.

| **Parameter** | **Parameter value** | **Explanation** |
| --- | --- | --- |
| **Weather** | Rothamsted (2011) | weather option defining daily suitable weather conditions for foraging trips over the year |
| N_INITIAL_BEES | 10000 | initial colony size |
| N_INITIAL_MITES_HEALTHY | 0 | initial number of healthy mites |
| N_INITIAL_MITES_INFECTED | 0 | initial number of infected mites |
| CRITICAL_COLONY_SIZE_WINTER | 4000 | threshold colony size for winter survival on day 365 (31^st^ December) |
| SeasonalFoodFlow | **OFF** | if off / false: constant food availability of patch |
| QueenAgeing | OFF | if off / false: egg laying rate does not decrease with queen age |
| EggLaying_IH | ON | if on / true: egg laying is affected by available nurse bees |
| Swarming | No Swarming | Swarming options: no swarming at all |
| MAX_EGG_LAYING | 1600 | maximum egg laying rate per day (eggs/day) |
| MORTALITY_FOR_PER_SEC | 0.00001 | mortality rate of foragers per second foraging (1/s) |
| MAX_BROOD_NURSE_RATIO | 3 | maximum amount of brood, nurse bees can care for |
| ConstantHandlingTime | OFF | if off / false, handling time increases with depletion of flower patch |
| **Important parameter settings for nectar and pollen supply of farmland scenarios** | | |
| ReadInfile | **true** | Forage patch data are read from INPUT_FILE |
| INPUT_FILE | **e.g. “Input_simpleLand**  **85CE_5OSR_5M_**  **5SF.txt”** | contains data on nectar and pollen availability etc. for each of 171 (simple-structured farmland) or 1887 (complex-structured farmland) forage patches for each of 365days |
| patchType | **e.g. “Cereals”** | Crop species that occupies the patch |
| distance_m | **patch-specific** | Distance from the forage patch to the colony (m) |
| size_sqm | **patch-specific** | size of the forage patch (m²) |
| quantityPollen_g | **crop species- and patch size-specific** | Amount of daily available pollen at the forage patch depending on patch size (size_sqm) and crop species identity given in g |
| Concentration | **crop species-specific** | sugar concentration of provided nectar of crop species (mol / l) |
| quantityNectar_l | **crop species- and patch size-specific** | Amount of daily available nectar at the forage patch depending on patch size (size_sqm) and crop species identity given in l |
| calc_DetectProb | 0.2 | calculated detection probability that a scout finds the forage patch (calculated on the basis of the size of the forage patch and its distance to the hive determined in BEESCOUT model) |
| model_DetectProb | 0.2 | modelled detection probability that a scout finds the forage patch (determined in BEESCOUT ) |
| NectarGathering_s | 1200 | time to gather a nectar load on the forage patch (s) |
| PollenGathering_s | 600 | time to gather a pollen load on the forage patch (s) |

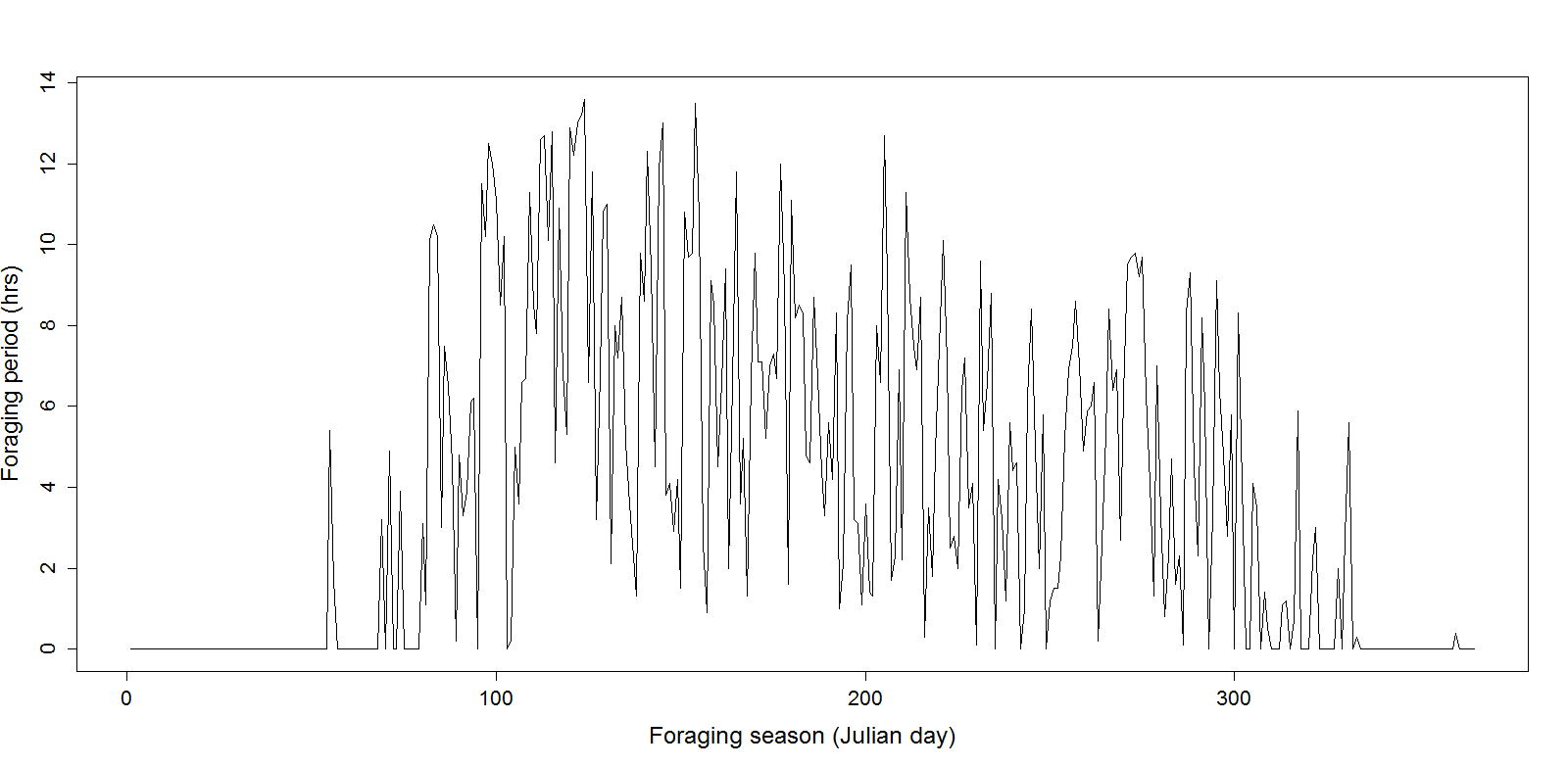

**Figure S1**: Daily foraging period in hours resulting from annual weather condition of used weather data of Rothamsted, UK (2011) throughout the year.

Table S2 shows important parameters and model equations of the BEEHAVE foraging module (Becher et al. 2014). Farmland context in terms of farmland structure, crop identity and crop diversity determine the spatio-temporal availability of nectar and pollen. The available nectar and pollen amounts at the forage patch can be completely depleted during a day, as a function of the crop species identity on the respective forage patch, i.e. whether this crop species provides copious or low amounts of nectar and pollen during its blooming period. Handling time, i.e. the time a forager needs to collect a nectar or pollen load at the patch, increases with the degree of forage depletion at this patch. Handling time in turn strongly influences the duration of a foraging trip. Thus, under low forage availability on the respective forage patch, foraging costs in terms of energy expenditure per trip tend to be higher.

Moreover, increasing foraging distance increases the duration of a foraging trip, the energetic costs for flying to and returning from the forage patch, and the mortality risk during a foraging trip, which increases with increasing time that a bee spends outside the hive.

**Table S2:** Important parameters and model equations of the BEEHAVE foraging module (Becher et al. 2014).

| Foraging parameter | Equation / parameter value | Explanation |
| --- | --- | --- |
| Handling time | set handlingTimeNectar  NectarGathering_s *  ((FlowerPatchesMaxFoodAvailableTodayREP who "Nectar") / quantityMyl)  analogous to handlingTimePollen (PollenGathering_s) | time a forager needs to collect a nectar or pollen load at the patch, proportionally increased with the depletion of the patch during the day |
| Trip duration | set tripDuration  2*distanceToColony*(1 / FLIGHT_VELOCITY ) + handlingTimeNectar | duration of a foraging trip depends on the flight distance from the hive to the patch, the flight velocity, and the handling time at the forage patch |
| flightCostsNectar | set flightCostsNectar  (2 * distanceToColony * FLIGHTCOSTS_PER_m) + (FLIGHTCOSTS_PER_m * handlingTimeNectar * FLIGHT_VELOCITY *energyFactor_onFlower) | flight costs are based on the energetic costs for flying to and returning from the patch and the energy spend at the patch while collecting nectar |
| FLIGHTCOSTS_PER_m | 0.000006531 kJ / m | energy consumption on flight per meter |
| energyFactor_onFlower | 0.2 | a bee saves energy while sitting on a flower to collect the nectar or pollen |
| mortalityRisk | set mortalityRisk  1 – ((1 - MORTALITY_FOR_PER_SEC) ^ tripDuration) | mortality risk during a foraging trip depends on the time that a bee spends outside the hive |
| quantityMyl  amountPollen_g | set quantityMyl  (quantityMyl - ( CROPVOLUME*SQUADRON_SIZE))    set amountPollen_g  (amountPollen_g - (POLLENLOAD*SQUADRON_SIZE)) | The amount of pollen or nectar available at the forage patch is reduced by the amount of pollen and nectar collected by a forager. |
| CROPVOLUME | 50 µl | volume of a forager's crop, is completely filled at flower patch |
| NectarGathering_s PollenGathering_s | 1200 s  600 s | time to fill crop with nectar / to collect a pollen load |
| FLIGHT_VELOCITY | 6.5 m/s | flight speed of a forager |
| MORTALITY_FOR_PER_SEC | 0.00001 s^-1^ | mortality rate of foragers per second foraging |
| SQUADRON_SIZE | 100 | number of foragers in the super-individuals "foragerSquadron" |
| POLLENLOAD | 0.015 g | amount of pollen collected during a pollen foraging trip |

### S2 Farmland maps

Here we provide detailed information on how to combine vector data shape files of area and route characters (freely available GIS data) into one area map including all field boundaries. Such a map describes a farmland template. We also describe how to analyse and extract the size and location of the field patches in these farmland maps. We created the farmland configuration maps from vector data by using R version 3.1.2 (R Development Core Team 2014) and required packages for spatial data, and QGIS version 2.6 (QGIS Development Team 2014). For extraction of field patch characters we used the model BEEScout (Becher et al. 2016).

**Combining vector data of area and route characters**

To create and rasterize farmland maps from OS VectorMap Local data, we combined several vector shape files representing area of a 5 km by 5 km farmlands into one Area layer containing all field boundaries between patches using R version 3.1.2 (R Development Core Team 2014) and required packages for spatial data, and QGIS version 2.6 (QGIS Development Team 2014) for its rasterization.

**Required OS VectorMap Local data and R-packages**

We chose freely available vector data of 5 by 5 km farmland areas (OS VectorMap Local, 2014) in UK for the Nocton Heath area (Lincolnshire) as a simple-structured farmland and for the Trenwheal area (Cornwall) as a complex-structured farmland and downloaded shapefiles from [www.ordnancesurvey.co.uk/buisness-and-government/products/vectormap-local.html](https://www.ordnancesurvey.co.uk/buisness-and-government/products/vectormap-local.html). The Nocton Heath area with the grid reference tf06sw represents a landscape with quite intensive agriculture (large fields resulting in simple structure), whereas the Trenwheal area with the grid reference sw63sw is a landscape with extensive agriculture (small fields resulting in complex structure).

**Required shape files from OS VectorMap Local**

To create an area layer of agricultural fields including all field boundaries, all area files containing polygons and all road files containing lines (General_Road_Casing and General_Line hereafter called as “GLC” and “GL”) for transfer into polygons are needed. We used the following area and line character shapefiles from OS VectorMap Local to create filed boundaries between recorded polygons:

- building
- general line
- general road casing
- landform area
- landform line
- railways
- road centerline
- water area
- urban

**Required R-packages**

For combining several shapefiles of area characters into one area layer, for merging area and road shapefiles together, and for rasterization several R-packages are necessary.

- sp (Bivand et al. 2013)
- maptools (Bivand and Lewin-Koh 2014)
- rgeos (Bivand and Rundel 2014)
- igraph (Csardi and Nepusz 2006)
- rgdal (Bivand et al. 2014)
- raster (Hijmans 2014)

**Combining shape files of area and route characters into one area layer shape file in R**

At first, we read GRC and GL in as shapefiles and made a list of shapefiles for area layer containing landform, buildings, water and urban polygons as follows:

*file.listArea <- c (list.files ("path/vml_666445/sw", pattern= "sw63sw_Landform_Area.shp$"),*

*list.files ("path/vml_666445/sw", pattern= "sw63sw_Building.shp$"),*

*list.files ("path/vml_666445/sw", pattern= "sw63sw_Urban_Extent.shp$"),*

*list.files ("path/vml_666445/sw", pattern= "sw63sw_Water_Area.shp$"))*

Then, we combined buildings, water area, landform area and urban area shapefiles into one combined area shapefile by setting up a load of blank fields to add features to the attribute table, reading an OGR data source and layer into a suitable vector object (can only handle layers with conformable geometry features, not mixtures of lines, points and polygons in one layer), searching for matches to argument pattern within each element of a character vector and matched polygons out of all files together into one file. The following steps are:

*polygons <- NULL*

*shp.id.list <- NULL*

*Fid.full <- NULL*

*Fid.temp <- NULL*

*FeatCode.full <- NULL*

*FeatCode.temp <- NULL*

*FeatDesc.full <- NULL*

*FeatDesc.temp <- NULL*

*n.polys <- NULL*

*id.list <- NULL*

*setwd ("path/vml_666445/sw")*

*data.first <- readOGR (file.listArea[1], gsub(".shp","",file.listArea[1]))*

*Fid.full <- data.first$Fid*

*FeatCode.full <- data.first$FeatCode*

*FeatDesc.ful l<- data.first$FeatDesc*

*n.polys <- length(data.first$Fid)*

*shp.id.list [1:n.polys] <- file.listArea[1]*

*polygons <- slot (data.first, "polygons")*

*data.second <- readOGR(file.listArea[2], gsub(".shp","",file.listArea[2])) # [2] file index*

*polygons2 <- slot(data.second, "polygons")*

*data.third <- readOGR(file.listArea[3], gsub(".shp","",file.listArea[3]))*

*polygons3 <- slot(data.third, "polygons")*

Each field gets taken from the temporary field and then appended to the full field.

*for (i in 2:length(file.listArea))*

*{*

*data.temp <- readOGR(file.listArea[i], gsub(".shp","",file.listArea[i]))*

*Fid.temp <- data.temp$Fid*

*FeatCode.temp <- data.temp$FeatCode*

*FeatDesc.temp <- data.temp$FeatDesc*

These fields set up the temporary field containing the data from each file to be added.

*Fid.full <- append (Fid.full, Fid.temp)*

*FeatCode.full <- append (FeatCode.full, FeatCode.temp)*

*FeatDesc.full <- append (FeatDesc.full, FeatDesc.temp)*

*n.polys <- NULL*

*n.polys <- length (Fid.temp)*

*n.ids <- length (id.list)*

*shp.id.list.to.add <- rep (file.listArea[i],n.polys)*

*shp.id.list <- append (shp.id.list,shp.id.list.to.add)*

*polygons <- c(polygons, slot (data.temp, "polygons"))*

*}*

Rename the IDs of polygons.

*for (i in 1:length (polygons))*

*{*

*slot (polygons[[i]], "ID") <- paste(i)*

*}*

*spatialPolygons <- SpatialPolygons (polygons)*

*spdf <- SpatialPolygonsDataFrame (spatialPolygons, data.frame (Fid=Fid.full, id_shp=shp.id.list,*

*FeatCode=FeatCode.full, FeatDesc=FeatDesc.full))*

Output of the combined area shapefile.

*setwd ("path")*

*writeOGR (spdf, dsn="name", layer="combined", driver="ESRI Shapefile")*

**Rasterization in QGIS**

We rasterized vector data files of GRC, GL and the combined area file in QGIS using a high raster resolution and the British National Grid as reference system.

**Raster reclassification in R**

Then, we read in all raster files (GL-raster, GRC-raster and area-raster) in R and aligned the extent and origin of the GL – and GRC-raster to the area raster by reclassification of the values in raster to 1 for central points and 0 for not included areas (i.e. field boundaries) as follows:

Read in all raster files using the raster function.

*e.g. r.Arc.GRC_NE<-raster("path/name_GRC.tif")*

Check extent and dimensions of GRC- and GL-rasters and shift extent to extent of area raster. e.g. *r.Arc.GRC_NE*.crop<-crop(*r.Arc.GRC_NE*, extent(r.Arc.Area_NE))

*r.Arc.GRC_NE.shift<-shift(r.Arc.GRC_NE.crop, x=-2.5, y=-2.5)*

Reclassify the values in the raster so that they can be used for clumping. Classification is: 1 for central points and 0 for not included areas

(see help http://gis.stackexchange.com/questions/69734/what-is-the-equivalent-of-arcpy-con-in-qgis-and-or-r-raster-package)

*Con=function(condition, trueValue, falseValue)*

*{*

*return(condition * trueValue + (!condition)*falseValue)*

*}*

e.g. *r.Arc.GRC_NE.reclass<- Con (r.Arc.GRC_NE.shift>15699, 0, 1r.Arc.GRC_NE.reclass)*

Afterwards we created an empty raster, assigned the cell values to 1 and merged this raster with all reclassified rasters (GL-reclass, GRC-reclass and area-reclass) using the merge function.

*ext.1<-extent(r.Arc.GRC_NE.shift)*

*one.raster<-raster(ext.1)*

*values(one.raster)=1*

e.g. *r.Arc.GRC_NE01<-merge(r.Arc.GRC_NE.reclass,one.raster)*

*writeRaster (r.Arc.GRC_NE01, "r.Arc.GRC_NE01.tif", format="GTiff",overwrite=TRUE)*

At next, all reclassified raster were merged together. All cells that are separated by 0 are grouped into shapes using the clump-function of the raster package that detects clumps (patches) of connected cells. These detected patches get an unique ID and 0’s are used to separate patches.

*allraster_NE01Add<-r.Arc.GRC_NE01+r.Arc.RL_NE01+r.Arc.Area_NE01*

*allraster_NE01Add.reclass<-Con (allraster_NE01Add==3, 1, 0)*

*allraster_NEAdd.clump<-clump(allraster_NE01Add.reclass, directions=4)*

Finally, this clumped raster was transferred into an .asci file using R, converted into vector format with adding an attribute table with a list of polygon ID’s and exported as a bitmap, jpeg, or tiff-file using QGIS.

*writeRaster (allraster_NEAdd.clump, "allraster_NEAddClump", format="ascii", overwrite=TRUE)*

**Calculation of patch sizes and distances using BEEScout (Becher et al. 2016)**

To calculate the size, location, and distance from the hive to the farmland patches of both farmland maps we used BEESCOUT, a spatially explicit, individual-based model, developed in the freely available programming language Netlogo (Wilensky 1999). BEESCOUT can be used to determine the number, location and size of forage patches for a bee colony (honeybees or bumblebees) on the basis of real or artificial landscape maps and to assess the probabilities of these forage patches being detected by a scouting bee (Becher et al.,2016). A detailed model description, following the ODD (Overview, Design concepts, Details) protocol (Grimm et al. 2006, 2010), the model itself, a list of all state variables, input maps and a user manual is provided in the Supporting information of the BEESCOUT model publication.

We imported both farmland maps as image files and adopted the scaling of the model world depending on the scaling of our farmland maps by relating the distance between two grids in the modelled map to the real distance between two patches. In Table S3 parameter settings for map analysis (location, size, distance of field patches) for both configuration maps are listed.

**Table S3**: Parameter settings for map analysis of both simple- and complex-structured farmland maps using BEEScout (Becher et al. 2016).

| Parameter | Setting |
| --- | --- |
| *Search options* | |
| SearchMode | known flower patch (colony) |
| N_Bees | 10000 |
| RandSeed | 1 |
| LinearisationFactor | 1.00 |
| DisplacementFactor | 1.0 |
| TripDuration_s | 1200 |
| ScoutingPeriod_hrs | 9 |
| RandomWalk | Off |
| RandomTripDuration | On |
| ImmediateReturn | Off |
| FixTurningAngle | Off |
| MaxForagingRange_m | 5000 |
| MaxVisitsColour | 1000 |
| *Definition Food Patches* | |
| YellowPatches | Off |
| GreenPatches | Off |
| BluePatches | Off |
| Lakes | Off |
| RedPatches | On |
| Red_min | 14.25 |
| Red_max | 15.5 |
| Start_R | 1 |
| Stop_R | 365 |
| Nectar_R | 1 |
| Pollen_R | 1 |
| t_Nectar_R | 1200 |
| t_Pollen_R | 600 |
| Conc_R | 1.5 |
| *Scaling and Hive Postion* | |
| ScaleDistance_m | 1170 |
| Scale_X1 | 103 |
| Scale_X2 | 150 |
| Col_X | 157 |
| Col_Y | 125 |

### S3 Crop species and their nectar and pollen data

**Selected crop species**

To analyse effects of various farmland contexts, and thus of forage availability, on honeybee colony performance we did not try to compile data on the crop species that were actually used in the two landscapes considered, because that would have yielded only two points in the multi-dimensional space of possible crop compositions. We assigned agricultural crops, livestock forage plants typical of intensively used pastures, and important minor and cover crops that are frequently used in Europe (EFG 2007, Statistisches Bundesamt 2014, Eurostat 2015, FAO 2015) (Fig. 2). Details on crop and forage plants most common in European agriculture are provided in Table S4 and S5 and in Figures S2-S7.

As crop species we considered cereals (wheat, rye, barley and oats were pooled together) providing no resources to the bees, oilseed rape (*Brassica napus* L.), maize (*Zea mays* L.), sunflower (*Helianthuus annuus* L.), and field bean (*Vicia* *faba* L.) as these are the most important crops in European agriculture (see Table S4 and Figs S2-S4).

**Table S4: Data on cultivated crops (harvested area in 10^6^ ha) in Europe from 2014 (FAO 2015, Eurostat 2015).**

| **Country / Region** | **Harvested area in 10^6^ ha** | | | | | |
| --- | --- | --- | --- | --- | --- | --- |
|  | Cereals | Maize | Oilseed rape | Sunflower | Field bean (in ha) | Buckwheat (in ha) |
| France | 9.6 | 0.1 | 1.5 | 0.66 | 74900 | 30100 |
| Germany | 6.5 | 0.5 | 1.39 | 0.02 | 20500 | NA |
| Italy | 3.4 | 0.9 | 0.017 | 0.11 | 57700 | NA |
| Czech Republic | 1.4 | 0.1 | 0.39 | 0.02 | 0 | 1000 |
| Poland | 7.5 | 0.7 | 0.95 | 0 | 13500 | 62710 |
| Slovenia | 0.1 | 0.04 | NA | 0 | 100 | 1551 |
| Spain | 6.3 | 0.4 | 0.043 | 0.78 | 22800 | NA |
| Switzerland  (non-EU member) | 0.1 | 0.01 | 0.023 | 0 | 493 | NA |
| United Kingdom | 3.2 | 0.2 | 0.67 | NA | 107000 | NA |
| Ukraine (non-EU member) | 14.4 | 4.6 | 0.86 | 5.21 | 2200 | 136700 |
| European Union | 58.1 | 9.6 | NA | 4.22 | 396900 | 143847 |

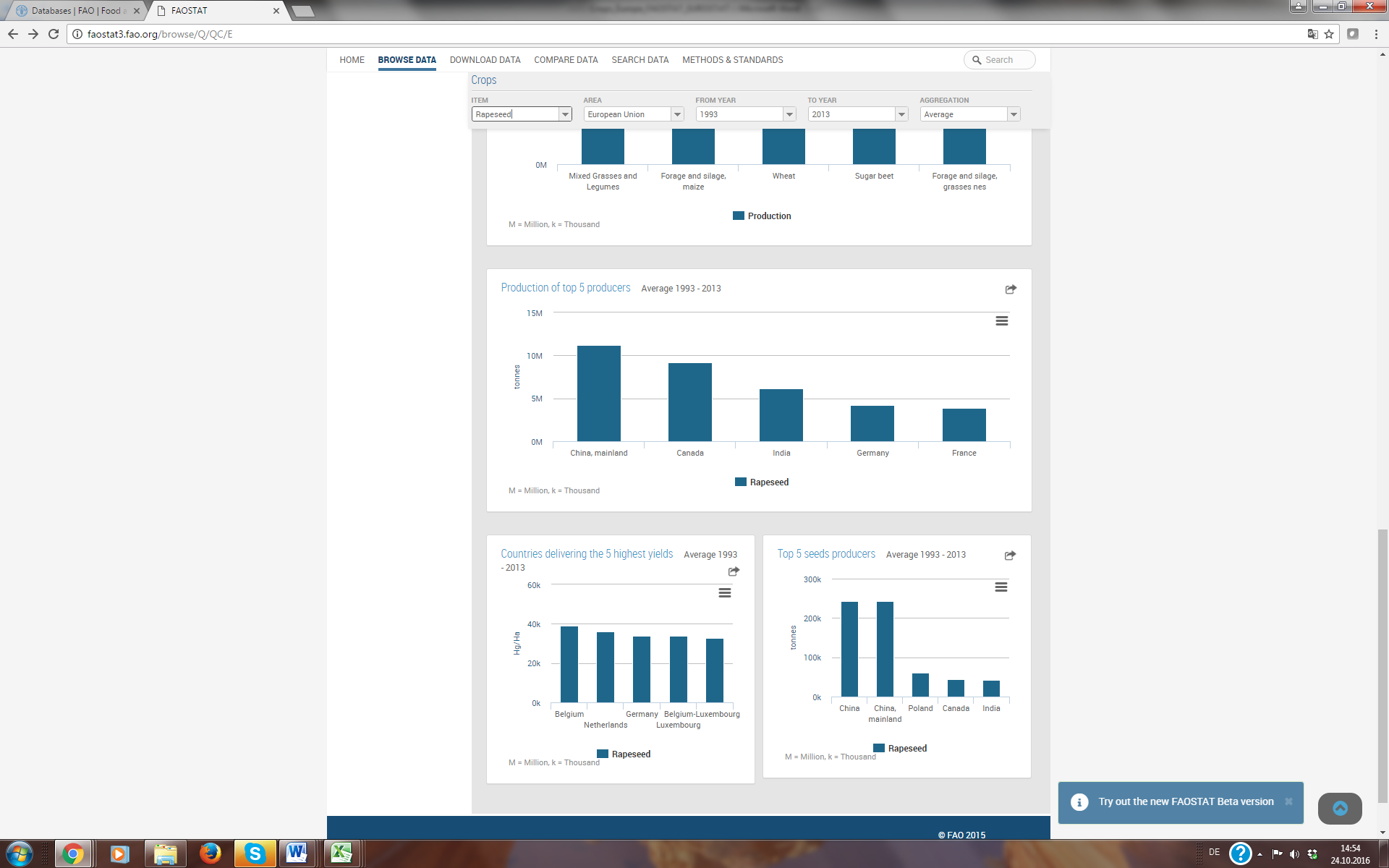

**Figure S2:** Top producers of rapeseed (© FAO 2015).

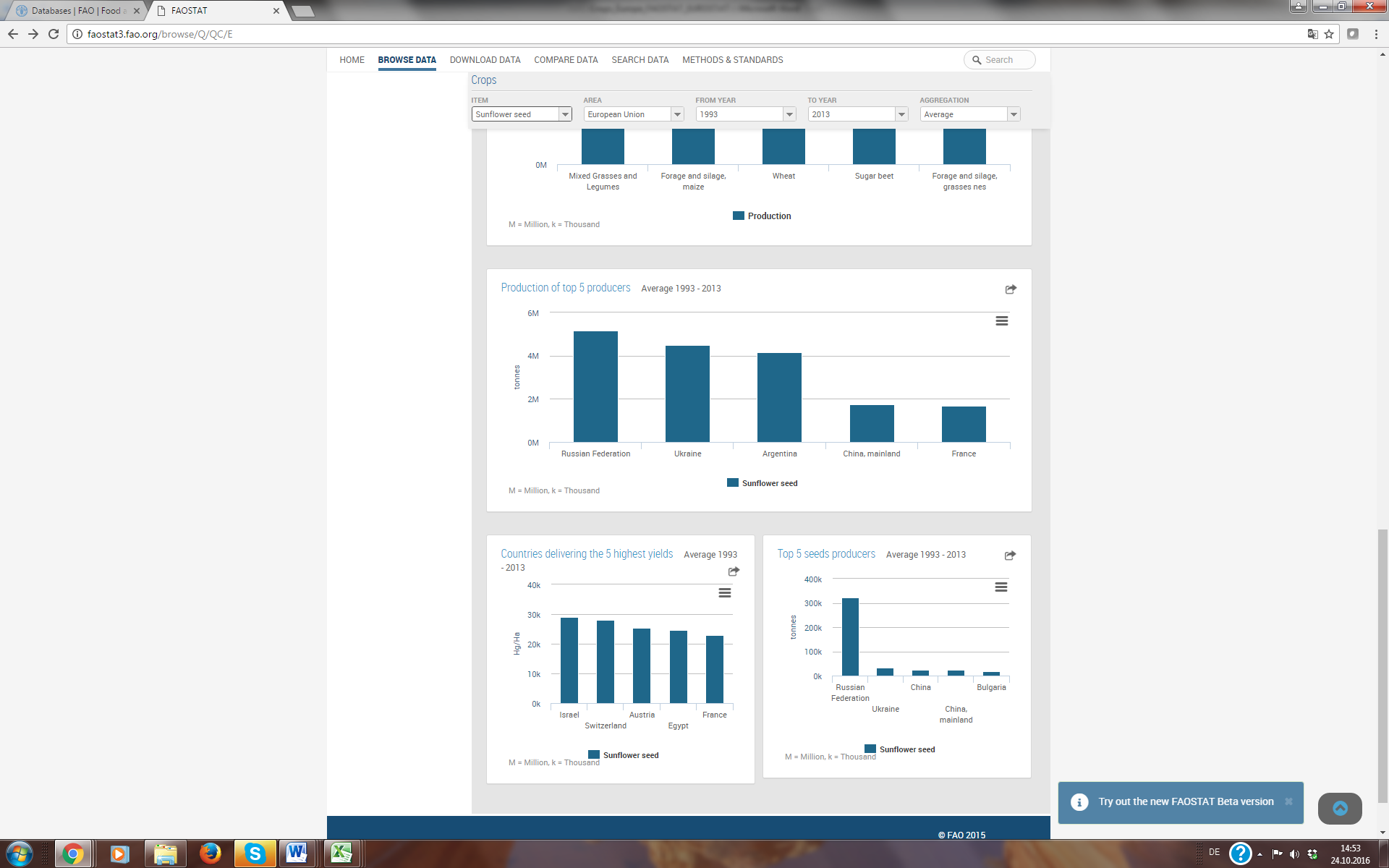

**Figure S3:** Top producers of sunflower (© FAO 2015).

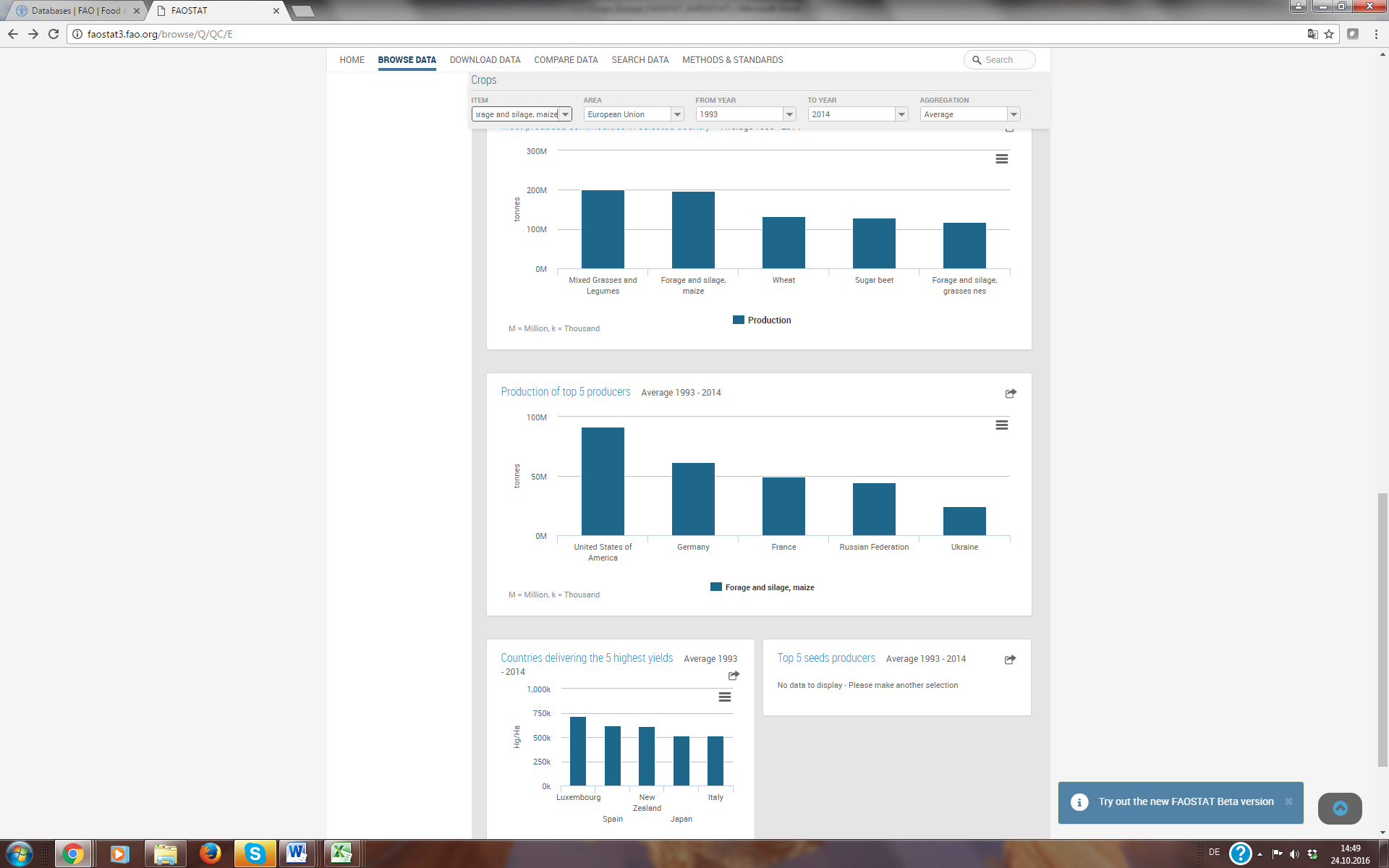

**Figure S4:** Top producers of maize silage (© FAO 2015).

We further considered three different kinds of temporary grasslands typical in many temperate parts of Europe representing the most common intensively used pastures for grazing, forage harvesting, silage and hay production (EFG 2007). Most temporary grasslands are solely sown with grasses. The most important cultivated grasses in grassland and forage cropping are rye-grasses (e.g. *Lolium perenne*) whereas other grasses such as *Dactylis glomerata* are rarely sown (EFG 2007, Eurostat 2015). Especially, if pastures in arable land are lasting for a longer time, perennial grasses (mainly *Lolium perenne*) combined white clover (*Trifolium repens*) are preferred (EFG 2007). White clover as the most important leguminous plant in a pasture of grass-clover mixture will ensure a high forage quality improving both energy and protein content for livestock (Hall 1993). As shown in Figures S5 and S6 Germany, France, Denmark and Czech Replubilic are top producers of rye-grasses and clovers as forage crops. As third temporary grassland type, we implemented long-lived perennial dandelion (*Taraxacum officinale*), that is common along roads and field margins, infesting nearby sown grass pastures (Gibson 1997). As this weed is used by cattle grazing on pastures as readily as grasses and as herbicide usage for control is usually unnecessary, because up to 28 % abundance on total vegetation cover in rye-grass pastures total yields are not affected (Bergen et al. 1990), we considered dandelion as a valuable forage plant for livestock invading rye-grass pastures.

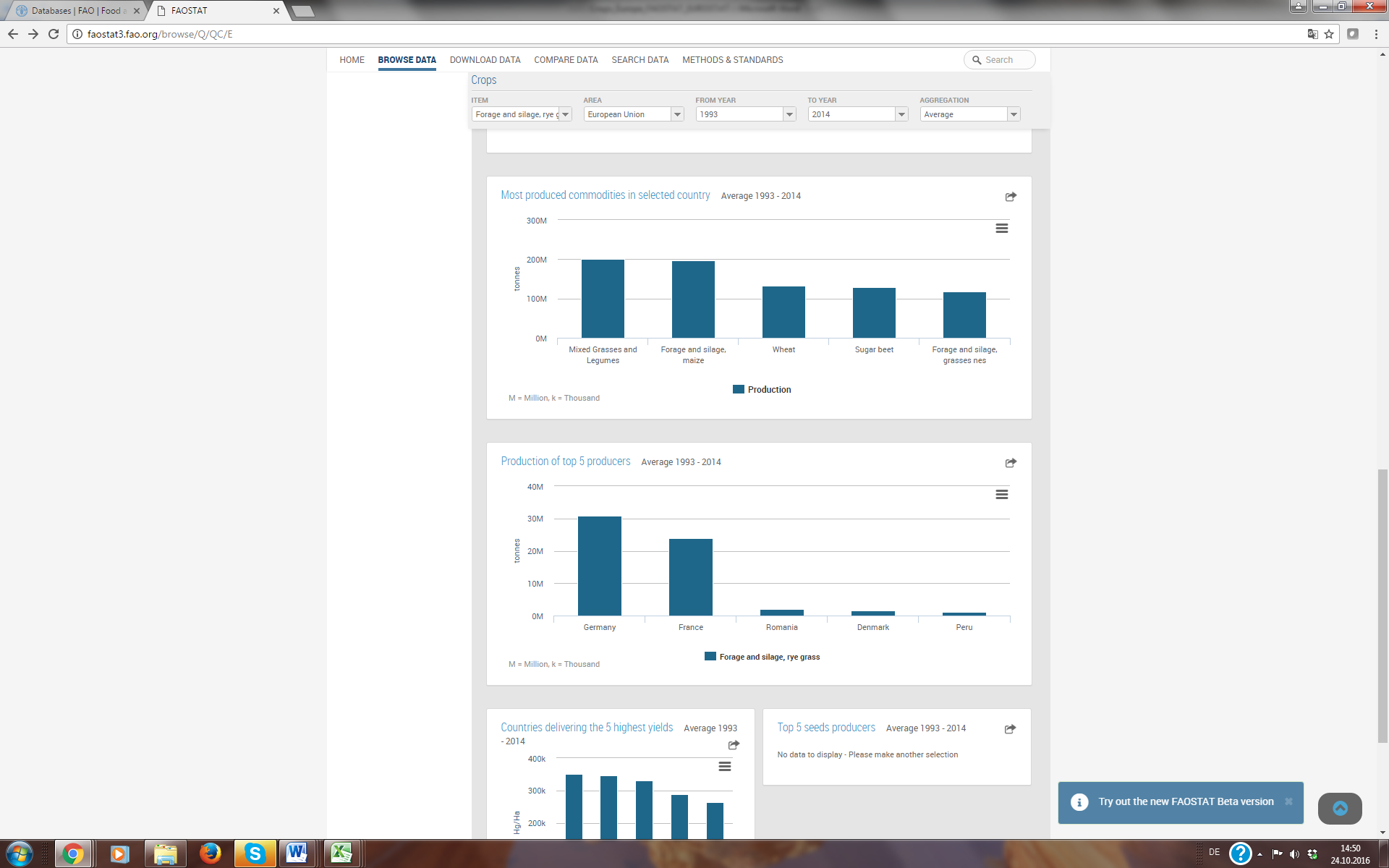

**Figure S5:** Top producers of rye-grasses as forage crops (© FAO 2015).

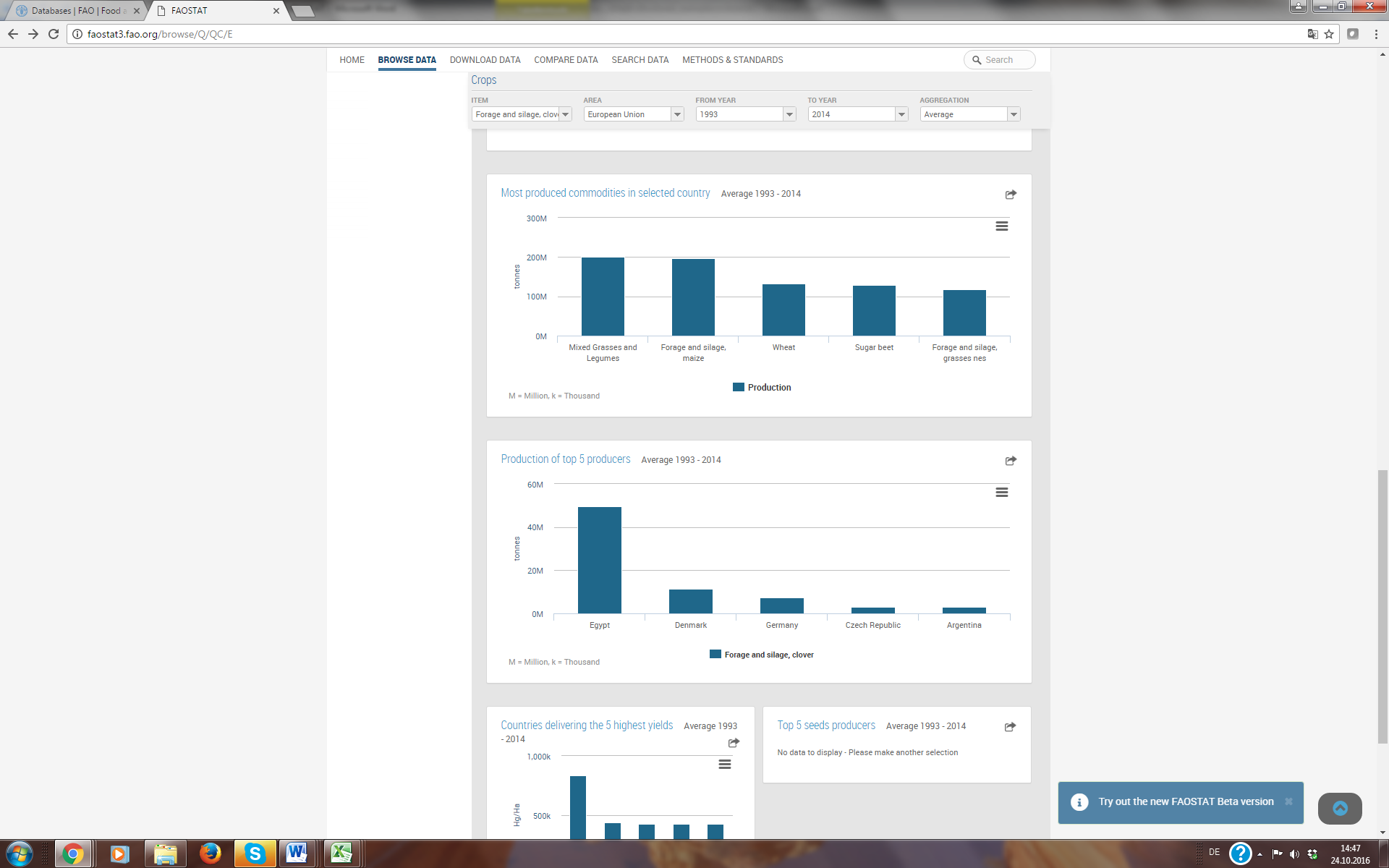

**Figure S6:** Top producers of clover as forage crop (© FAO 2015).

We further implemented important examples of European minor and cover crops. Phacelia (*Phacelia tanacetifolia*) is an annual plant with rapid flowering life cycle, which is used extensively in Europe as a cover crop in annual cropping systems grown to improve soil quality and environmental benefits to insects such as the honeybee (e.g. Decourtye et al. 2010, Baets et al. 2011). Buckwheat (*Fagopyrum esculentum*) is an important minor crop in Europe grown for grain-like seeds (Michalova 2001), whereby major producers are France, Ukraine and Poland (FAO 2015, Eurostat 2015) as shown in Figure S7.

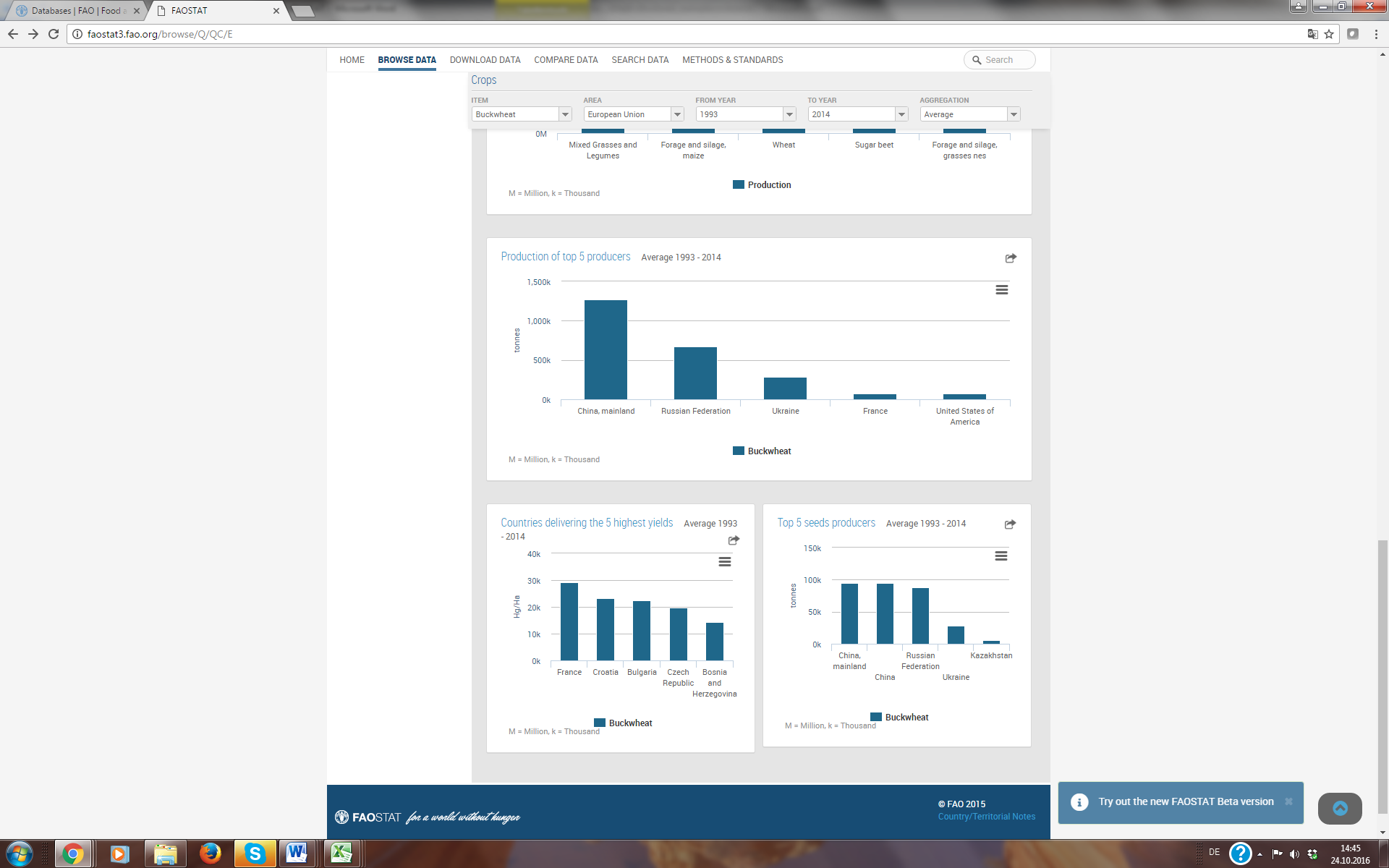

**Figure S7:** Top producers of buckwheat as minor crop (© FAO 2015).

To show abundances of different crop types on total agricultural area exemplary for European agricultural systems, we provide here some German statistics about abundance of different crop types on agricultural landscapes at the district level (Statistische Ämter der Länder und des Bundes (2014) www.regionalstatistik.de/genesis/online/). On average 56 % of arable land is cultivated with cereals, followed by 21 % maize (silage and corn), 22.5 % feeding plants (mainly grasses as forage crops), 10.3 % oilseed rape and oil fruits (e.g. sunflower). Only 0.8 % of arable land in Germany is cultivated with legumes (clover, field bean). Sugar beets and potatoes occupy on average 3.4 and 2 % of German agricultural area, but are yielded before they start blooming. On average 7.2 different crop species are cultivated per German district, whereby number of flowering crops varies from 0 to 5 averaging 4.2 crops (Table S4, Figs S2-S7). However, averaged proportion values are at regional level of overall districts and urban districts ranging from 288 ha (Offenbach am Main – urban district) up to 175902 ha (Uckermark - district) of land-use area (forest area, urban and circulation area, water area, and others are not included = approx. 47 % of total area). So, crop proportion on arable land varies widely among different regions (Table S5, Figs. S8-S13).

**Table S5: Number of crop types and their abundances on arable land in Germany according to Statistische Ämter der Länder und des Bundes (2014). www.regionalstatistik.de/genesis/online/**

| **Proportion on arable land** | **# districts** | **mean** | **sd** | **min** | **max** |
| --- | --- | --- | --- | --- | --- |
| **# crop species** | 411 | 7.2 | 2.2 | 0 | 9 |
| **# flowering crop species** | 411 | 4.1 | 1.2 | 0 | 5 |
| **% cereals** | 382 | 56 | 11.4 | 0 | 78.6 |
| **% maize (silage)** | 372 | 15.6 | 15.5 | 0 | 80.6 |
| **% maize (corn)** | 287 | 5.1 | 8.8 | 0 | 60.3 |
| **% oilseed rape** | 334 | 10.3 | 6.7 | 0 | 28.4 |
| **% oilfruits**  **(rapeseed, sunflower etc.)** | 288 | 10.3 | 6.6 | 0 | 28 |
| **% feeding plants**  **(grass, treifoil-grass, clover)** | 388 | 22.5 % | 15.5 | 0 | 80.6 |
| **% leguminous**  **(field bean, clover etc.)** | 307 | 0.8 | 0.8 | 0 | 7 |
| **% sugar beet** | 339 | 3.4 | 5.1 | 0 | 25 |
| **% potatoes** | 344 | 2.0 | 3.6 | 0 | 27.7 |

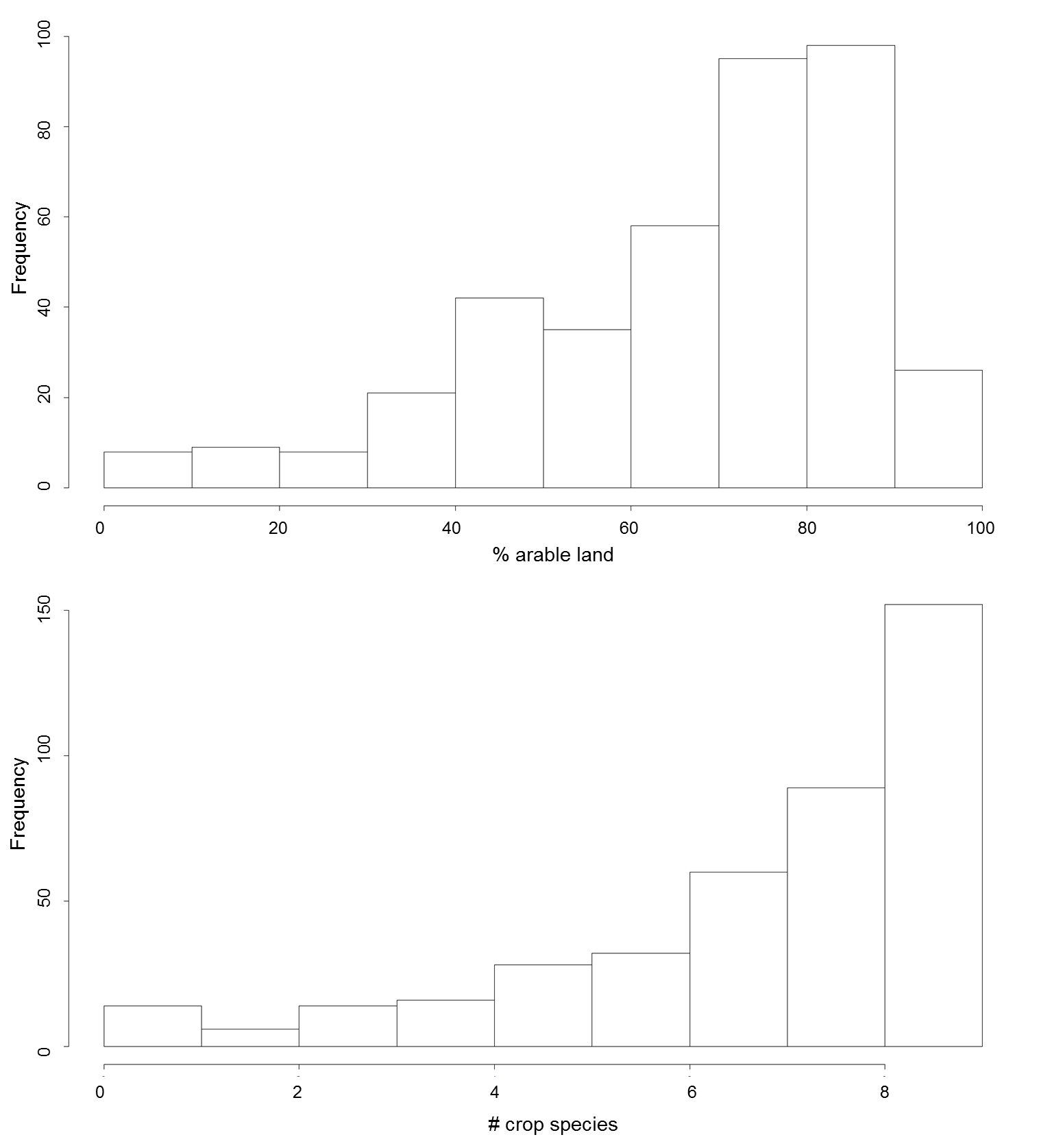

**Figure S8**: Histogram of proportion of arable land (top) and crop species number on arable land (bottom) for all 411 German districts.

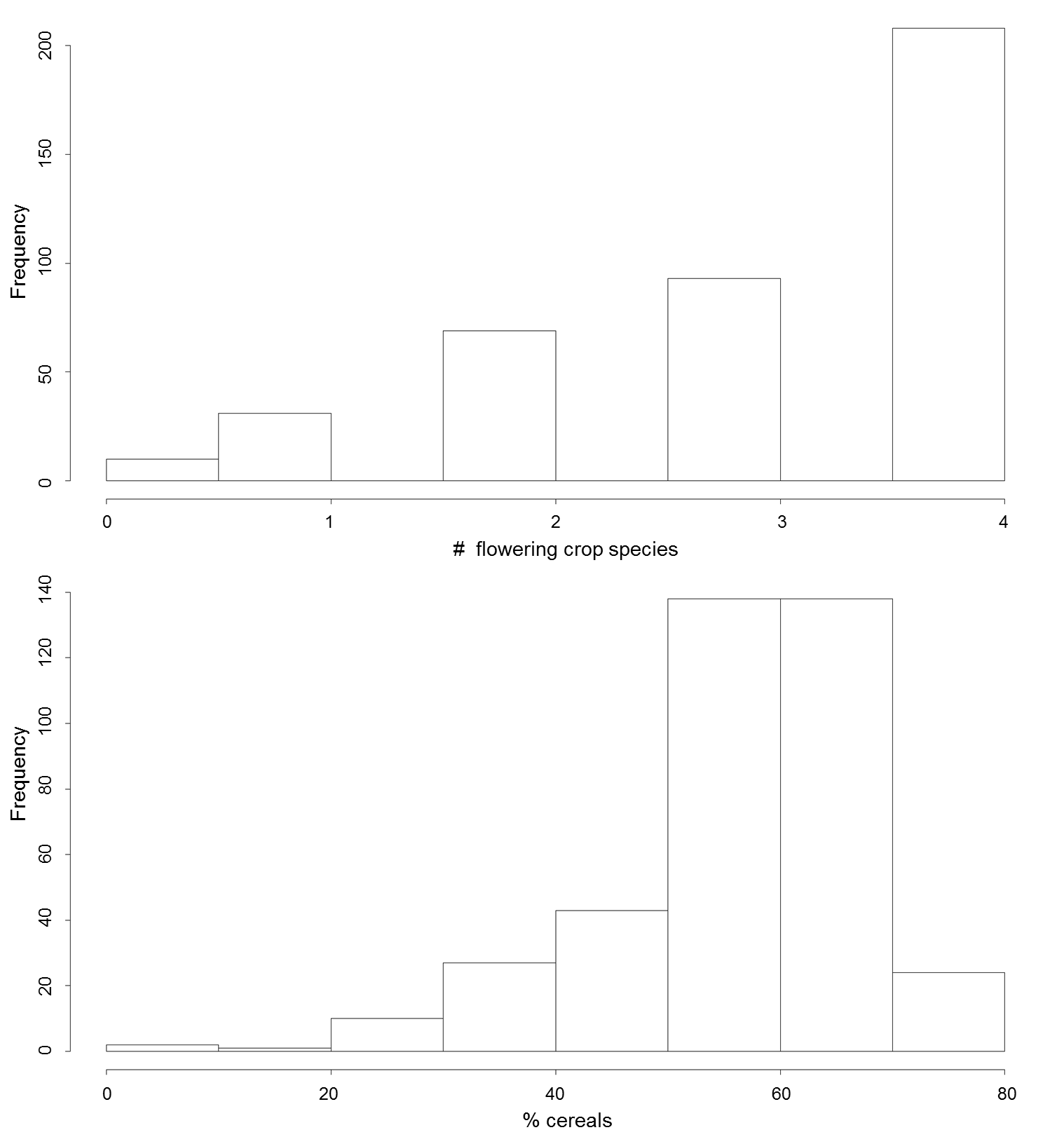

**Figure S9:** Histogram of flowering crop species number and proportion of cereals on arable land for all 411 German districts.

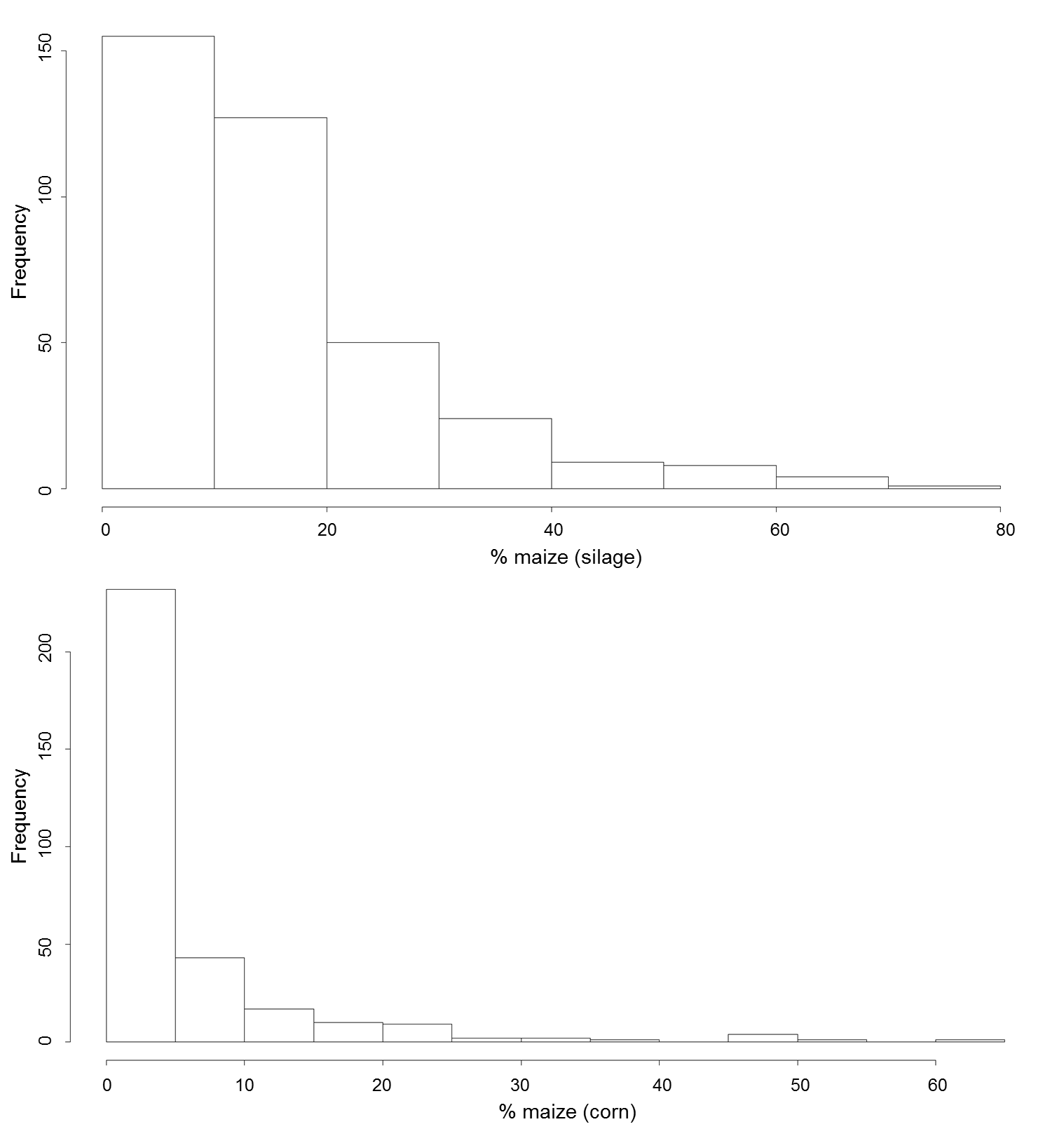

**Figure S10:** Histogram of proportion of maize (silage and corn) on arable land for all 411 German districts.

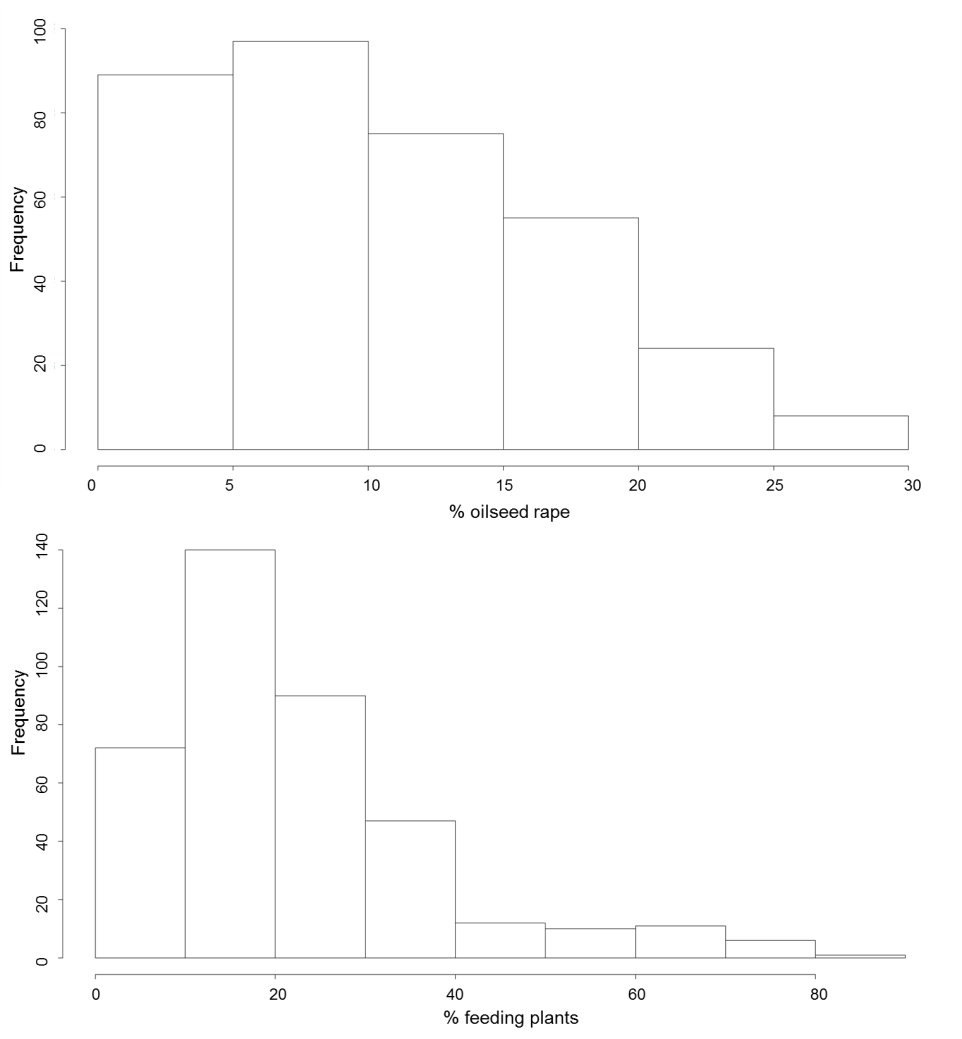

**Figure S11: Histogram of proportion of oilseed rape and feeding plants (forage crops such as grass, trefoil-grass, clover) on arable land for all 411 German districts.**

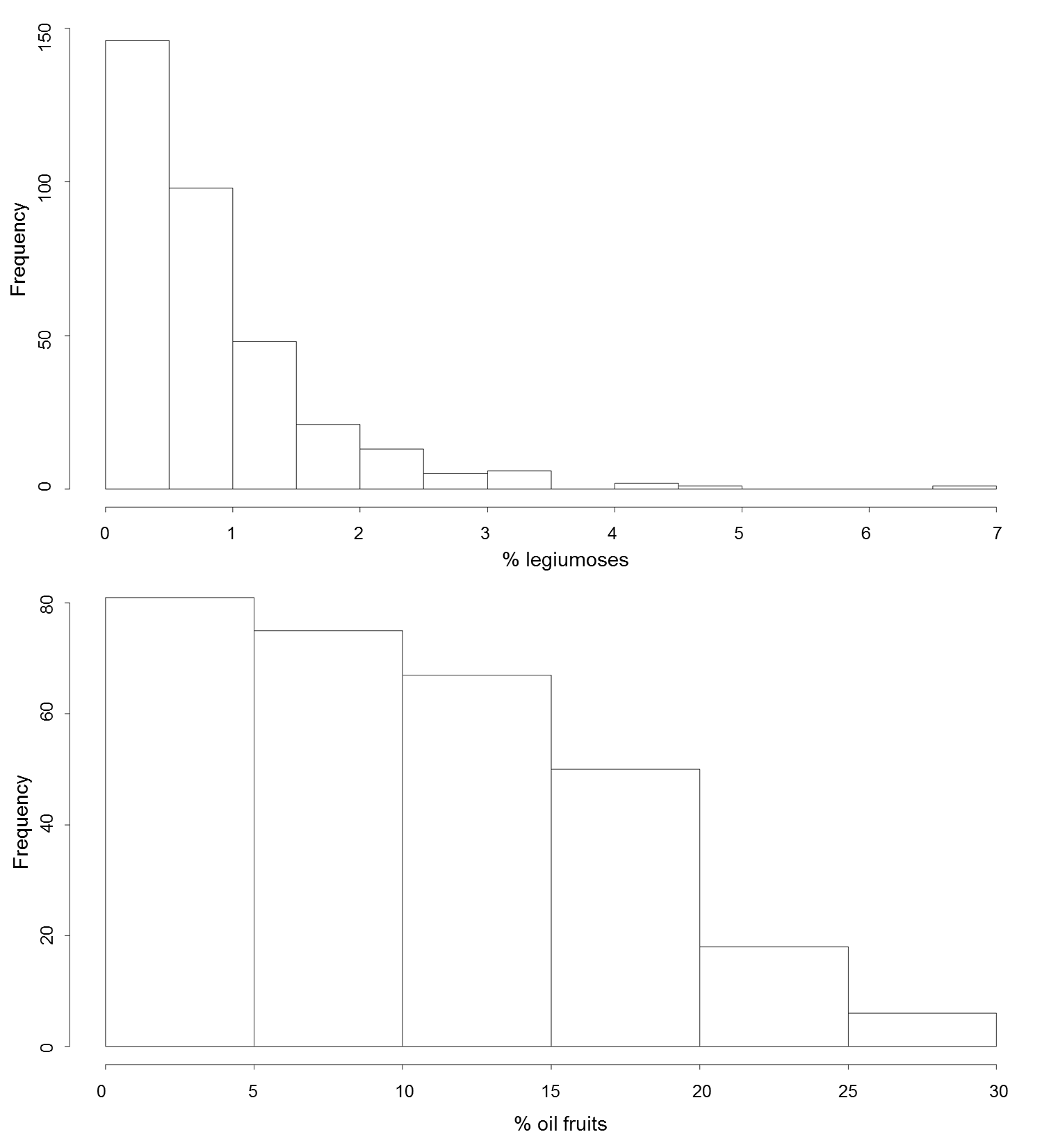

**Figure S12: Histogram of proportion of legumes and oil fruits (sunflower, rapeseed etc.) on arable land for all 411 German districts.**

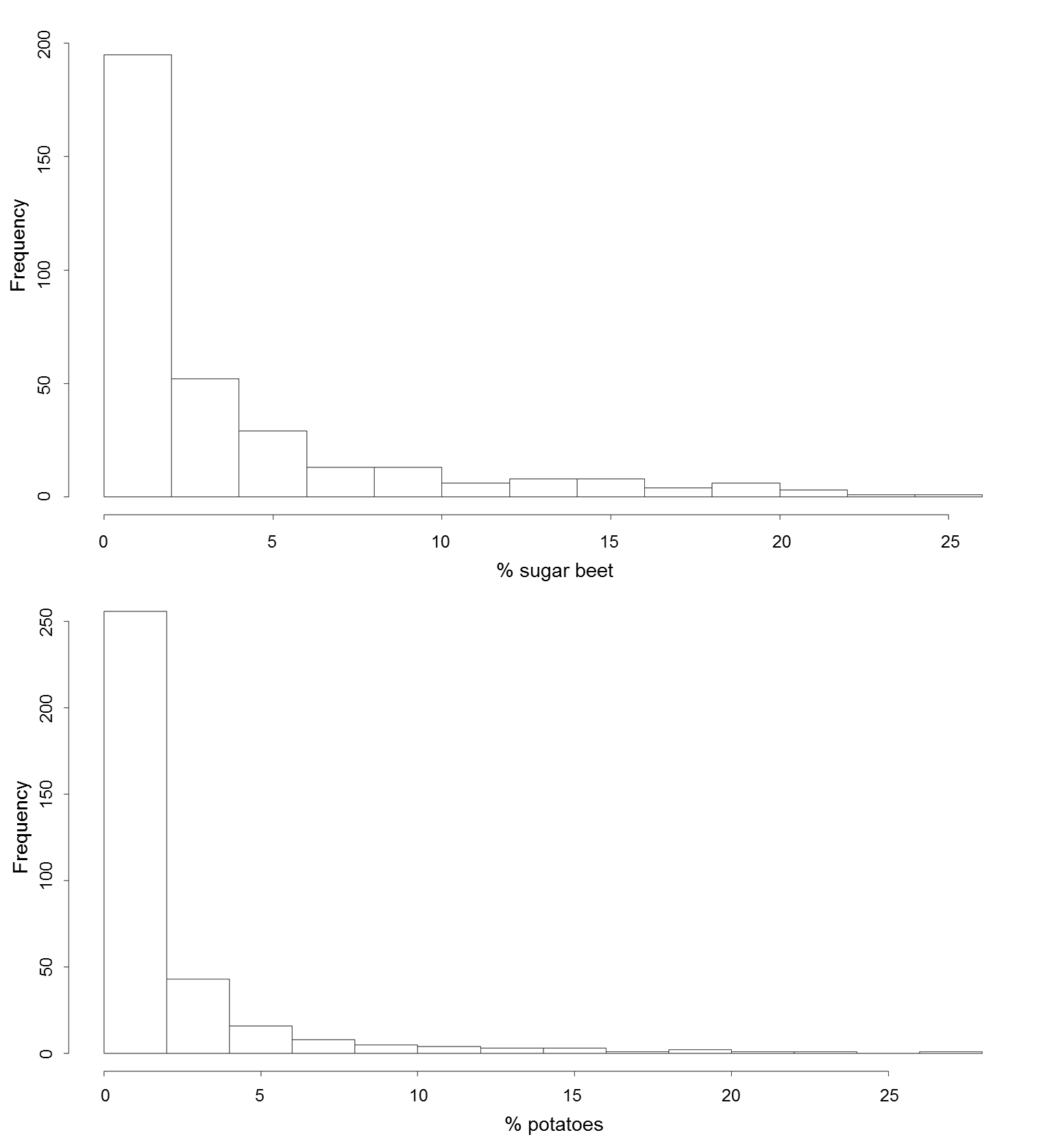

**Figure S13: Histogram of proportion of sugar beets and potatoes on arable land for all 411 German districts.**

**Crop species-specific nectar and pollen data as inputs for BEEHAVE**

To translate the identity of crop species’ on a given patch to units of nectar and pollen that could be collected by honeybees, we distilled species-specific flower density, nectar and pollen contents from existing literature and summarized these literature data in Table S7. We used the flowering period, defined by start and end day, the amount of nectar and pollen provided per day and the sugar concentration of each crop species as input data for BEEHAVE. As there is a lack in quantitative and qualitative data of floral rewards, the presented nectar and pollen data are quite uncertain and more data of this type are needed for a more accurate assessment. But presented data are in the range of nectar productivity for arable land according to Baude et al. (2016). An overview of literature data and equations for calculations of species-specific nectar and pollen contents is listed in Tables S6 and S7). In order to convert available data to required input format of BEEHAVE we made simplified assumptions. For example for converting sugar concentration from frequently reported weight percentage [%] w/w (g sugar/100 g nectar) into BEEHAVE (required concentration of mol / l) we assumed the following: nectar is an aqueous solution that mainly consists of water and sugar (Sucrose ---> Glucose + Fructose) and too much lower proportions of proteins, amino acids and vitamins. Detailed information about density of real nectar solutions (due to greatly varying water and sugar content) and about ratio Sucrose/ Fructose + Glucose is lacking. Therefore, we used a density table of sucrose solutions, the molar mass of sucrose and the molarity equation as approximation (see Table S6).

**Table S6**: Equations of nectar and pollen data for specific BEEHAVE input.

|  | Equation formula | Used assumptions, and units | Explanation |
| --- | --- | --- | --- |
| N_Flowers_plant_day_ | - | single floret (e.g. oilseed rape) or head = composition of single florets (e.g. sunflower) | Daily number of open flowers per plant |
| N_Plants_m²_ | - | e.g. oilseed rape (30 plants per m²) | Number of plants per m² |
| V_Nectar_flower_day_ | - | [µl] per single floret or head in 24 hours | Amount of nectar of a single flower unit [µl] |
| M_Pollen_flower_day_ | M_Pollen_flower_day =_  M_Pollen_flower_flowering_ / T_Flowering_ | when only pollen amount over whole flowering period is given  [mg] per single floret or head in 24 hours | Amount of pollen of a single flower unit [g] |
| N_Flowers_m²_day_ | N_Flowers_m²_ =  N_Flowers_plant_day *_ N_Plants_m²_ | Assuming constant number of daily opening flowers over flowering period | Daily number of open flowers per m² |
| V_Nectar_m²_day_ | V_Nectar_m²_ =  N_Flowers_m²_ _*_ V_Nectar_flower_ | Assuming constant number of daily opening flowers and constant daily nectar content of florets over flowering period | Daily nectar amount per m²  [l / m²] |
| M_Pollen_m²_day_ | M_Pollen_m²_ =  N_Flowers_m²_ _*_ M_Pollen_flower_ | Assuming constant number of daily opening flowers and constant daily pollen content of florets over flowering period | Daily pollen amount per m²  [g / m²] |
| C_Sugar_ | C_Sugar =_ 10 * c[%] * ρ  M | Assumed to be constant over flowering period.  To convert sucrose equivalents in weight percentage [%] w/w (g sugar/100 g nectar) [mol / l]:  c = concentration of solution [%]  ρ = density [kg / l], density of an aqueous solution consisting of sucrose is changed depending on its sugar concentration (and temperature  M = Molar Mass [g/mol]  M_Sucrose_ [g/mol] = 342.3 g / mol | Sugar concentration  [mol / l] |

Another example, maize provides only large amounts of pollen (no nectar). The pollen amount of maize in average was given in amount of pollen per plant in total, i.e. over whole flowering period, 3.5 g per plant in total according to Nowakowski and Morse (1982); and in amount of pollen per flower in total (494 mg according to Percival 1955). Data of pollen amount per flower per day were not available. Using amount of pollen per plant in total, amount of pollen per flower in total and flowering time per flower (14 days according to Emberlin 1999) we calculated the amount of pollen per flower per day. To calculate the maximum amount of pollen per plant in total we used the maximum number of pollen grains per plant (50 * 10^6^ pollen grains per plant in total according to Miller 1985), the weight of one maize pollen grain (0.00025 mg according to Miller 1985), number of flowers per plant (7.1) and flowering time per flower (14 days).

The daily nectar amount per head of sunflower was given in mg / head (mean = 0.29 mg / head and max = 0.5 mg / head). For calculation of nectar amount per head in µl we used the density ρ_nectar_ = 360 mg / ml and m_nectar_ = 0.29 mg / head (or 0.5 mg head) to solve the equation V_nectar_ = m ⃰ ρ (see Table S7).

**Table S7: Literature overview of flowering, nectar and pollen data of important crop species**: The duration of flowering is given in days. The daily amount of nectar (µl per flower unit per day) and pollen (mg per flower unit per day) refer to flower heads (composition of several single florets) for sunflower and maize. The nectar and pollen amounts of oilseed rape, white clover and field bean refer to single florets. Sugar concentrations (%) were converted into mol / l using the molar mass of sucrose (M = 342.3 g / mol).

| Crop | Period | Flowering  [days] | nectar  [µl/flower/day] | | | pollen  [mg/flower/day] | | | Concentration  [%] | | # Flowers per m² per day | | Reference |
| --- | --- | --- | --- | --- | --- | --- | --- | --- | --- | --- | --- | --- | --- |
|  |  |  | **Min** | **Mean** | **Max** | **Min** | **Mean** | **Max** | **Min** | **Max** | **Min.** | **Max.** |  |
| Oilseed rape ** (*Brassica napus*) | April - May | 22  22 – 45^a^ | **0.35**^b^ | 0.55^b^ | **0.82**^b^ | 0.187^d^ | 0.239^d^ | 0.292^d^ | 44^e^ | 59^e^ | 543^c^ | 1194^c^ | ^a^ Radchenko 1964  ^b^ Hedke 2000  ^c^ Blazyte-Cereskiene et al. 2010  ^d^ Von der Ohe et al. 1990  ^e^ Maurizio and Schaper 1994 |
| Maize ***  (*Zea mays*) | June - Sep | 14^a^ | - | - | - | 16^b^ | 35.3^c^ | 125.7^c^ | - | - | 21^b,d^ | 64^b,d^ | ^a^ Emberlin 1999  ^b^ Percival 1955  ^c^ Miller 1985  ^d^ Olson and Sanders 1988 |
| Sunflower ^&&^ (*Helianthus annuus*) | Aug -Oct | 28  (19 - 36)^a^ | 0.22^b,c^ | 0.81^b,c^ | 1.39^b,c^ | 26.6^d^ | 28.7^a,d^ | 30^a^ | 24^b^ | 61.3^c^ | 1.5^e^ | 6^e^ | ^a^ Minckley et al. 1994  ^b^ Hedtke 1998  ^c^ Zajácz et al. 2006  ^d^ Percival 1955  ^e^ AOF 2009 |
| Field bean  (*Vicia faba*) | June | 30  30 – 39^a^ | 0.19^b^ | 0.86^a^ | 4.44^c^ | 0.6^e^ | - | 0.7^c^ | 6^f^ | 50^f^ | 80^a^ | 135^a^ | ^a^ Brown and Scott 1992  ^b^ Pierre et al. 1996  ^c^ Prabucki et al. 1987  ^e^ Percival 1955  ^f^ Osborne et al. 1997 |
| White clover (*Trifolium repens*) | May - Oct | 102^a^ | 0.02^b^ | 0.1^b,c^ | 0.18^c^ | - | 0.019^c^ | - | 37^c,d^ | 65^b^ | 247^e^ | 741^e^ | ^a^ Percival 1950  ^b^ Weaver 1965  ^c^ Aleck 1997, unpubl.  ^d^ Montgomery 1958  ^e^ Free 1993 |
| Phacelia  (*Phacelia tanacetifolia)* | May - Dec | 60^a^  37^b^ | 0.07^b^ | 0.1475^b^ | 0.225^b^ | - | 0.3^c^ | - | 8.9^d^ | 35.9^d^ | 2000^a^ | 4000^a^ | ^a^ Williams and Christian 1991  ^b^ Petanidou 2003  ^c^ Percival 1955  ^d^ Zimna 1960 |
| Dandelion  (*Taraxacum officinale*) | April – June | 54^a,b^  25^c^ | 3.7^c^ | 5.55^c^ | 7.4^c^ | - | 1.2 | - | 16^c^ | 61^c^ | 8.9^c^ | 59.2^c^ | ^a^ Radford et al. 1968  ^b^ Bergen et al. 1990  ^c^ Szabo 1984 |
| Buckwheat  (*Fagopyrum esculentum*) | June - Sep | 89^a^ | 0.142^b,c^ | 0.307^b,c^ | 0.472^b,c^ | 0.110^a^ | 0.173^a^ | 0.236^a^ | 35^d^ | 45^d^ | 6543^e^ | 15259^e^ | ^a^ Alekseyeva and Bureyko 2000  ^b^ Kirillenko and Bockkareva 1983  ^c^ Sim and Choi 1999  ^d^ Jablonski and Szklanowska 1987  ^e^ Free 1993 |

### S4 Landscape generator NePoFarm to create scenarios as input files for the simulation model BEEHAVE

**Purpose**

The landscape generator NePoFarm was implemented in the freely available programming language R for creation of farmland scenarios as input files for analyses in the honeybee simulation model BEEHAVE. The corresponding R code and input file are available in two separate files, “S4-NePoFarm.R” and “S4-Input.txt”.

**Format of input data of daily forage supply for BEEHAVE**

To read in input data of daily forage supply in BEEHAVE, we created txt-files that provide information on the number of available patches, their distance to the colony, the detection probability, the amount of nectar and pollen they offer on each day of the year and the handling times to collect a nectar or a pollen load (Table S8. The BEEHAVE input file consists of 15 data columns (input variables) and 365 x (number of farmland patches) lines, defining the nectar and pollen availability of each farmland patch on each day of one year.

**Table S8:** Listed are the input variables of the BEEHAVE input file of the daily forage supply. The file consists of 365 lines, defining the nectar and pollen availability of each farmland patch on each day of one year and the time to gather a nectar or pollen load on this forage patch on each day of one year.

| **Input variable** | **Explanation** |
| --- | --- |
| day | day of year |
| ID | ID of the farmland patch |
| oldPatchID | ID of the farmland patch |
| patchType | Crop type of the farmland patch e.g. “oilseed rape” |
| distance_m | distance from the farmland patch to the hive (m) |
| xcor | x-coordinates of patch centre (used for visualization in 'foraging map' plot) |
| ycor | y-coordinates of patch centre (used for visualization in 'foraging map' plot) |
| size_sqm | size of the farmland patch (m²) (used for visualization in 'foraging map' plot) |
| quantityPollen_g | amount of available pollen today (g) |
| Concentration | sugar concentration of provided nectar (mol / l) |
| quantityNectar_l | amount of available nectar today (l) |
| calc_DetectProb | calculated detection probability that a scout finds the farmland patch (calculated on the basis of the size of the farmland patch and its distance to the hive determined in BEESCOUT model) |
| model_DetectProb | modelled detection probability that a scout finds the farmland patch (determined by an spatially explicit, individual based model, determined in BEESCOUT model) |
| NectarGathering_s | time to gather a nectar load on the farmland patch (s) |
| PollenGathering_s | time to gather a pollen load on the farmland patch (s) |

**Landscape structure and Scales**

Two different farmland structures located in Lincolnshire (Nocton Heath area, simple-structured landscape example) and Cornwall (Trenwheal, complex-structured landscape example) with an area of 5 km x 5 km in total are used as input. These farmlands (serving as landscape map templates to establish a broad range of different farmland scenarios) are represented by a grid of 171 (simple-structured farmland) and 1186 (complex-structured farmland) patches with different patch sizes.

A farmland patch is characterized by its location, patch size and crop species. Crop species determines its forage input provided to the bees (Table S9).

**Table S9: Attributes of state variable ‘farmland patch’ and its associated crop species.**

|  | **Name of variable** | **Explanation** | **Value** | **Unit** |
| --- | --- | --- | --- | --- |
| **Farmland patch** | patchID  distance  size_sqm  xcor  ycor | ID of the respective patch  Flight distance from the hive to the patch  Size of the patch  Patch location  Patch location | Patch-specific  Patch-specific  Patch-specific  Patch-specific  Patch-specific | -  m  m²  -  - |
| **Crop species** | | | | |
| Arable crops  Cultivated crops of forage cropping and permanent grassland  Bee pastures | patchType  quantityPollen_g  quantityNectar_l  Concentration  NectarGathering_s  PollenGathering_s | Crop species occurring on the patch  Amount of pollen provided by the patch per day  Amount of nectar provided by the patch per day  Sucrose concentration of the nectar provided by the patch  Time to fill a full crop volume at the patch  Time to collect a full pollen load at the patch | “Oilseed”, “Maize”, “Sunflower”, “Cereals”, “Bean”,  “Rye-grass”, “Rye-grass-dandelion”, “Rye-grass-white clover”,  “buckwheat”, “phacelia”  depending on crop species and patch size  depending on crop species and patch size  crop species-specific  crop species-specific  crop species-specific | -  -  -  g/m²  l/m²  mol/l  s  s |

**Procedures**

An overview of procedures of the landscape generator NePoFarm is presented in Figure S14.

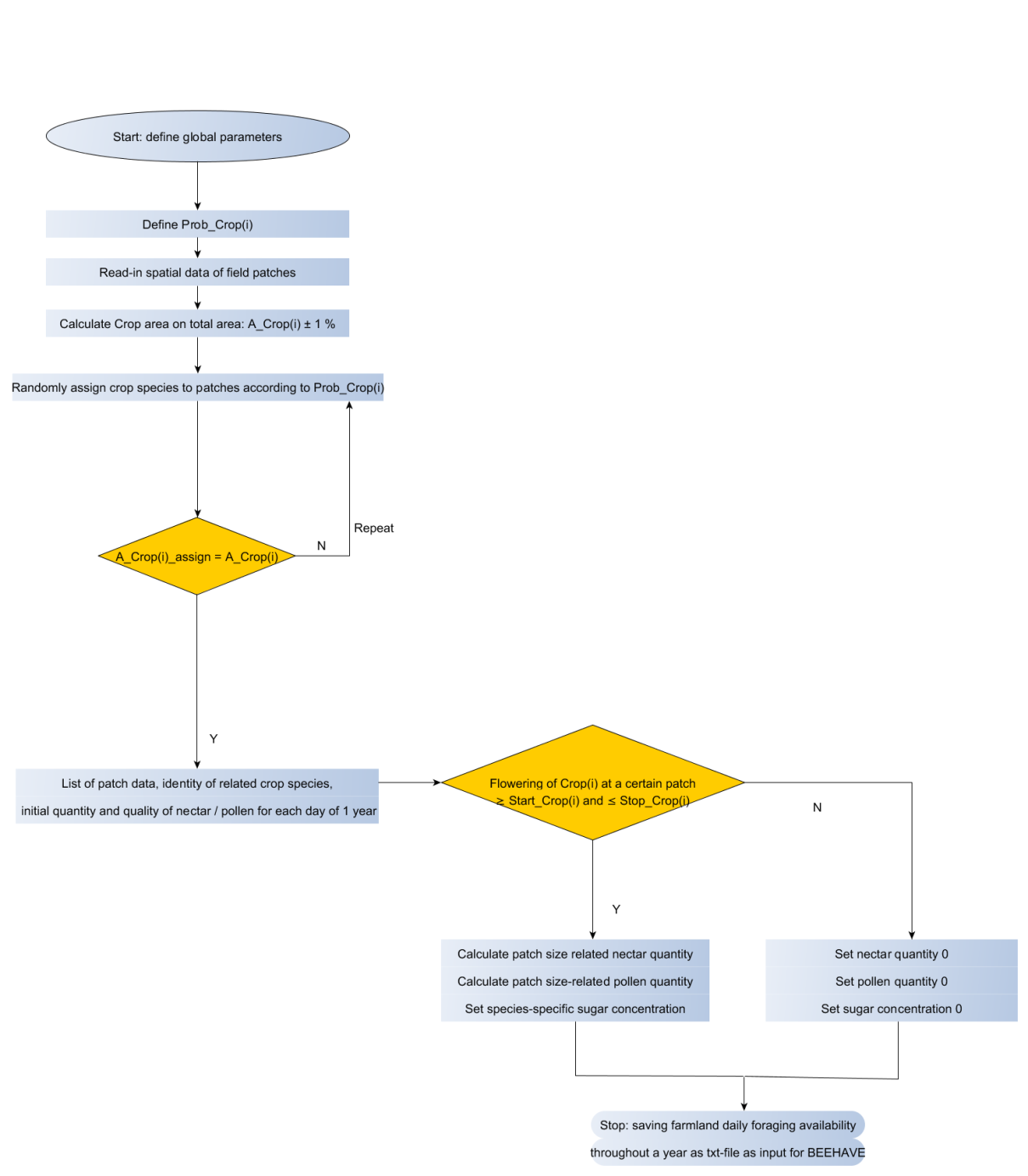

**Figure S14:** Overview of NePoFarm procedures.

*Procedure 1: Load attributes of farmland patches and define abundances of crop species* Attributes of farmland patches (patch size, patch location, distance of the patch to the hive, patch ID) saved in an external table as text file *data.row* were read in. At first, we initialized a vector *datall* for the patch attributes *patchType* containing 0 for each farmland patch (for simple-structured farmland: 171 patches) and created a data frame, which contains the patch attributes *ID*, *oldPatchID*, *distance_m*, *xcor*, *ycor*, *patchType*, *distance_m* and *size_sqm* for each farmland patch. Then, we calculated total agricultural area and range of crop species area according to its specific abundance ± 1 %. The body for the described procedure looks as follows:

Set pathway and read-in row data of patch characteristics.

*setwd("path")*

*data.row <- read.table("SimpleLandscape_Data.txt",header=T,sep="\t")*

Initialize vetor *datall* containing 0s as initial patch attribute *patchType* and create data frame containing spatial patch characteristics.

*datall <- rep(0,171)*

*ID <- c(0:170)*

*oldPatchID <- c(0:170)*

*patchType <- c(datall)*

*distance_m <- c(data.row$distance_m)*

*xcor <- c(data.row$xcor)*

*ycor <- c(data.row$ycor)*

*size_sqm <- c(data.row$size_sqm)*

*data.name <- data.frame(ID,oldPatchID,patchType,distance_m,xcor,ycor,size_sqm)*

Calculate total agricultural area.

*Total_AgroArea_sqm = sum(data.name$size_sqm)*

Define crop species abundance and calculate crop species area range depending on its specific abundance ± 1%.

*Prob_OSR =	0.25*

*Prob_Cereals =	0.25*

*Prob_Sunflower =	0.25*

*Prob_Maize =	0.25*

*Prob_Bean =	0.0*

*Prob_Rye =	0.0*

*Prob_Dandelion =	0.0*

*Prob_RyeClover =	0.0*

*OSR_Area_sqm = Total_AgroArea_sqm * Prob_OSR*

*OSR_Area_sqm_range_upper = OSR_Area_sqm + OSR_Area_sqm * 0.01*

*OSR_Area_sqm_range_under = OSR_Area_sqm - OSR_Area_sqm * 0.01*

*Procedure 2: Assign crop species randomly to farmland patches according to their abundance*

We draw random numbers from a uniform distribution to assign crop species to available farmland patches according to their given abundance in farmland area (Table S10) using the *sample*-instruction. Thereby, one has to specify the range of random numbers, the number of available patches (observations), the onset of repeated drawings of the same random number and the probability of the random numbers to be drawn from the uniform distribution. A *while* loop was used to repeat the drawing procedure of random numbers until all crop species occupy as much farmland area as given by their abundance. The *break* instruction within the *while* loop stops the procedure, when the final condition is reached (all crop species are assigned to available patches according to their abundance). So, the body of the *while*-procedure looks as follows:

*z <- 1*

*while(z > 0)*

*{*

*data.name$patchType<-sample(0:3,171,rep=TRUE,prob=c(Prob_OSR,Prob_Cereals,Prob_Maize,Prob_Sunflower))*

*OSR_Area <- sum (data.name$size_sqm[data.name$patchType == 0])*

*Cereals_Area <- sum (data.name$size_sqm[data.name$patchType == 1])*

*Maize_Area <- sum (data.name$size_sqm[data.name$patchType == 2])*

*Sunflower_Area <- sum (data.name$size_sqm[data.name$patchType == 3])*

*if (OSR_Area >= OSR_Area_sqm_range_under && OSR_Area <= OSR_Area_sqm_range_upper &&*

*Cereals_Area >= Cereals_Area_sqm_range_under && Cereals_Area <=*

*Cereals_Area_sqm_range_upper &&*

*Maize_Area >= Maize_Area_sqm_range_under && Maize_Area <= Maize_Area_sqm_range_upper &&*

*Sunflower_Area >= Sunflower_Area_sqm_range_under && Sunflower_Area <=*

*Sunflower_Area_sqm_range_upper)*

*{*

*break*

*}*

*}*

After this, we assigned to each random number the respective crop species (Table S9).

**Table S10: Crop types and their respective random numbers.**

| **Random number** | **Crop species** |
| --- | --- |
| 0 | Oilseed rape |
| 1 | Cereals |
| 2 | Sunflower |
| 3 | Maize |
| 4 | Bean |
| 5 | Rye-grass |
| 6 | Rye-grass-dandelion |
| 7 | Rye-grass-white clover |
| 8 | Buckwheat |
| 9 | Phacelia |

In the following, we defined vectors for each patch attribute *ID, oldPatchID, patchType, xcor, ycor, distance_m, size_sqm, quantityPollen_g*, *Concentration*, *quantityNectar_l*, *calc_DetectProb*, *model_DetectProb*, *NectarGathering_s* and *PollenGathering_s* and repeated each vector for each day of the year (365 times) using the *rep* instruction. In addition we add a spring feeder to the data frame and set *patchType* to “Boost”, *size_sqm* to ”10000” and *distance_m* to “1000”. The attributes *quantityPollen_g*, *Concentration*, *quantityNectar_l*, *calc_DetectProb*, *model_DetectProb*, *NectarGathering_s* and *PollenGathering_s* were set to their initial values 0, 0, 0, 0.2, 0.2, 1200, 600. Additionally, a column day was appended to the data frame that repeats day 1 to day 365 of the year for all farmland patches. The body of the appending procedure of all patch attributes to the data frame *data.name4* suitable as input file for the simulation model BEEHAVE looks as follows:

Create vectors describing all farmland patch characteristics for the BEEHAVE Input file and repeat for 365 times (each day of one year).

*n <- nrow(data.name)*

*ID <- rep(c(1:n-1,n),365)*

*oldPatchID <- rep(c(data.name$oldPatchID,400),365)*

*patchType <- rep(c(data.name$patchType,"Boost"),365)*

*xcor <- rep (c(data.name$xcor,1000),365)*

*ycor <- rep (c(data.name$ycor,1000),365)*

*size_sqm <- rep (c(data.name$size_sqm,10000),365)*

*distance_m <- rep (c(data.name$distance_m,1000),365)*

Write a data frame and arrange data frame according to ID

*data.name2 <-data.frame(ID,oldPatchID,patchType,distance_m,xcor,ycor,size_sqm)*

*data.name3 <- data.name2[with (data.name2, order(ID)),]*

Append columns *quantityPollen_g, Concentration, quantityNectar_l, calc_DetectProb, model_DetectProb, NectarGathering_s and PollenGathering*_s with initial values to data frame.

*v <- (n + 1) * 365*

*data.name3$quantityPollen_g <- rep (0,v)*

*data.name3$Concentration <- rep (0,v)*

*data.name3$quantityNectar_l <- rep (0,v)*

*data.name3$calc_DetectProb <- rep (0.2,v)*

*data.name3$model_DetectProb <- rep (0.2,v)*

*data.name3$NectarGathering_s <- rep (1200,v)*

*data.name3$PollenGathering_s <- rep (600,v)*

Append column day to data frame and write data frame with BEEHAVE suitable column names.

*data.name3$day <- rep(c(1:365),n+1)*

*data.name4 <- data.frame(data.name3$day,data.name3$ID,data.name3$oldPatchID,data.name3$patchType,data.name3$distance_m,data.name3$xcor,data.name3$ycor,data.name3$size_sqm,data.name3$quantityPollen_g,data.name3$Concentration,data.name3$quantityNectar_l,data.name3$calc_DetectProb,data.name3$model_DetectProb,data.name3$NectarGathering_s,data.name3$PollenGathering_s)*

*colnames(data.name4)<-c("day","ID","oldPatchID","patchType","distance_m","xcor","ycor","size_sqm","quantityPollen_g","Concentration","quantityNectar_l", "calc_DetectProb","model_DetectProb","NectarGathering_s","PollenGathering_s")*

Regarding that the initial values of *quantityPollen_g* *, Concentration* and *quantityNectar_l* were set to 0 and that we require the crop species-specific nectar and pollen quantities related to patch size (*size_sqm*) during species-specific flowering periods as input for simulations with BEEHAVE, we calculated the crop species-specific available amount of nectar and pollen to bees of each farmland patch as follows.

*Procedure 3: Amount of nectar and pollen of crop species depending on patch size*

At first, we calculated patch-size related amount of nectar and pollen of each crop species and the spring feeder *SpringBoost* using the species-specific amount of nectar and pollen per m² (initialized as global parameters e.g. *NectarOSR_l_sqm*, see below and Table S9) as follows (exemplary for oilseed rape (OSR)):

Calculate crop species-specific amount of nectar and pollen for each farmland patch depending on patch size.

*x_OSR <- data.name3$size_sqm [data.name3$patchType == "Oilseed"] * NectarOSR_l_sqm*

*y_OSR <- data.name3$size_sqm[data.name3$patchType=="Oilseed"] * PollenOSR_g_sqm*

Crop species-specific patch size-related nectar and pollen amounts and species-specific sugar concentrations were assigned to each patch within the species-specific flowering periods (initialized as global parameters e.g. *StartOSR* and *StopOSR,* see below and Table S8). Nectar and pollen amounts as well as sugar concentration before and after the species-specific flowering period were set to zero. The exemplary code for the patch size-related amount of nectar of oilseed rape during its flowering period is presented.

*data.name4$quantityNectar_l[data.name4$patchType=="Oilseed"]<-ifelse(data.name4$patchType*

*[data.name4$patchType=="Oilseed"]=="Oilseed" & data.name4$day[data.name4$patchType=="Oilseed"]*

*>=StartOSR & data.name4$day[data.name4$patchType=="Oilseed"]<=StopOSR,x_OSR,0)*

*Procedure 4: Output file*

Finally, the data table including all farmland patch attributes and crop species-specific amounts of nectar and pollen related to the patch size and sugar concentrations during their flowering periods was saved as a text file suitable as an input file for BEEHAVE using the instruction *write.table*.

**Initialization**

Several global parameters related to crop species abundance (e.g. *Prob_OSR*), flowering periods of crop species (e.g. *StartOSR* and *StopOSR*), crop species-specific amount of nectar and pollen per sqm² (e.g. *NectarOSR_l_sqm* and *PollenOSR_g_sqm*), crop species-specific sugar concentration (e.g. *ConcOSR*), handling times (*NectarGathering_s* and *PollenGathering_s*) and detection probabilities of patches (e.g. *calc_DetectProb*) were initialized and set to their specific values in NePoFarm (Table S11).

**Table S11: List of Parameters, default parameter set and parameter ranges.**

| **Parameter** | **Abbreviation** | **Value** | **Range** |
| --- | --- | --- | --- |
| ***Portion of crop species on available patches*** | | | |
| Proportion of oilseed rape patches on available patches | Prob_OSR | 0.25 | 0.0 -1.0 |
| Proportion of maize patches on available patches | Prob_Maize | 0.25 | 0.0 -1.0 |
| Proportion of cereals patches on available patches | Prob_Cereals | 0.25 | 0.0 -1.0 |
| Proportion of sunflower patches on available patches | Prob_Sunflower | 0.25 | 0.0 -1.0 |
| Proportion of field bean patches on available patches | Prob_Bean | 0.0 | 0.0 -1.0 |
| Proportion of rye-grass -white clover patches on available patches | Prob_Clover | 0.0 | 0.0 -1.0 |
| Proportion of rye-grass patches on available patches | Prob_Rye | 0.0 | 0.0 -1.0 |
| Proportion of rye-grass-dandelion patches on available patches | Prob_Dandelion | 0.0 | 0.0 -1.0 |
| Proportion of buckwheat patches on available patches | Prob_Buckwheat | 0.0 | 0.0 -1.0 |
| Proportion of phacelia patches on available patches | Prob_Phacelia | 0.0 | 0.0 -1.0 |
| ***Flowering period of crop species in day (start and end date of flowering)*** | | | |
| Flowering period of oilseed rape | StartOSR  StopOSR | 114  144 | - |
| Flowering period of maize | StartMaize  StopMaize | 197  210 | - |
| Flowering period of sunflower | StartSunflower  StopSunflower | 237  264 | - |
| Flowering period of field bean | StartBean  StopBean | 153  182 | - |
| Flowering period of white clover | StartClover  StopClover | 140  242 | - |
| Flowering period of dandelion | StartDandelion  StopDandelion | 92  145 | - |
| Flowering period of buckwheat | StartBuckwheat  StopBuckwheat | 172  260 | - |
| Flowering period of phacelia | StartPhacelia_1  StopPhacelia_1  StartPhacelia_2  StopPhacelia_2 | 196  255  265  301 | - |
| A feeder at one nearby patch (1000 m) and 1 ha size offering low amounts of nectar and pollen in early spring | Boost_Period | 90 | - |
| ***Nectar amount per crop species in l per m²*** | | | |
| Nectar amount of oilseed rape | NectarOSR_l_sqm | 0.001 | - |
| Nectar amount of maize | NectarMaize_l_sqm | 0 | - |
| Nectar amount of sunflower | NectarSunflower_l_sqm | 0.000008 | - |
| Nectar amount of cereals | NectarCereals_l_sqm | 0 | - |
| Nectar amount of field bean | NectarBean_l_sqm | 0.0006 | - |
| Nectar amount of white clover | NectarClover_l_sqm | 0.0013 | - |
| Nectar amount of dandelion | NectarDandelion_l_sqm | 0.000049 | - |
| Nectar amount of buckwheat | NectarBuckwheat_l_sqm | 0.0072 | - |
| Nectar amount of phacelia | NectarPhacelia_l_sqm_1  NectarPhacelia_l_sqm_2 | 0.00059  0.000443 | - |
| Nectar amount of feeder for total size of the feeder patch (1 ha) | BoostNectar | 1 | - |
| ***Pollen amount per crop species in g per m²*** | | | |
| Pollen amount of oilseed rape | PollenOSR_g_sqm | 0.349 | - |
| Pollen amount of maize | PollenMaize_g_sqm | 8.036 | - |
| Pollen amount of sunflower | PollenSunflower_g_sqm | 0.18 | - |
| Pollen amount of cereals | PollenCereals_g_sqm | 0 | - |
| Pollen amount of field bean | PollenBean_g_sqm | 0.0945 | - |
| Pollen amount of white clover | PollenClover_g_sqm | 0.0141 | - |
| Pollen amount of dandelion | PollenDandelion _g_sqm | 0.011 | - |
| Pollen amount of buckwheat | PollenBuckwheat_g_sqm | 3.6 | - |
| Pollen amount of phacelia | PollenPhacelia _g_sqm_1  PollenPhacelia _g_sqm_2 | 1.2  0.9 | - |
| Pollen amount of feeder for total size of the feeder patch (1 ha) | BoostPollen | 500 | - |
| ***Sugar concentration of crop species in mol per l*** | | | |
| Sugar concentration of oilseed rape | ConcOSR | 1.7 | - |
| Sugar concentration of maize | ConcMaize | 0.0 | - |
| Sugar concentration of sunflower | ConcSunflower | 0.7 | - |
| Sugar concentration of cereals | ConcCereals | 0.0 | - |
| Sugar concentration of field bean | ConcBean | 1.46 | - |
| Sugar concentration of white clover | ConcClover | 1.08 | - |
| Sugar concentration of dandelion | ConcDandelion | 1.19 | - |
| Sugar concentration of buckwheat | ConcBuckwheat | 1.17 | - |
| Sugar concentration of phacelia | ConcPhacelia | 1.05 | - |
| Sugar concentration of the feeder | BoostConc | 1.5 | - |
| ***Time to collect a full nectar or pollen load in s*** | | | |
| Time to fill up a full crop volume of nectar (50 µl) at a patch in s | NectarGathering_s | 1200 | - |
| Time to collect a full pollen load (0.015 g) at a patch in s | PollenGathering_s | 600 | - |
| ***Detection probability of available patches*** | | | |
| Calculated detection probability | calc_DetectProb | Constant (0.2) | BeeScout calculated |
| Modelled detection probability | model_DetectProb | Constant (0.2) | BeeScout calculated |

### S5 P_H_ / P_P_: Indices for comparing forage-induced stress at the colony level

To compare the effects of forage availability determined by the spatiotemporal nectar and pollen supply of the different farmland scenarios at the colony level, we calculated new indices: *P_H_* (honey-to-worker-productivity), and *P_P_* (pollen-to-worker-productivity). They evaluate how much nectar and pollen provided by a certain cropping system could be converted into the production and maintenance of the colony’s work force in terms of worker bees over the colony’s annual cycle_._ They quantify, cumulated over five years or over the time to extinction (*TTE*, see above), the amount of honey and pollen in kilogram which the colony used for actually producing and maintaining adult worker bees. Values of *P_H_* and *P_P_* were determined for each scenario by using the total number of adult worker bees produced by the colony over the entire simulation, or until extinction, *N*_Workers_, multiplied by the honey or pollen requirement, respectively, of a worker bee over its complete larval and adult life span. In these two indices, we ignored the production of queens and drones and their corresponding food requirements, as they do not directly contribute to the colony’s work force.

The corresponding parameter values and equations for calculations of a bee’s food requirements are listed in Table S13.

**Table S13:** Parameters and equations for calculation of the indices *P_H_* and *P_P_* which quantify, over simulation time or the time to extinction (*TTE*), how much nectar and pollen provided by a certain cropping system could be converted into the production and maintenance of the colony’s force of worker bees over the colony’s annual cycle

| Formula / Parameter | Explanation | Value | Unit | Reference |
| --- | --- | --- | --- | --- |
| M_Pollen_Larva_ | Required pollen amount to rear a single worker larva | 142 | mg | Seeley 1985, 1995  Hrassnigg & Crailsheim 2005 |
| M_Pollen_Worker_ | Daily required pollen amount of a single adult worker bee | 1.5 | mg d^-1^ | Crailsheim et al. 1992, Rortais et al. 2005, |
| T_Life_AdultWorker_ | Lifespan of an adult worker bee on average | 38 | d | Rortais et al. 2005, |
| M_PollenConsumption_Worker_ = M_Pollen_Worker_ * T_Life_AdultWorker_ | Pollen requirement of a single adult worker bee over lifespan | 57 | mg | Calculated from literature data |
| M_TotalPollen_ =  M_Pollen_Larva_ + M_PollenConsumption_Worker_ | Total pollen requirement of a single worker bee during its larval stage and adult stage | 0.000199 | kg | Calculated from literature data |
| M_Honey_Larva_ | Required honey amount to rear a single worker larva | 59.4 | mg | Rortais et al. 2005 |
| M_Honey_Worker_ | Daily required honey amount of by a single adult worker bee averaged over all activity categories of the bees, which did not include brood rearing and flight activities, but lower consumption during winter times cancels highly energy-consuming activities out | 11.47 | mg d^-1^ | Huang et al. 1998,  Rortais et al. 2005 |
| M_HoneyConsumption_Worker_ =  M_Honey_Worker_ * T_Life_AdultWorker_ | Honey requirement of a single adult worker bee over lifespan | 435.86 | mg | Calculated from literature data |
| M_TotalHoney_ =  M_Honey_Larva_ + M_HoneyConsumption_Worker_ | Total honey requirement of a worker bee during its larval and adult stage | 0.000495 | kg | Calculated from literature data |
| Cumul_Workers_ | Cumulative number of workers summed up over simulation time | - | - | calculated for each scenario in BEEHAVE |
| N_Workers_ =  Cumul_Workers_ / T_Life_AdultWorker_ | Number of all worker bees ever produced in the colony over simulation time | - | - | calculated for each scenario |
| P_H_ = *N*_workers_ * M_TotalHoney_ | Honey-to-colony’s worker productivity | - | kg | calculated for each scenario |
| P_P_ = *N*_workers_ * M_TotalPollen_ | Pollen-to-colony’s worker productivity | - | kg | calculated for each scenario |

### S6 Ranking farmland factors by mimicking a local sensitivity analysis procedure

**Ranking farmland factors: local sensitivity analyses**

To qualitatively rank farmland factors by their importance for colony performance, we applied a procedure mimicking a local sensitivity analysis by varying only one factor at a time while the others were fixed at their nominal values (Saltelli et al. 2000). For example, to assess the importance of farmland structure and the importance of crop species number, we let this variable vary over its possible values (simple- and complex-structured) while keeping all other variables fixed (for farmland structure: explanatory variables fixed at number of crop species = 4, crop species abundance = 5, crop species identity = oilseed rape and farmland structure varies; for number crop species: explanatory variables fixed at crop species abundance = 5, farmland structure = simple, crop species identity = oilseed rape and number of crop species varies). We then fitted a series of univariate linear models with the variable that was allowed to vary (i.e. farmland structure) used as an explanatory and *TTE*, *P_H_*, and *P_P_* as response variables. The adjusted R^2^ from these models were used to quantify the variation explained by the variable that was varied, and to qualitatively rank the contribution farmland variables made to explain the response variables. To satisfy the model diagnostics we log-transformed all response variables except *TTE*. *SurvProb* was omitted from analyses, because only 4.7 % of 15,523 data points resulted in success (i.e. colony survival over five years), making maximum likelihood estimation difficult and unstable (King & Zeng 2001). All analyses were conducted with R version 3.1.2 (R Development Core Team 2014).

### S7 Simulations - Landscape settings

**Sensitivity analysis of nectar and pollen data**

We conducted sensitivity analyses of nectar and pollen data under hypothetical landscape settings consisting of one single crop patch located in 500 m flight distance to the hive and constant nectar and pollen flow of one single crop throughout the year. To account for different nectar and pollen amounts we assigned either oilseed rape (providing 0.001 l nectar and 0.349 g pollen per m²) or sunflower (providing 0.000008 l nectar and 0.18 g pollen per m²) as exemplary high- or low-rewarding crops to this hypothetical landscape and varied patch sizes from 1.1 ha p to 69 ha. Moreover, to account for differences in nectar quality we varied sugar concentrations from 0.5 up to 2.3 mol/ l (17 – 80 %) in increments of 0.1 as shown in Table S14. Sensitivity analyses were run for five simulation years and replicated for 30 times per sensitivity scenario. We analysed colony performance in terms of time to colony extinction (TTE), colony’s productivity of force of worker bees over simulation time or until the colony became extinct (N_Workers_), colony size and honey stores at the end of the fifth simulation year (only for surviving colonies).

**Table S14:** Design of sensitivity analyses of nectar and pollen data under hypothetical landscape settings.

| Patch size (ha) | Distance  (m) | Max nectar  (OSR l / ha) | Max pollen  (OSR kg / ha) | Min nectar  (SF l / ha) | Min pollen  (SF kg / ha) | Sugar concentration (mol / l) |
| --- | --- | --- | --- | --- | --- | --- |
| 69 | 500 | 690 | 240.8 | 5.52 | 124.2 | 0.5 – 2.3 |
| 10 | 500 | 100 | 34.9 | 0.8 | 18 | 0.5 – 2.3 |
| 4 | 500 | 40 | 14 | 0.32 | 7.2 | 0.5 – 2.3 |
| 1.1 | 500 | 11 | 3.84 | 0.09 | 1.98 | 0.5 – 2.3 |

As shown in Figs. S15 – S17 the exemplary high-rewarding crop species oilseed rape was most sensitive to sugar concentrations in all output variables, whereas the exemplary low-rewarding crop species sunflower was most sensitive to sugar concentration and patch size. For oilseed rape a sugar concentration of 1.2 mol / l was sufficient to ensure survival of the 5 simulation years independent from patch size representing nectar and pollen quantities offered at the patch. So, for a 1.1 ha patch, oilseed rape produced 11 l nectar and 3.84 kg pollen per day. For sunflower, a minimum sugar concentration of 1.5 mol / l and a minimum patch size of 69 ha was required to ensure survival of the modelled honeybee colony. For a 1.1 ha patch, sunflower produces 0.09 l nectar and 1.98 kg pollen. For a 69 ha patch, sunflower produces 5.52 l nectar and 124.2 kg pollen per day.

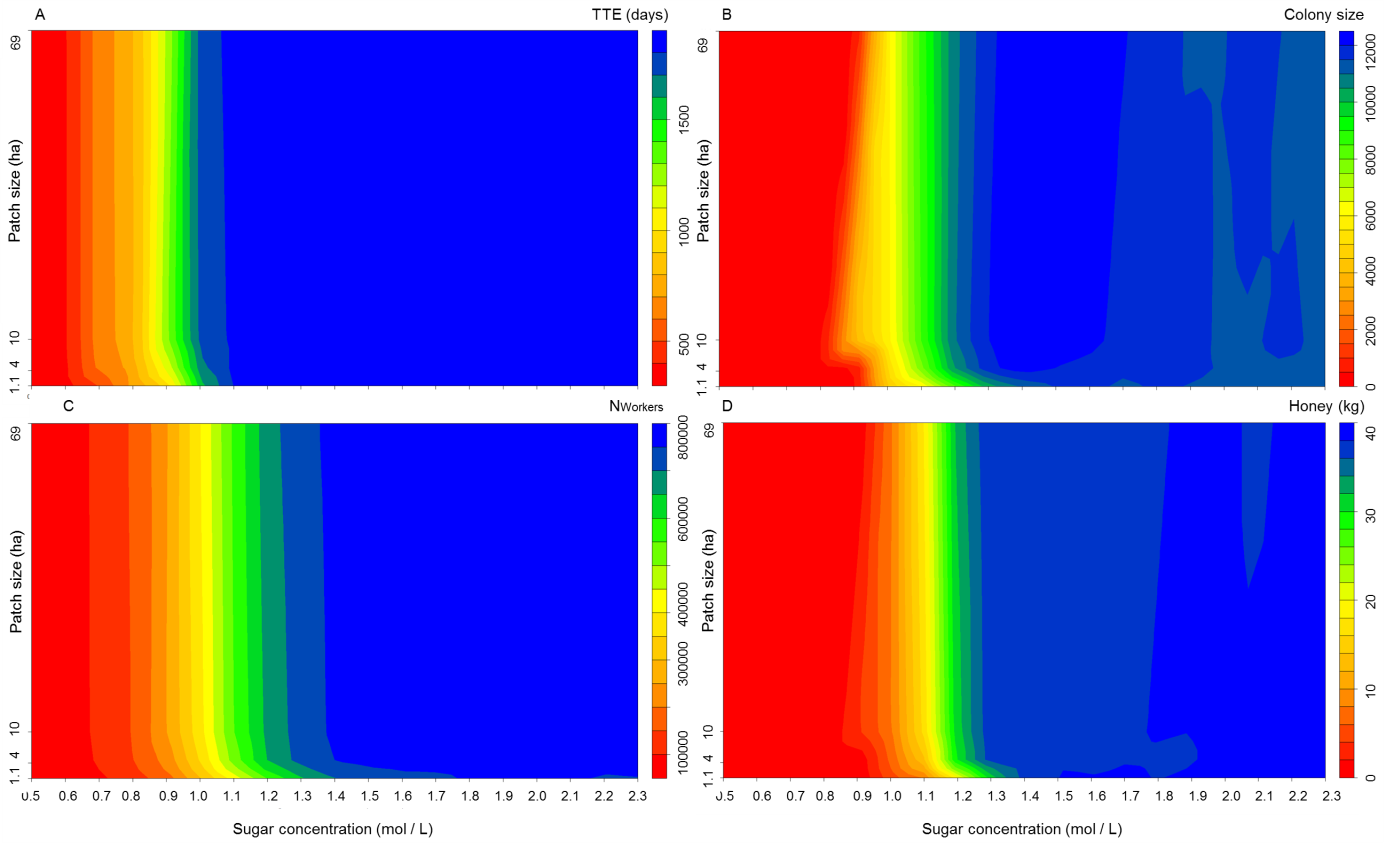
**Figure S15:** Sensitivity of nectar and pollen data of oilseed rape (exemplary high-rewarding crop). Sensitivity in colony performance in terms of **(A)** time to colony extinction (given in days), **(B)** colony size at the end of the fifth year of simulation time (Colony size), **(C)** colony’s productivity of force of worker bees over simulation time of five years or until the colony became extinct (N_Workers_), and **(D)** colony’s honey stores at the end of the fifth year of simulation (given in kg), in response to patch size increasing from 1.1 ha up to 69 ha and sugar concentration ranging from 0.5 mol/l up to 2.3 mol/l was analysed.

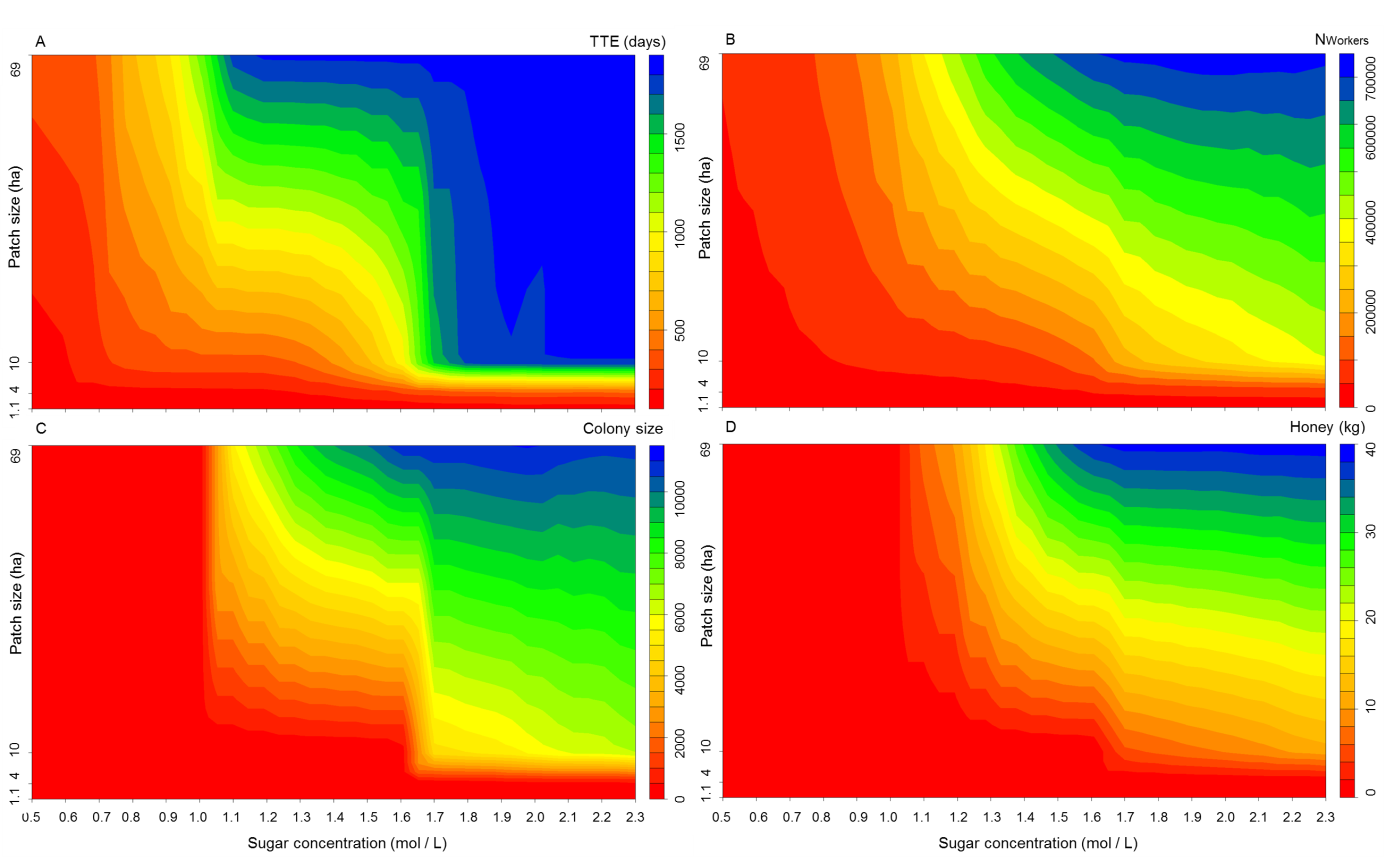
**Figure S16:** Sensitivity of nectar and pollen data of sunflower (exemplary low-rewarding crop). Sensitivity in colony performance in terms of **(A)** time to colony extinction (given in days), **(B)** colony’s productivity of force of worker bees over simulation time of five years or until the colony became extinct (N_Workers_), **(C)** colony size at the end of the fifth year of simulation time (Colony size), and **(D)** colony’s honey stores at the end of the fifth year of simulation (given in kg), in response to patch size increasing from 1.1 ha up to 69 ha and sugar concentration ranging from 0.5 mol/l up to 2.3 mol/l was analysed.

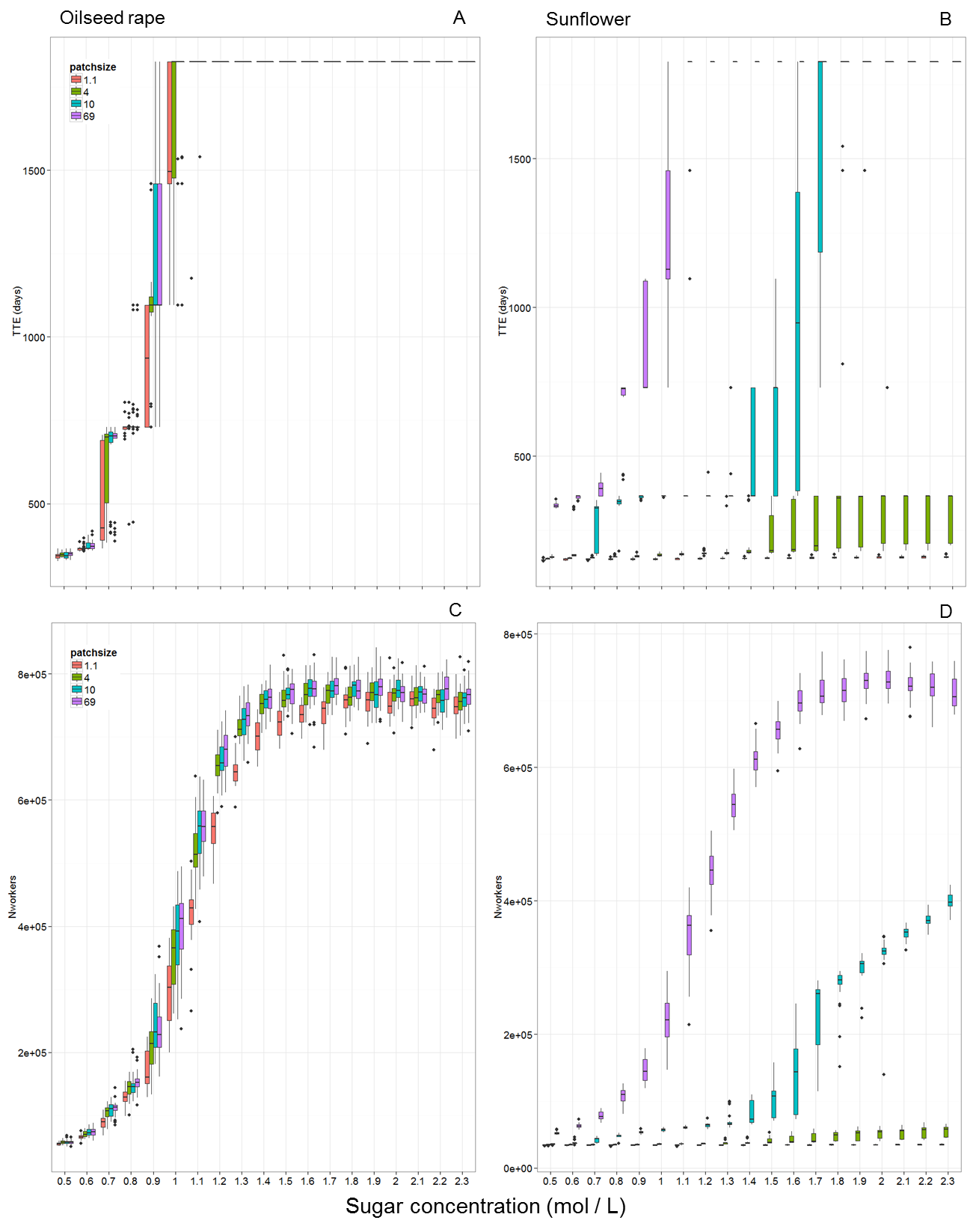
**Figure S17:** Effects of increasing patch size and sugar concentration of high-rewarding oilseed rape **(A, C)** and low-rewarding sunflower **(B, D)** on **(A, B)** time to colony extinction (TTE, given in days) and **(C, B)** colony’s productivity of force of worker bees over simulation time of five years or until the colony became extinct (N_Workers_) under hypothetical landscape settings. Patch sizes are illustrated by colored bars ranging from 1.1 ha up to 69 ha.

**Distances**

To analyse the effects of foraging distance on colony performance in our model world, we run simulations in hypothetical landscape consisting of one single forage patch of 69 ha size (maximum patch size in simple-structured landscape). Furthermore, we assigned the highest rewarding crop of our study, oilseed rape (maximum sugar concentration with 1.7 mol/l and rich nectar and pollen amounts with 0.001 l and 0.349 g per m²), to this forage patch. In this hypothetical landscape setting oilseed rape flowers during the whole year and provides constant nectar and pollen rewards every day of the year. To test distance effects from the hive to the forage patch, we ran simulations for foraging distances ranging from 0.1 m up to 3500 m in increments of 100 m. Each distance scenarios ran for 5 simulation year and 30 replicates, and colony performance in terms of survival probability (SurvProb), time to colony extinction (TTE), colony’s productivity of force of worker bees over simulation time or until the colony became extinct (N_Workers_), colony size and honey stores at the end of the fifth simulation year (only for surviving colonies) was analysed.

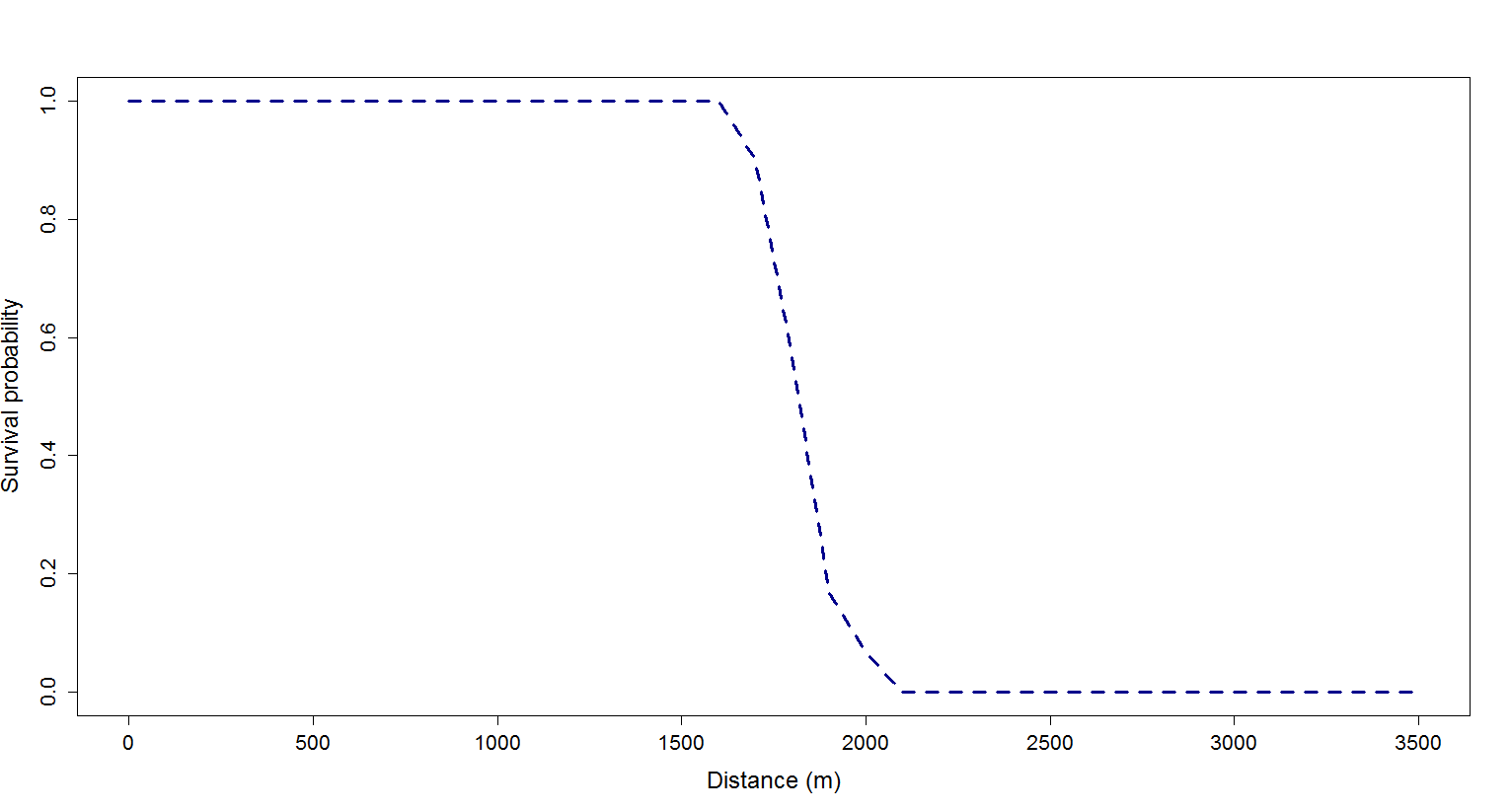

**Figure S18:** Survival probability with increasing distance (given in metres) under hypothetical landscape settings (one single forage patch of the highest rewarding crop oilseed rape).

From 1500 m foraging distance to the single high-rewarding patch onwards honey stores at the end of the five simulation years is distinctly reduced compared to the 1000 m distance scenario. From 2000 metere foraging distance all colonies went extinct within the third or fourth year at latest, because energy requirements of the colony are no longer sufficiently met by the long-distant foraging.

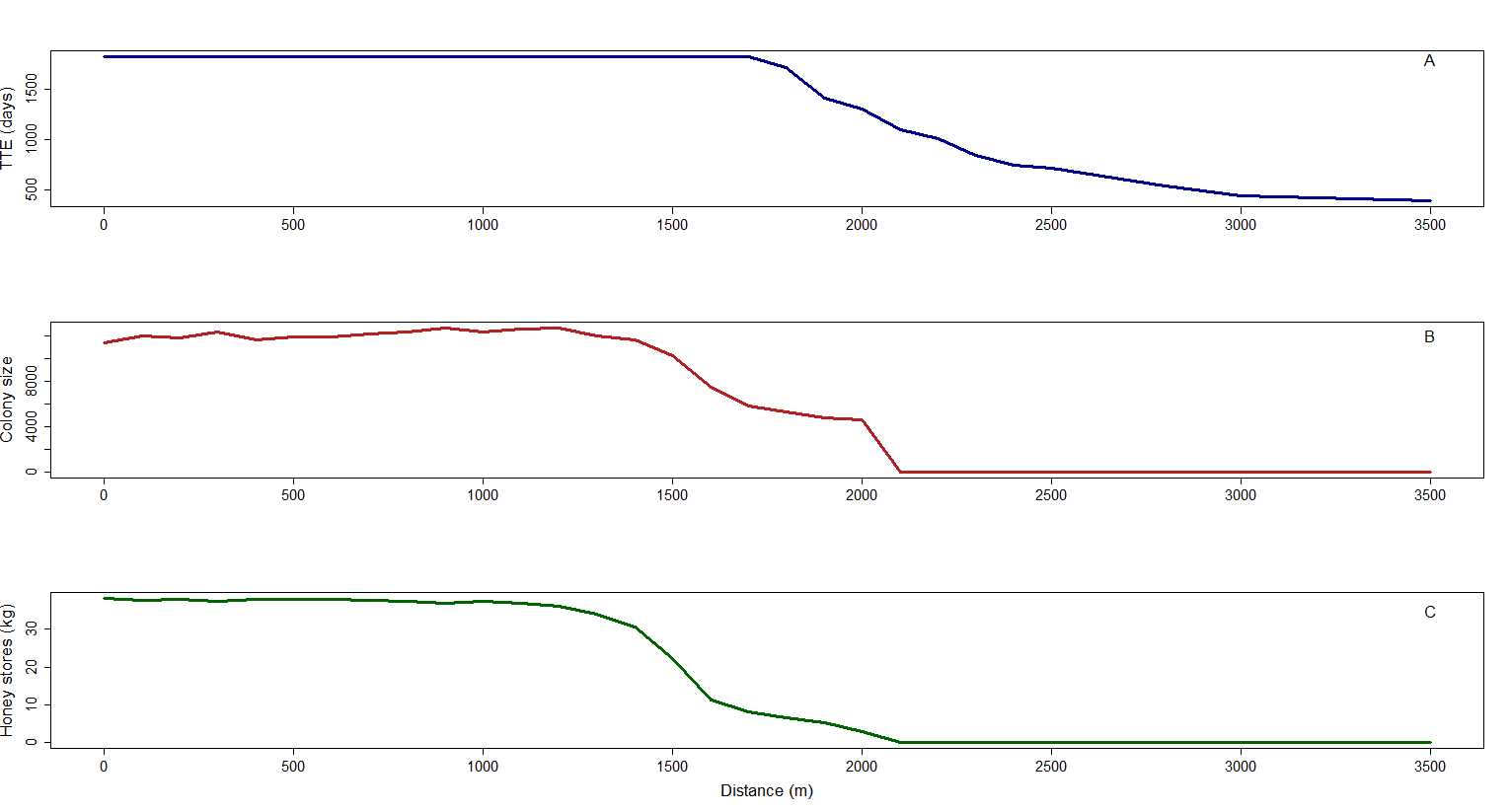

**Figure S19:** Time to colony extinction (TTE, given in days – upper panel), colony size (middle panel) and honey stores (given in kg - lower panel) at the end of the fifth simulation year with increasing distance (given in metres) under hypothetical landscape settings (one single forage patch of the highest rewarding crop oilseed rape).

**Table S15:** Effects of increasing foraging distance on colony performance in hypothetical landscapes consisting of the highest rewarding crop oilseed rape (one single patch). For all foraging distances survival probability (SurvProb), time to colony extinction (TTE, given in days), colony size (at the end of the fifth year of simulation, only for surviving colonies), and honey stores (at the end of the fifth year, only for surviving colonies) are presented.

| Distance (m) | SurvProb | TTE (days) | Colony size | Honey (kg) |
| --- | --- | --- | --- | --- |
| 0.1 | 1 | 1825 | 11325 | 38.2 |
| 100 | 1 | 1825 | 119505 | 37.7 |
| 200 | 1 | 1825 | 118035 | 37.9 |
| 300 | 1 | 1825 | 122745 | 37.3 |
| 400 | 1 | 1825 | 116365 | 38 |
| 500 | 1 | 1825 | 118655 | 37.9 |
| 600 | 1 | 1825 | 11904 | 37.8 |
| 700 | 1 | 1825 | 12112 | 37.7 |
| 800 | 1 | 1825 | 12293 | 37.5 |
| 900 | 1 | 1825 | 12685 | 37 |
| 1000 | 1 | 1825 | 12300 | 37.3 |
| 1100 | 1 | 1825 | 12558 | 36.9 |
| 1200 | 1 | 1825 | 12675 | 36 |
| 1300 | 1 | 1825 | 11963 | 34.1 |
| 1400 | 1 | 1825 | 11647 | 30.6 |
| 1500 | 1 | 1825 | 10186 | 22.3 |
| 1600 | 1 | 1825 | 7446 | 11.2 |
| 1700 | 0.9 | 1825 | 5822 | 8 |
| 1800 | 0.57 | 1717.3 | 5312 | 6.5 |
| 1900 | 0.17 | 1418.1 | 4736 | 5.2 |
| 2000 | 0.067 | 1300 | 4582 | 2.9 |
| 2100 | 0 | 1095.6 | 0 | 0 |
| 2200 | 0 | 1005.8 | 0 | 0 |
| 2300 | 0 | 839.7 | 0 | 0 |
| 2400 | 0 | 738.8 | 0 | 0 |
| 2500 | 0 | 711.6 | 0 | 0 |
| 2800 | 0 | 531.4 | 0 | 0 |
| 3000 | 0 | 432.2 | 0 | 0 |
| 3500 | 0 | 383 | 0 | 0 |

**Replicates**

We conducted simulations to test the sensitivity of our metrices of colony performance to the number of model replicates. For this, we chose scenarios of semi-realistic landscape settings with a survival probability of 50 % reflecting highest uncertainty in the outcome. For 4 crops cropping system scenario the composition of ryegrass-dandelion pasture, ryegrass-clover pasture, maize and 85 % phacelia (all other crops had 5 % abundance) showed highest uncertainty in colony performance. For an 8 crops cropping system senario the composition of oilseed rape, maize, 65 % sunflower, bean, ryegrass-dandelion pasture, ryegrass-clover pasture, buckwheat and phacelia (all other crops had a 5 % abundance) resulted in highest varying outcomes of colony performance. Both semi-realistic scenarios were run for five simulation years and replicated for 1 up to 100 times. We analysed colony performance in terms of time to colony extinction (TTE) and colony’s productivity of force of worker bees over simulation time or until the colony became extinct (N_Workers_).

4 crops

TTE (days)

NWorkers

| No Replicates | mean_TTE | mean_NWorkers |
| --- | --- | --- |
| 1 | 1825 | 444593 |
| 5 | 1679 | 382351 |
| 10 | 1606 | 367924 |
| 15 | 1655 | 362981 |
| 20 | 1697.2 | 373335 |
| 25 | 1709.2 | 379513 |
| 30 | 1715.5 | 379060 |
| 35 | 1689 | 370036 |
| 40 | 1671.7 | 364128 |
| 50 | 1673.2 | 360245 |
| 60 | 1686.3 | 361140 |
| 70 | 1685.3 | 361542 |
| 80 | 1692 | 364741 |
| 90 | 1688 | 359154 |
| 100 | 1687 | 359617 |

**Figure S20:** Sensitivity of time to colony extinction (TTE, given in days) and colony’s productivity of force of worker bees over simulation time of five years or until the colony became extinct (N_Workers_) to the number of model replicates for the semi-realistic cropping systems composed of ryegrass-dandelion pasture, ryegrass-clover pasture, maize and 85 % phacelia (all other crops had 5 % abundance).

Our preliminary sensitivity test for the response variables representing colony viability to the number of replicates for two exemplary scenarios of a 4 crop and an 8 crop cropping system scenario indicated that 30 replicates per scenario were sufficient to reduce variation in the resulting outcomes and did not artificially increase the significance levels by increasing the number of simulations (see Figs. S20 and S21).

8 crops

Averaged Survival time (says)

Averaged NWorkers

| No Replicates | mean_TTE |
| --- | --- |
| 1 | 1825 |
| 5 | 1752 |
| 10 | 1686.9 |
| 15 | 1708.6 |
| 20 | 1707.1 |
| 25 | 1726 |
| 30 | 1730.3 |
| 35 | 1724.8 |
| 40 | 1737.3 |
| 50 | 1725.7 |
| 60 | 1724 |
| 70 | 1723.9 |
| 80 | 1718.3 |
| 90 | 1709.9 |
| 100 | 1700.2 |

|  | mean_NWorkers |
| --- | --- |
| 1 | 370683 |
| 5 | 389893 |
| 10 | 376801 |
| 15 | 362145 |
| 20 | 356926 |
| 25 | 366078 |
| 30 | 366618 |
| 35 | 363122 |
| 40 | 372337 |
| 50 | 371768 |
| 60 | 377459 |
| 70 | 379395 |
| 80 | 379520 |
| 90 | 380428 |
| 100 | 380656 |

**Figure S21:** Sensitivity of time to colony extinction (TTE, given in days) and colony’s productivity of force of worker bees over simulation time of five years or until the colony became extinct (N_Workers_) to the number of model replicates for the semi-realistic cropping system composed of oilseed rape, maize, 65 % sunflower, bean, ryegrass-dandelion pasture, ryegrass-clover pasture, buckwheat and phacelia (all other crops had a 5 % abundance).

**Distance and patch size**

To analyse the effects of patch size and foraging distance of different cropping systems on colony performance, we performed simulations of hypothetical landscapes for exemplary monocultures, 3 crop, 4 crop and 8 crop cropping system scenarios. In these hypothetical landscapes each crop species occupies one single patch of 69 ha (maximum patch size of simple-structured landscape), 10 ha (mean patch size of simple-structured landscape), 1.1 ha (mean patch size of complex-structured landscape) or 0.06 ha (minimum patch size of complex-structured landscape; see table S16). For all patch sizes we analysed foraging distances from the hive to the crop patch of 0.1 m, 500 m or 1000 m. To represent early spring foraging on days when weather conditions are suitable for foraging flights, we placed a forage patch providing low daily amounts of nectar (1 l) and pollen (0.5 kg) from January until end-March at 0.1 m from the hive, which allows the colony to survive the early spring period (see also Horn et al. 2016). This forage patch represents early flowering plants such as snowdrop (*Galanthus nivalis*), crocus (*Crocus spec*.) and willow trees (*Salix spec*.). The available nectar and pollen amounts at this forage patch can be completely depleted on a given day, so the time a forager needs to collect a full nectar or pollen load (handling time) increases with the degree of forage depletion at this patch. We ran simulations for 5 years and for 30 replicates per scenario, and analysed colony performance in terms of survival probability (SurvProb), time to colony extinction (TTE), colony’s productivity of force of worker bees over simulation time or until the colony became extinct (N_Workers_), colony size and honey stores at the end of the fifth simulation year (only for surviving colonies).

**Table S16:** Patch size distributions in the semi-realistic landscapes.

| Landscape structure | Patch size (ha) | | | # patches | | | | |
| --- | --- | --- | --- | --- | --- | --- | --- | --- |
|  | mean ± sd | min | max | Total | ≤ 1ha | >1ha and ≤ 4ha | > 4ha and ≤ 10ha | > 10ha |
| Simple | 10.82 ± 10.24 | 1.11 | 69 | 171 | 0 | 40 | 61 | 70 |
| complex | 1.1 ± 1.54 | 0.06 | 13.3 | 1186 | 787 | 336 | 58 | 5 |

**Monocultures**

As exemplary monocultures we analysed the effects of increasing patch size on colony performance for an oilseed rape monoculture and a clover monoculture in a hypothetical landscape, whereby the single patch of the monoculture was located immediately to the hive (0.1 m distance). SurvProb, TTE and NWorkers were similar for all patch sizes of the artificial landscape and for the semi-realistic landscape of the respective monoculture (Table S17). So, both increasing patch size determining the amount of delivered nectar and pollen over the crop’s flowering period and short flight distance determining the low energy level spent for the foraging trip to the nectar and pollen resource are not able to compensate for the following long-lasting gaps in nectar and pollen supply within this monoculture landscape.

**Table S17:** Effects of increasing patch size on colony performance in artificial monoculture landscapes (one single patch of monoculture located in 0.1 m distance to the hive). For all patch sizes of monocultures survival probability (SurvProb), time to colony extinction (TTE, given in days), and number of produced worker bees until colony became extinct (N_Workers_) are presented.

| Monoculture | Patch size (ha) | SurvProb | TTE (days) | N_Workers_ |
| --- | --- | --- | --- | --- |
| OSR | 69 | 0 | 254.2 | 34145 |
|  | 10 | 0 | 252.9 | 34030 |
|  | 1.1 | 0 | 254.4 | 33949 |
|  | 0.06 | 0 | 253.4 | 33120 |
|  | Semi-realistic landscape | 0 | 251.4 | 32504 |
| Clover | 69 | 0 | 343.4 | 41166 |
|  | 10 | 0 | 342.1 | 41141 |
|  | 1.1 | 0 | 346.6 | 40900 |
|  | 0.06 | 0 | 365 | 29161 |
|  | Semi-realistic landscape | 0 | 343.5 | 39942 |

**3 crop cropping system scenarios**

As exemplary scenarios in artificial landscapes we analysed the effects of increasing patch size and increasing foraging distance for the cropping system composed of oilseed rape, ryegrass-clover and phacelia, which resulted in 20 % survival under our semi-realistic landscape settings. As an exemplary cropping system without any surviving colonies under the semi-realistic landscape settings we chose the crop composition of oilseed rape, sunflower and ryegrass-clover.

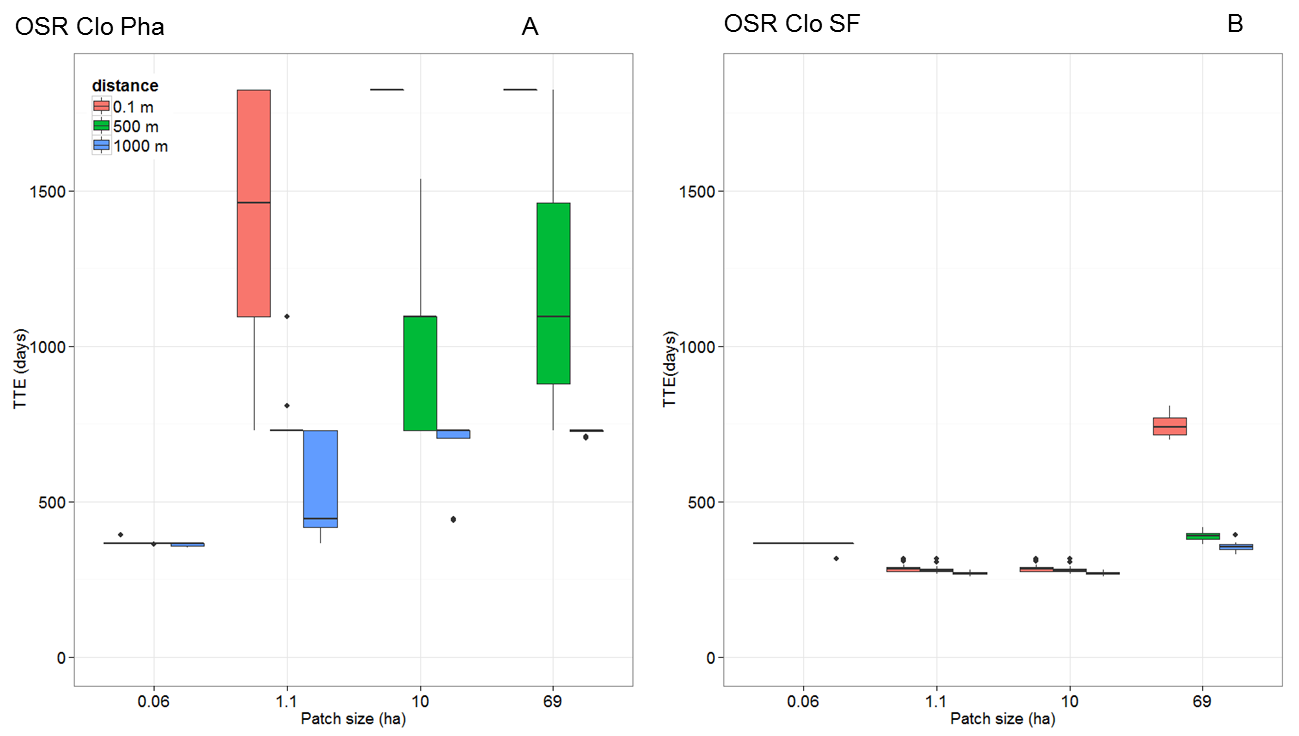

**Figure S22**: Effects of increasing patch size and foraging distance on time to colony extinction (TTE, given in days, y-axis) for 3 crop cropping systems (A) oilseed rape, ryegrass-clover, phacelia (OSR Clo Pha) and (B) oilseed rape, ryegrass-clover, sunflower (OSR Clo SF) in a hypothetical landscape (one single patch per crop species). Patch sizes are given in ha (x-axis) and foraging distances are illustrated by colored bars.

For three crop cropping systems consisting of oilseed rape, phacelia and ryegrass-clover pasture even for biggest patch size of 69 ha and shortest distance of 0.1 m for each of these crops colonies survived over the simulation time of years but colony size and honey stores were very low (12.4 kg honey). If distance increases up to 500 m, patch size doesn’t matter anymore. Even for the shortest foraging distance one hectare of each crop species isn’t sufficient to fulfill colony’s energy requirements. As under the semi-realistic landscape settings most of the large-sized patches of the high-rewarding crops oilseed rape and phacelia (> 10 ha) were located in more than 500 metres flight distance to the hive (Tab. S19), the energy expenditure is not sufficient in order to buffer the low forage supply (1.43 litre nectar and 0.141 kg pollen per hectare) offered by rye-grass clover pastures over a two-month period from mid-May to mid-July (Figs. S22A; S23A; Tab. S18). So, only 20 % of the colonies survived over the simulation time under the semi-realistic landscape settings. In contrast, for a cropping systems consisting of oilseed rape, sunflower and rye-grass clover none of both patch size and distance does matter, because the late-seasonal gap in forage supply after sunflower ceases flowering result in colony collapse within the first year under both artificial and semi-realistic landscape settings (Figs. S22B; S23B; Tab. S18).

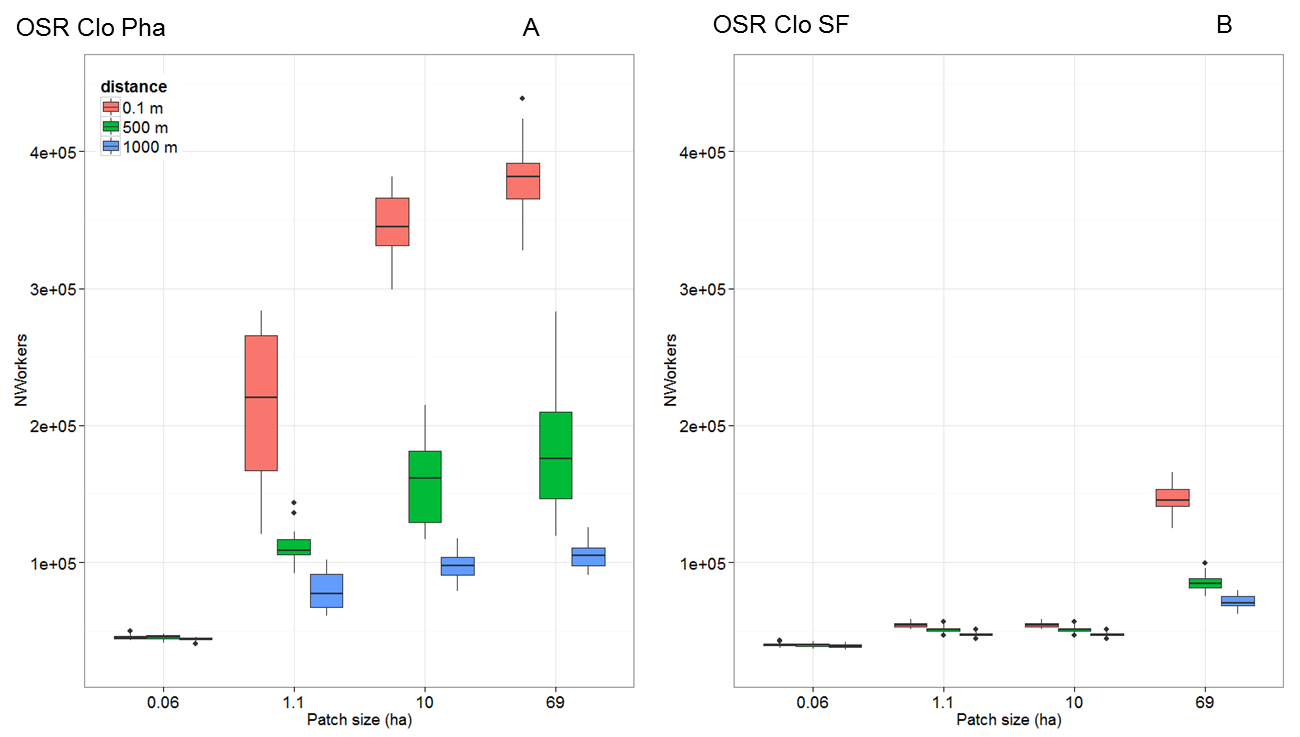

**Figure S23**: Effects of increasing patch size and foraging distance on colony’s productivity of force of worker bees over simulation time of five years or until the colony became extinct (N_Workers_, y-axis) for 3 crop cropping systems (A) oilseed rape, ryegrass-clover, phacelia (OSR Clo Pha) and (B) oilseed rape, ryegrass-clover, sunflower (OSR Clo SF) in a hypothetical landscape (one single patch per crop species). Patch sizes are given in ha (x-axis) and foraging distances are illustrated by colored bars.

**Table S18:** Effects of increasing patch size and distance on colony performance in artificial landscapes for cropping systems of three crop species (one single patch per crop species). For all patch sizes and distances of 3 crop species cropping systems survival probability (SurvProb), time to colony extinction (TTE, given in days), colony size (at the end of the fifth year of simulation, only for surviving colonies), honey stores (at the end of the fifth year, only for surviving colonies) and number of produced worker bees over simulation time of five years or until colony became extinct (N_Workers_) are presented.

| Crop composition | Patch size (ha | Distance  (m) | SurvProb | TTE  (days) | Colony size | Honey  (kg) | N_Workers_ |
| --- | --- | --- | --- | --- | --- | --- | --- |
| OSR Clo Pha | 69 | 0.1 | 1 | 1825 | 6480 | 12.4 | 381688 |
|  | 10 |  | 1 | 1825 | 5921 | 9.9 | 347211 |
|  | 1.1 |  | 0.33 | 1413.6 | 4677 | 6.1 | 212128 |
|  | 0.06 |  | 0 | 366 | 0 | 0 | 45497 |
|  | 69 | 500 | 0.1 | 1197.5 | 4521 | 4.3 | 183180 |
|  | 10 |  | 0 | 1061.1 | 0 | 0 | 162293 |
|  | 1.1 |  | 0 | 757 | 0 | 0 | 111043 |
|  | 0.06 |  | 0 | 364.8 | 0 | 0 | 45379 |
|  | 69 | 1000 | 0 | 726.2 | 0 | 0 | 104100 |
|  | 10 |  | 0 | 686.5 | 0 | 0 | 97810 |
|  | 1.1 |  | 0 | 553.8 | 0 | 0 | 79700 |
|  | 0.06 |  | 0 | 361.9 | 0 | 0 | 44223 |
|  | Semi-realistic landscape |  | 0.2 | 1331.2 | 4708 | 5.8 | 210601 |
| OSR Clo SF | 69 | 500 | 0 | 390.1 | 0 | 0 | 85324 |
|  | 10 |  | 0 | 281.5 | 0 | 0 | 51132 |
|  | 1.1 |  | 0 | 281.5 | 0 | 0 | 51132 |
|  | 0.06 |  | 0 | 365 | 0 | 0 | 39623 |
|  | 69 | 1000 | 0 | 355.4 | 0 | 0 | 71253 |
|  | 10 |  | 0 | 269.7 | 0 | 0 | 47270 |
|  | 1.1 |  | 0 | 269.7 | 0 | 0 | 47270 |
|  | 0.06 |  | 0 | 365 | 0 | 0 | 39034 |
|  | 69 | 0.1 | 0 | 745.2 | 0 | 0 | 146591 |
|  | 10 |  | 0 | 287.1 | 0 | 0 | 44392 |
|  | 1.1 |  | 0 | 287.1 | 0 | 0 | 44392 |
|  | 0.06 |  | 0 | 365 | 0 | 0 | 40033 |
|  | Semi-realistic landscape |  | 0 | 373.5 | 0 | 0 | 82015 |

**Table S19:** Patch sizes and foraging distances for 3 crop cropping systems oilseed rape, ryegrass-clover, phacelia (OSR Clo Pha) and oilseed rape, ryegrass-clover, sunflower (OSR Clo SF) under semi-realistic landscape settings (simple-structured landscape). For each crop species included in the cropping system its number of patches within 1000 m and 500 m foraging range (# patches < 1000 m, # patches < 500 m), its range of foraging distances from the hive to the crop patches (range distance), its number of patches above 10 ha patch size (# patches < 10 ha), its ranges of patch sizes (range patch size) and its number of patches above 10 ha within 1000 m foraging distance (# patches > 10 ha and < 1000 m), is listed.

| Crop | # patches < 1000m | # patches < 500m | Range distance (m) | # patches > 10 ha | Range patch size (ha) | # patches > 10 ha and < 1000 m |
| --- | --- | --- | --- | --- | --- | --- |
| OSR Clo Pha | | | | | | |
| OSR | 5 | 0 | 606.2 – 3730.1 | 22 | 1.1 – 45.2 | 5 |
| Clo | 4 | 2 | 0.1 – 3530.4 | 25 | 1.1 – 69 | 4 |
| Pha | 9 | 3 | 111.3 – 3660.7 | 23 | 1.2 – 59.7 | 9 |
| OSR Clo SF |  |  |  |  |  |  |
| OSR | 7 | 4 | 0.1 – 3545.7 | 24 | 1.2 – 39.5 | 7 |
| Clo | 3 | 1 | 370.1 – 3730.1 | 21 | 1.1 – 59.7 | 3 |
| SF | 8 | 0 | 513.2 – 3594.7 | 25 | 1.3 – 69 | 8 |

**4 crop cropping system scenarios**

As exemplary scenarios for 4 crop cropping systems, we chose two scenarios with 80 - 100 % survival probability (e.g. cropping system composed of ryegrass-dandelion pasture, ryegrass-clover pasture, buckwheat and phacelia), two scenarios with 3.3 – 36.7 % survival probability (e.g. ryegrass-clover pasture, oilseed rape, sunflower and phacelia), and three cropping scenarios with 0 % survival probability (e.g. cereals, oilseed rape, maize and sunflower) for all abundances under our semi-realistic landscape settings. For the presented scenarios we analysed the effects of increasing patch size and increasing foraging distance under hypothetical landscape settings.

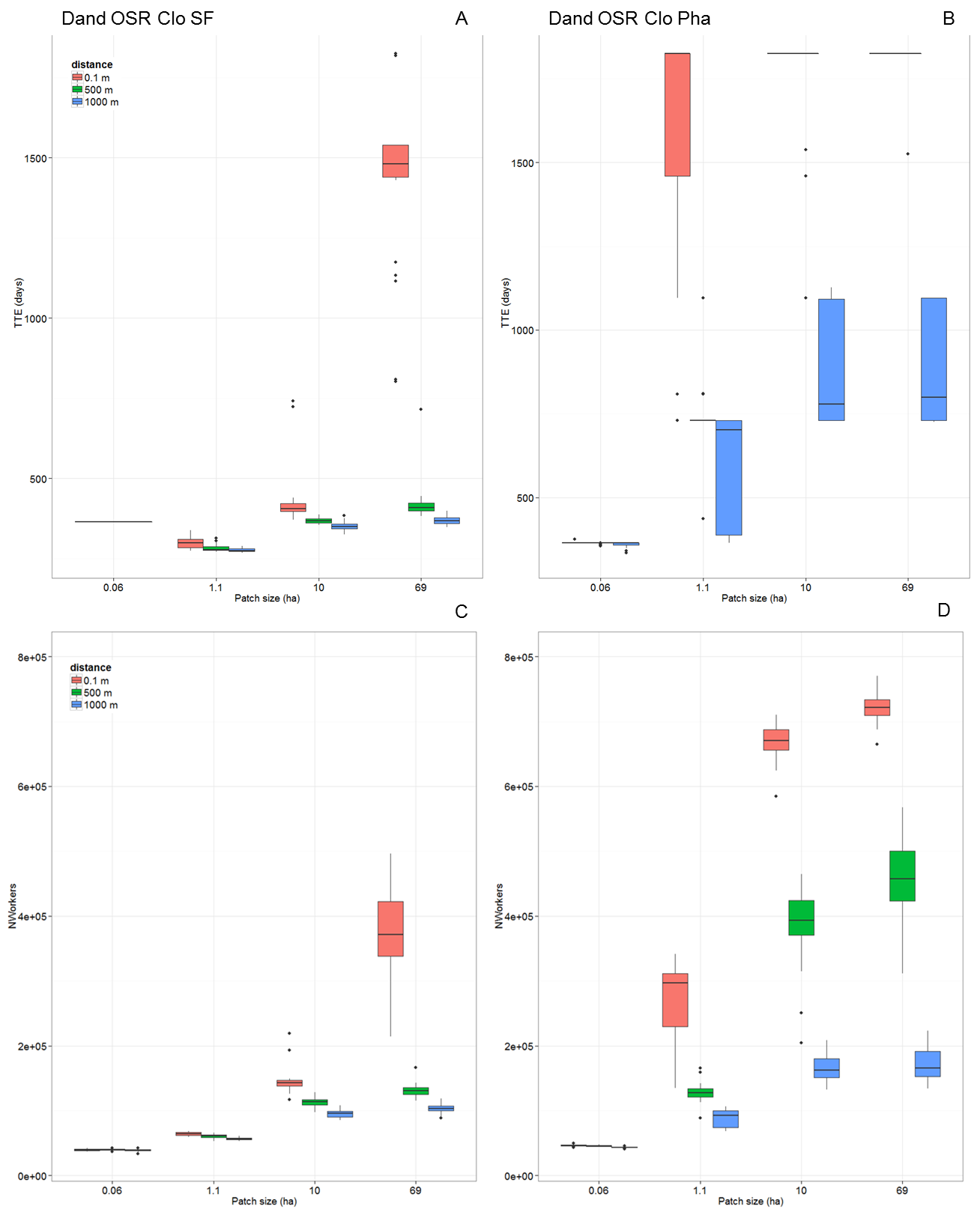

**Figure S24:** Effects of increasing patch size and foraging distance on time to colony extinction (TTE, given in days – upper panels) and colony’s productivity of force of worker bees over simulation time of five years or until the colony became extinct (N_Workers_ - lower panels) for 4 crop cropping systems **(A, C)** ryegrass-dandelion, oilseed rape, ryegrass-clover, sunflower (Dand OSR Clo SF) and **(B, D)** ryegrass-dandelion, oilseed rape, ryegrass-clover, phacelia (Dand OSR Clo Pha) in a hypothetical landscape (one single patch per crop species). Patch sizes are given in ha (x-axis) and foraging distances are illustrated by colored bars.

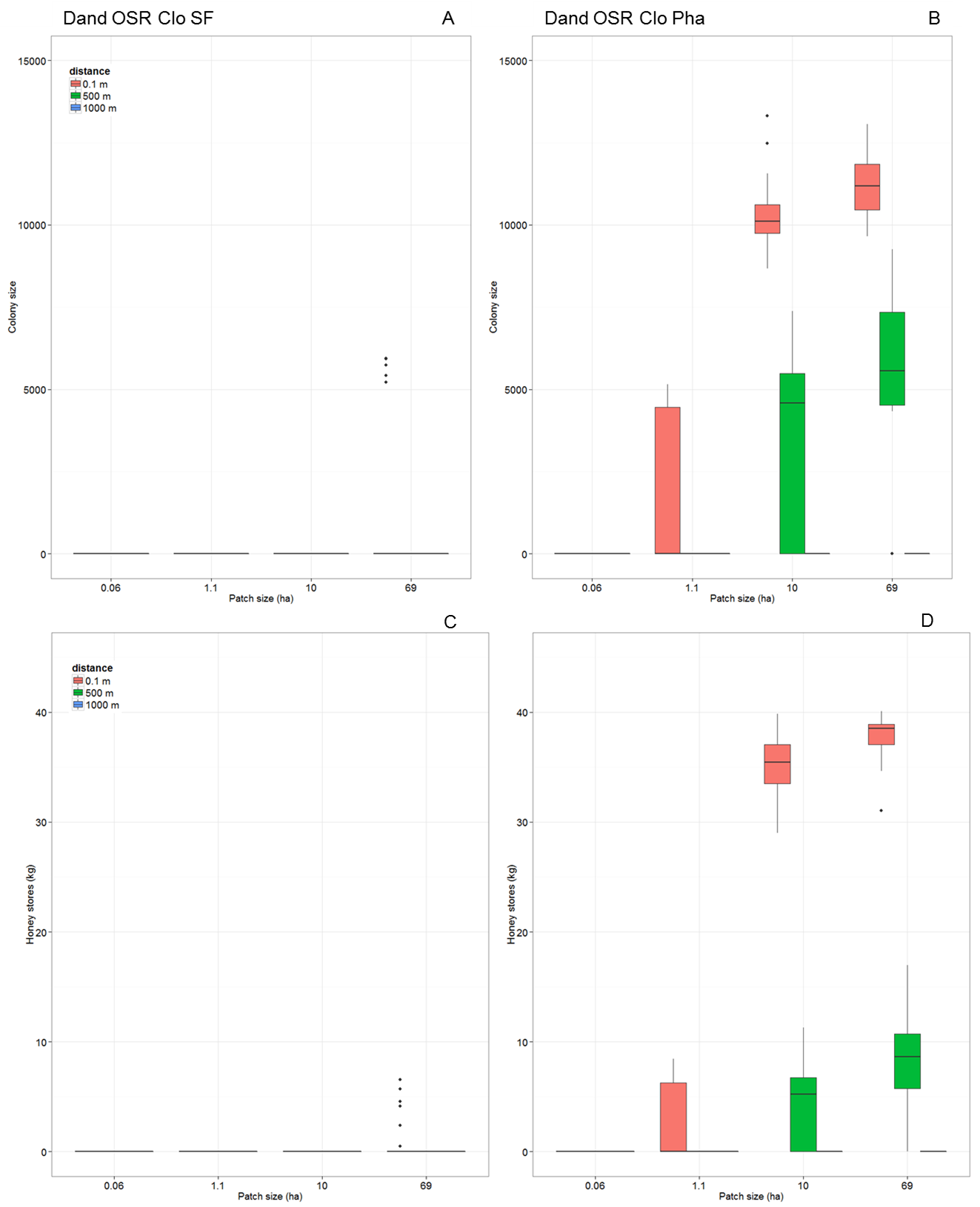

**Figure S25:** Effects of increasing patch size and foraging distance on colony size at the end of the fifth year of simulation time (Colony size - upper panels) and colony’s honey stores at the end of the fifth year of simulation (Honey stores, given in kg - lower panels) for 4 crop cropping systems **(A, C)** ryegrass-dandelion, oilseed rape, ryegrass-clover, sunflower (Dand OSR Clo SF) and **(B, D)** ryegrass-dandelion, oilseed rape, ryegrass-clover, phacelia (Dand OSR Clo Pha) in a hypothetical landscape (one single patch per crop species). Patch sizes are given in ha (x-axis) and foraging distances are illustrated by colored bars.

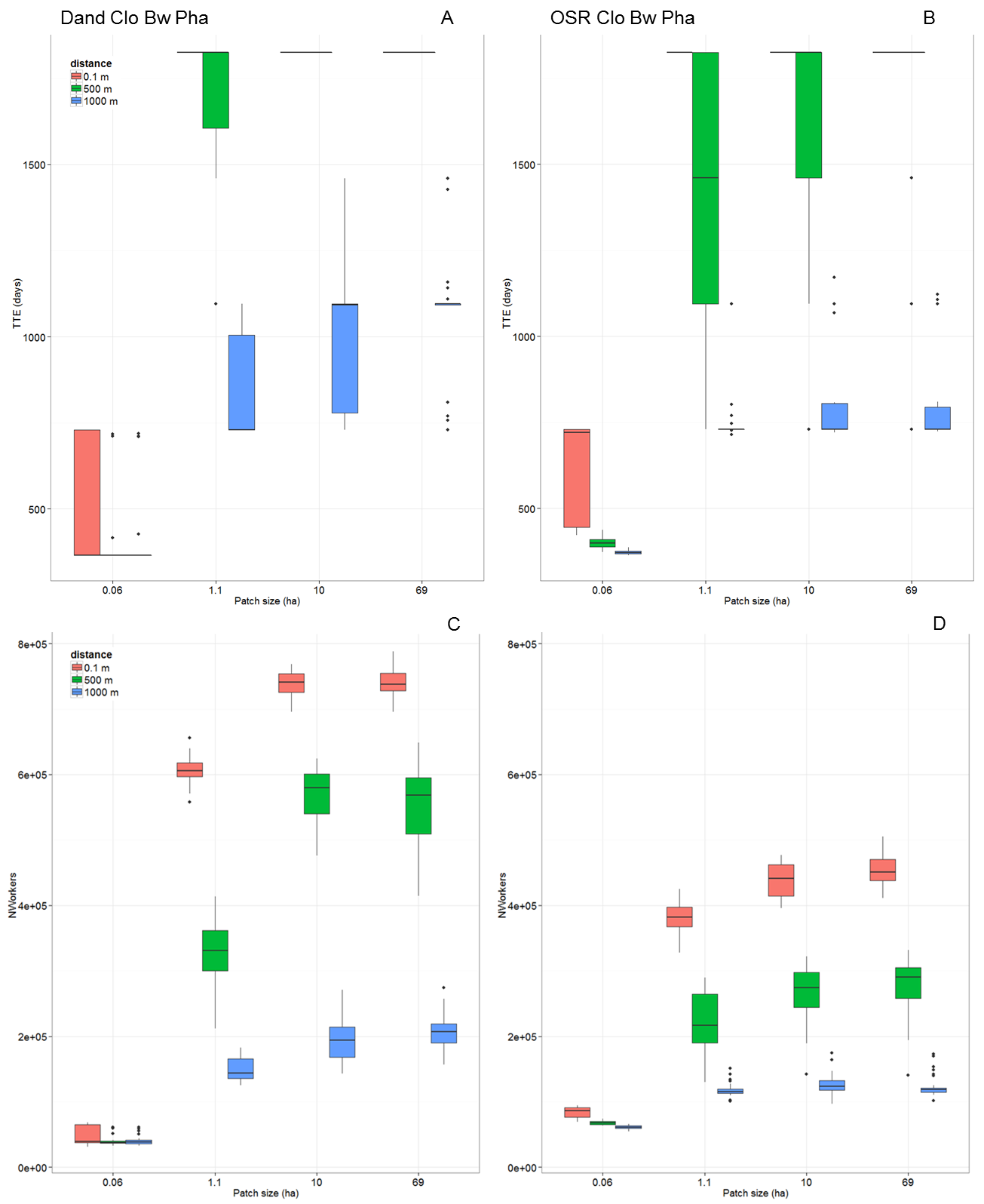

**Figure S26:** Effects of increasing patch size and foraging distance on time to colony extinction (TTE, given in days – upper panels) and colony’s productivity of force of worker bees over simulation time of five years or until the colony became extinct (N_Workers_ - lower panels) for 4 crop cropping systems **(A, C)** ryegrass-dandelion, ryegrass-clover, buckwheat, phacelia (Dand Clo Bw Pha) and **(B, D)** oilseed rape, ryegrass-clover, buckwheat, phacelia (OSR Clo Bw Pha) in a hypothetical landscape (one single patch per crop species). Patch sizes are given in ha (x-axis) and foraging distances are illustrated by colored bars.

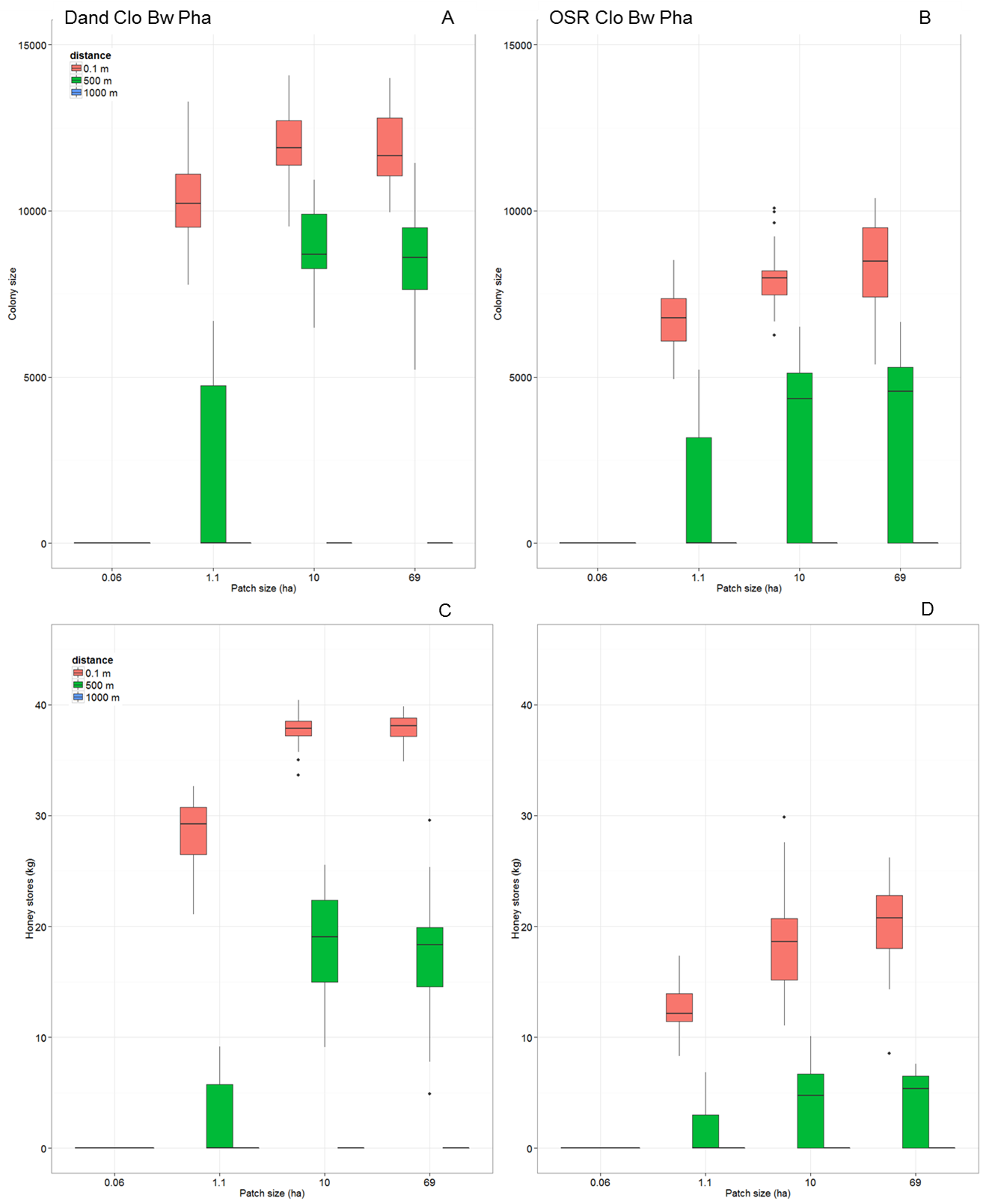

**Figure S27:** Effects of increasing patch size and foraging distance on colony size at the end of the fifth year of simulation time (Colony size - upper panels) and colony’s honey stores at the end of the fifth year of simulation (Honey stores, given in kg - lower panels) for 4 crop cropping systems **(A, C)** ryegrass-dandelion, ryegrass-clover, buckwheat, phacelia (Dand Clo Bw Pha) and **(B, D)** oilseed rape, ryegrass-clover, buckwheat, phacelia (OSR Clo Bw Pha) in a hypothetical landscape (one single patch per crop species). Patch sizes are given in ha (x-axis) and foraging distances are illustrated by colored bars.

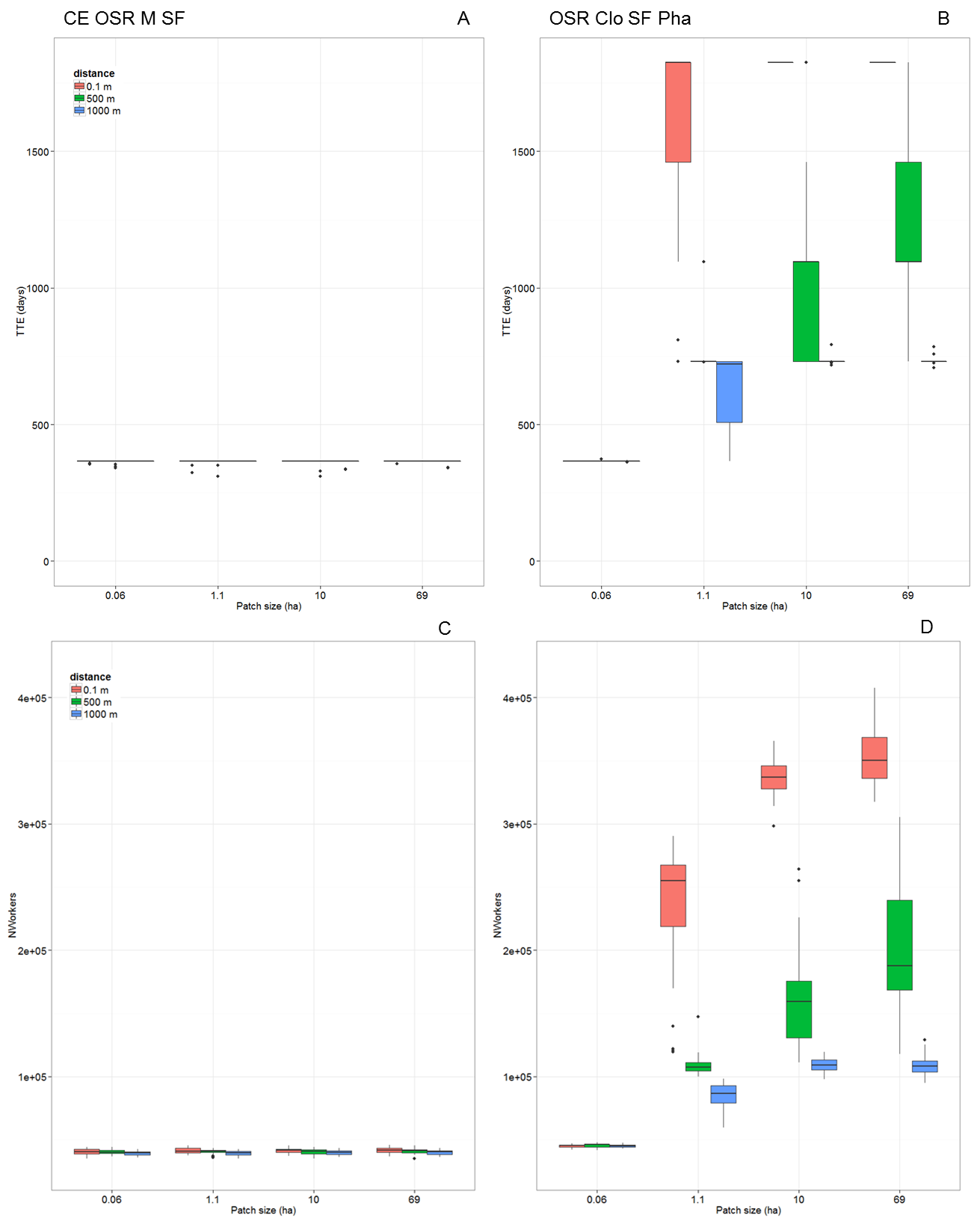

**Figure S28:** Effects of increasing patch size and foraging distance on time to colony extinction (TTE, given in days – upper panels) and colony’s productivity of force of worker bees over simulation time of five years or until the colony became extinct (N_Workers_ - lower panels) for 4 crop cropping systems **(A, C)** cereals, oilseed rape, maize, sunflower (CE OSR M SF) and **(B, D)** oilseed rape, ryegrass-clover, sunflower, phacelia (OSR Clo SF Pha) in a hypothetical landscape (one single patch per crop species). Patch sizes are given in ha (x-axis) and foraging distances are illustrated by colored bars.

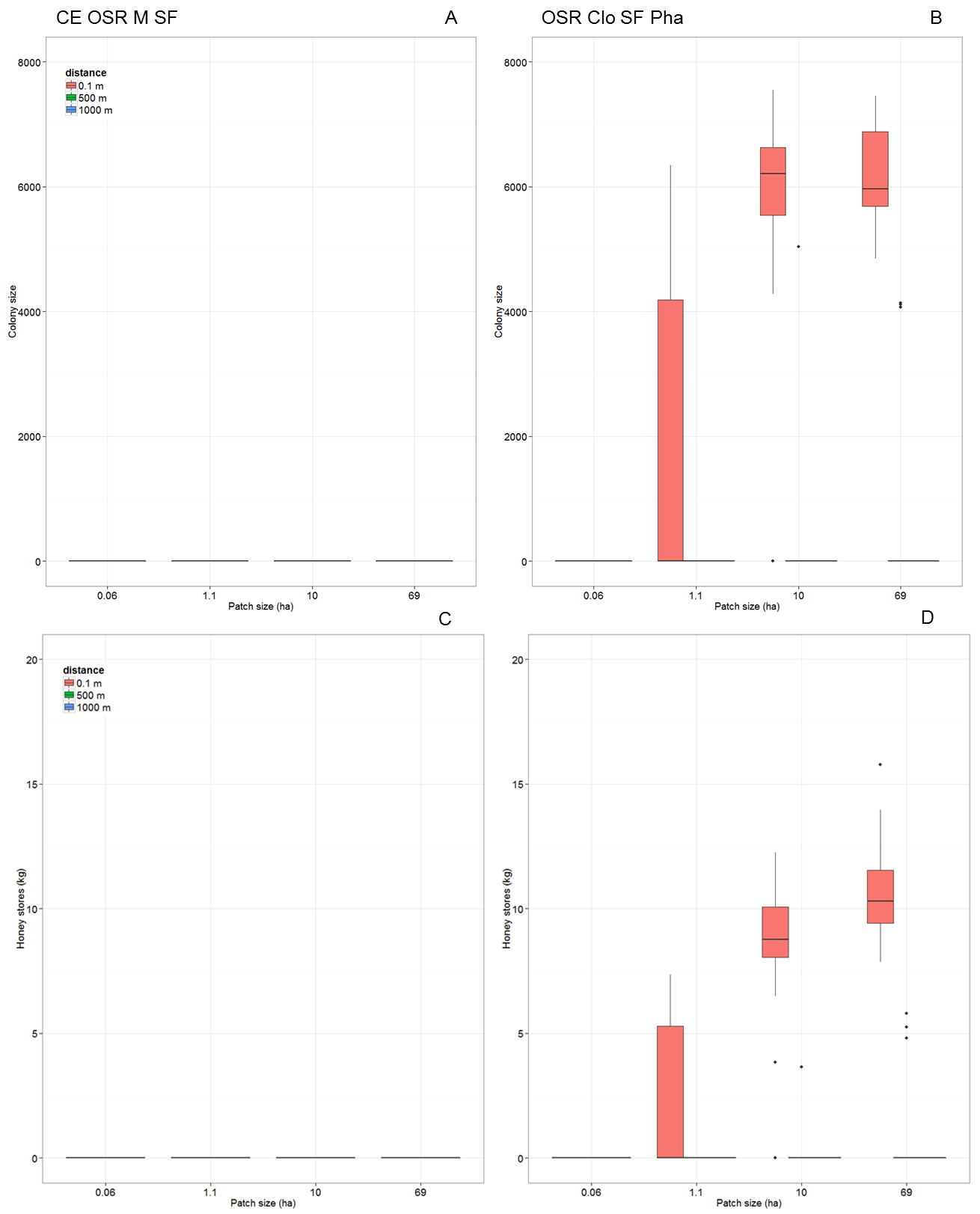

**Figure S29:** Effects of increasing patch size and foraging distance on colony size at the end of the fifth year of simulation time (Colony size - upper panels) and colony’s honey stores at the end of the fifth year of simulation (Honey stores, given in kg - lower panels) for 4 crop cropping systems **(A, C)** cereals, oilseed rape, maize, sunflower (CE OSR M SF) and **(B, D)** oilseed rape, ryegrass-clover, sunflower, phacelia (OSR Clo SF Pha) in a hypothetical landscape (one single patch per crop species). Patch sizes are given in ha (x-axis) and foraging distances are illustrated by colored bars.

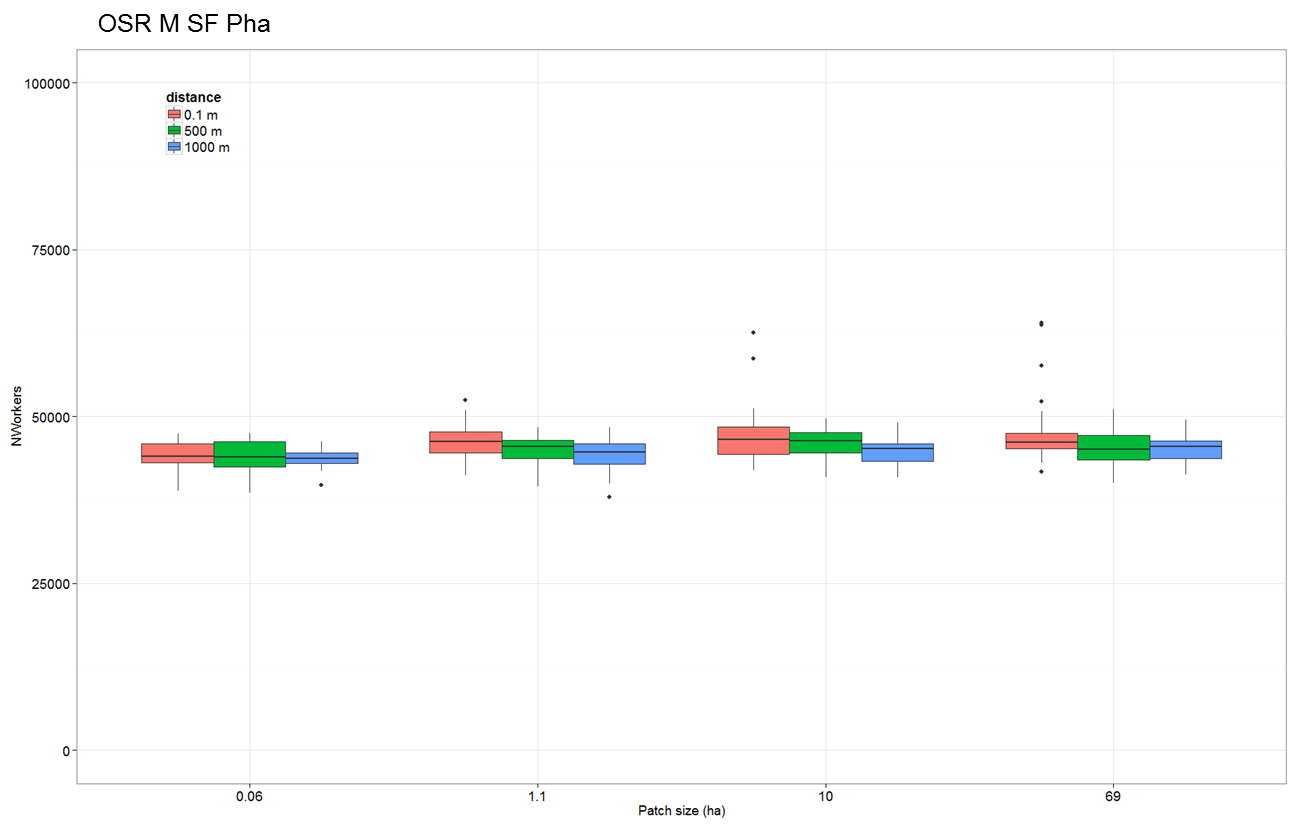

**Figure S30:** Effects of increasing patch size and foraging distance on colony’s productivity of force of worker bees over simulation time of five years or until the colony became extinct (N_Workers_) for the 4 crop cropping systems oilseed rape, maize, sunflower, phacelia (OSR M SF Pha) in a hypothetical landscape (one single patch per crop species). Patch sizes are given in ha (x-axis) and foraging distances are illustrated by colored bars.

**Table S20:** Effects of increasing patch size and distance on colony performance in hypothetical landscapes for cropping systems of four crop species (one single patch per crop species). For all patch sizes and distances of 4 crop species cropping systems survival probability (SurvProb), time to colony extinction (TTE, given in days), colony size (at the end of the fifth year of simulation, only for surviving colonies), honey stores (at the end of the fifth year, only for surviving colonies) and number of produced worker bees over simulation time of five years or until colony became extinct (N_Workers_) are presented.

| Crop composition | Patch size  (ha) | Distance  (m) | SurvProb | TTE  (days) | Colony size | Honey  (kg) | N_Workers_ |
| --- | --- | --- | --- | --- | --- | --- | --- |
| Dand OSR Clo SF | 69 | 0.1 | 0.2 | 1471.4 | 5577 | 4 | 371276 |
|  | 10 | 0.1 | 0 | 428.9 | 0 | 0 | 145277 |
|  | 1.1 | 0.1 | 0 | 296.6 | 0 | 0 | 64159 |
|  | 0.06 | 0.1 | 0 | 365 | 0 | 0 | 39410 |
|  | 69 | 500 | 0 | 419.1 | 0 | 0 | 131112 |
|  | 10 | 500 | 0 | 367.8 | 0 | 0 | 113624 |
|  | 1.1 | 500 | 0 | 282.2 | 0 | 0 | 60691 |
|  | 0.06 | 500 | 0 | 365 | 0 | 0 | 39865 |
|  | 69 | 1000 | 0 | 367.6 | 0 | 0 | 103685 |
|  | 10 | 1000 | 0 | 350.6 | 0 | 0 | 95569 |
|  | 1.1 | 1000 | 0 | 276.9 | 0 | 0 | 56684 |
|  | 0.06 | 1000 | 0 | 365 | 0 | 0 | 38876 |
|  | semi-realistic | |  |  |  |  |  |
| Dand OSR Clo Pha | 69 | 0.1 | 1 | 1825 | 11183 | 37.8 | 722011 |
|  | 10 | 0.1 | 1 | 1825 | 10269 | 35 | 668780 |
|  | 1.1 | 0.1 | 0.43 | 1586.7 | 4607 | 6.6 | 270525 |
|  | 0.06 | 0.1 | 0 | 365.4 | 0 | 0 | 46762 |
|  | 69 | 500 | 0.83 | 1815 | 6403 | 9.3 | 459504 |
|  | 10 | 500 | 0.7 | 1754.6 | 5139 | 6.5 | 388796 |
|  | 1.1 | 500 | 0 | 762 | 0 | 0 | 129251 |
|  | 0.06 | 500 | 0 | 364 | 0 | 0 | 45949 |
|  | 69 | 1000 | 0 | 887 | 0 | 0 | 171382 |
|  | 10 | 1000 | 0 | 858.7 | 0 | 0 | 166005 |
|  | 1.1 | 1000 | 0 | 568.7 | 0 | 0 | 88643 |
|  | 0.06 | 1000 | 0 | 360 | 0 | 0 | 43370 |
|  | semi-realistic | |  |  |  |  |  |
| Dand Clo Bw Pha | 69 | 0.1 | 1 | 1825 | 11856 | 37.9 | 741949 |
|  | 10 | 0.1 | 1 | 1825 | 11927 | 37.8 | 738025 |
|  | 1.1 | 0.1 | 1 | 1825 | 10315.8 | 28.3 | 605973 |
|  | 0.06 | 0.1 | 0 | 486.4 | 0 | 0 | 47210 |
|  | 69 | 500 | 1 | 1825 | 8372 | 17.2 | 551600 |
|  | 10 | 500 | 1 | 1825 | 8973 | 18.7 | 566522 |
|  | 1.1 | 500 | 0.47 | 1705.7 | 5085 | 6.3 | 330339 |
|  | 0.06 | 500 | 0 | 290 | 0 | 0 | 39487 |
|  | 69 | 1000 | 0 | 1054.2 | 0 | 0 | 204832 |
|  | 10 | 1000 | 0 | 979.9 | 0 | 0 | 192929 |
|  | 1.1 | 1000 | 0 | 835.3 | 0 | 0 | 148237 |
|  | 0.06 | 1000 | 0 | 401.9 | 0 | 0 | 40554 |
|  | semi-realistic | |  |  |  |  |  |
| OSR Clo Bw Pha | 69 | 0.1 | 1 | 1825 | 8350 | 20.1 | 454072 |
|  | 10 | 0.1 | 1 | 1825 | 8000 | 18.3 | 437639 |
|  | 1.1 | 0.1 | 1 | 1825 | 6836 | 12.5 | 380973 |
|  | 0.06 | 0.1 | 0 | 613.7 | 0 | 0 | 84082 |
|  | 69 | 500 | 0.67 | 1679 | 5114 | 6.1 | 275210 |
|  | 10 | 500 | 0.53 | 1623 | 5142 | 6.7 | 264988 |
|  | 1.1 | 500 | 0.27 | 1449.8 | 4753 | 5.8 | 222947 |
|  | 0.06 | 500 | 0 | 399.7 | 0 | 0 | 67699 |
|  | 69 | 1000 | 0 | 793.2 | 0 | 0 | 124008 |
|  | 10 | 1000 | 0 | 821.9 | 0 | 0 | 127637 |
|  | 1.1 | 1000 | 0 | 758 | 0 | 0 | 117706 |
|  | 0.06 | 1000 | 0 | 372 | 0 | 0 | 60995 |
|  | semi-realistic | |  |  |  |  |  |
| CE OSR M SF | 69 | 0.1 | 0 | 365 | 0 | 0 | 41672 |
|  | 10 | 0.1 | 0 | 365 | 0 | 0 | 41589 |
|  | 1.1 | 0.1 | 0 | 361.7 | 0 | 0 | 41535 |
|  | 0.06 | 0.1 | 0 | 364.4 | 0 | 0 | 40676 |
|  | 69 | 500 | 0 | 365 | 0 | 0 | 40889 |
|  | 10 | 500 | 0 | 362 | 0 | 0 | 40743 |
|  | 1.1 | 500 | 0 | 362.7 | 0 | 0 | 40414 |
|  | 0.06 | 500 | 0 | 362.9 | 0 | 0 | 40370 |
|  | 69 | 1000 | 0 | 363.4 | 0 | 0 | 40088 |
|  | 10 | 1000 | 0 | 363 | 0 | 0 | 39914 |
|  | 1.1 | 1000 | 0 | 365 | 0 | 0 | 39514 |
|  | 0.06 | 1000 | 0 | 365 | 0 | 0 | 39368 |
|  | semi-realistic | |  |  |  |  |  |
| OSR Clo SF Pha | 69 | 0.1 | 1 | 1825 | 6190 | 10.6 | 354092 |
|  | 10 | 0.1 | 0.97 | 1825 | 6151 | 8.9 | 335535 |
|  | 1.1 | 0.1 | 0.3 | 1550.3 | 4778 | 6.2 | 232070 |
|  | 0.06 | 0.1 | 0 | 365 | 0 | 0 | 45142 |
|  | 69 | 500 | 0.1 | 1268.6 | 4104 | 5.3 | 199381 |
|  | 10 | 500 | 0.03 | 1041.8 | 5034 | 3.6 | 162882 |
|  | 1.1 | 500 | 0 | 742.1 | 0 | 0 | 108802 |
|  | 0.06 | 500 | 0 | 365.3 | 0 | 0 | 45567 |
|  | 69 | 1000 | 0 | 731.4 | 0 | 0 | 108648 |
|  | 10 | 1000 | 0 | 731.4 | 0 | 0 | 109365 |
|  | 1.1 | 1000 | 0 | 635.8 | 0 | 0 | 84768 |
|  | 0.06 | 1000 | 0 | 364.5 | 0 | 0 | 45047 |
|  | semi-realistic | |  |  |  |  |  |
| OSR M SF Pha | 69 | 0.1 | 0 | 378.1 | 0 | 0 | 47721 |
|  | 10 | 0.1 | 0 | 375.9 | 0 | 0 | 47239 |
|  | 1.1 | 0.1 | 0 | 365 | 0 | 0 | 46245 |
|  | 0.06 | 0.1 | 0 | 365 | 0 | 0 | 44369 |
|  | 69 | 500 | 0 | 365 | 0 | 0 | 45384 |
|  | 10 | 500 | 0 | 365 | 0 | 0 | 45903 |
|  | 1.1 | 500 | 0 | 365 | 0 | 0 | 44872 |
|  | 0.06 | 500 | 0 | 364.7 | 0 | 0 | 44021 |
|  | 69 | 1000 | 0 | 364.5 | 0 | 0 | 45153 |
|  | 10 | 1000 | 0 | 364.8 | 0 | 0 | 44704 |
|  | 1.1 | 1000 | 0 | 364.7 | 0 | 0 | 44288 |
|  | 0.06 | 1000 | 0 | 363.9 | 0 | 0 | 43749 |
|  | semi-realistic | |  |  |  |  |  |

**Table S21:**Patch sizes and foraging distances for 3 crop cropping systems oilseed rape, ryegrass-clover, phacelia (OSR Clo Pha) and oilseed rape, ryegrass-clover, sunflower (OSR Clo SF) under semi-realistic landscape settings (simple-structured landscape). For each crop species included in the cropping system its number of patches within 1000 m and 500 m foraging range (# patches < 1000 m, # patches < 500 m), its range of foraging distances from the hive to the crop patches (range distance), its number of patches above 10 ha patch size (# patches < 10 ha), its ranges of patch sizes (range patch size) and its number of patches above 10 ha within 1000 m foraging distance (# patches > 10 ha and < 1000 m), is listed.

| Crop | # patches < 1000m | # patches < 500m | Range distance (m) | # patches > 10 ha | Range patch size (ha) | # patches > 10 ha and < 1000 m |
| --- | --- | --- | --- | --- | --- | --- |
| Dand OSR Clo SF | | | | | | |
| Dand |  |  |  |  |  |  |
| OSR |  |  |  |  |  |  |
| Clo |  |  |  |  |  |  |
| SF |  |  |  |  |  |  |
| Dand OSR Clo Pha |  |  |  |  |  |  |
| Dand | 5 | 0 | 622.3 - 3594.7 | 14 | 1.11 - 69 | 5 |
| OSR | 6 | 3 | 111.3 – 3730.1 | 22 | 15.5 – 59.7 | 6 |
| Clo | 2 | 1 | 222.7 – 3545.7 | 15 | 1.18 – 39.54 | 2 |
| Pha | 5 | 1 | 0.1 - 3371 | 19 | 1.24 – 39.4 | 5 |
| Dand Clo Bw Pha |  |  |  |  |  |  |
| Dand | 6 | 0 | 606.2 – 3545.7 | 18 | 1.11 – 39.54 | 6 |
| Clo | 4 | 1 | 370.1 – 3730.1 | 18 | 1.86 – 59.74 | 4 |
| Bw | 4 | 2 | 111.3 – 3594.7 | 16 | 1.11 – 15.74 | 4 |
| Pha | 4 | 2 | 0.1 – 3660.7 | 18 | 1.24 - 69 | 4 |
| OSR Clo Bw Pha |  |  |  |  |  |  |
| OSR | 6 | 2 | 0.1 – 3594.7 | 14 | 1.24 – 39.54 | 6 |
| Clo | 3 | 1 | 222.7 – 3545.7 | 17 | 1.11 – 39.22 | 3 |
| Bw | 5 | 1 | 370.1 – 3660.7 | 23 | 1.61 – 59.74 | 5 |
| Pha | 4 | 1 | 111.3 – 3730.1 | 16 | 1.11 - 69 | 4 |
| CE OSR M SF |  |  |  |  |  |  |
| CE | 5 | 0 | 606.2 – 3730.1 | 14 | 1.11 – 34.76 | 5 |
| OSR | 5 | 1 | 0.1 - 3371 | 16 | 1.24 - 69 | 5 |
| M | 2 | 1 | 480.8 – 3660.7 | 21 | 1.55 – 34.47 | 2 |
| SF | 6 | 3 | 111.3 – 3139.8 | 19 | 1.11 – 59.74 | 6 |
| OSR Clo SF Pha |  |  |  |  |  |  |
| OSR | 4 | 2 | 222.7 - 3371 | 18 | 1.11 - 69 | 4 |
| Clo | 4 | 0 | 513.2 – 3660.7 | 14 | 1.18 – 39.47 | 4 |
| SF | 4 | 1 | 480.8 - 3072 | 21 | 1.42 – 28.75 | 4 |
| Pha | 6 | 2 | 0.1 – 3730.1 | 17 | 1.11 – 39.41 | 5 |
| OSR M SF Pha |  |  |  |  |  |  |
| OSR | 6 | 1 | 480.8 – 3660.7 | 21 | 1.12 – 45.24 | 6 |
| M | 3 | 0 | 606.2 – 3530.4 | 18 | 1.11 - 69 | 3 |
| SF | 3 | 2 | 0.1 – 3594.7 | 15 | 1.3 – 39.41 | 3 |
| Pha | 6 | 2 | 111.3 – 3730.1 | 16 | 1.11 – 59.74 | 6 |

**8 crop cropping system scenarios**

As exemplary scenarios for 8 crop cropping systems, we chose two scenarios with 50 - 100 % survival probability (cropping systems composed of cereals, oilseed rape, maize, sunflower, bean, ryegrass-dandelion pasture, ryegrass-clover pasture, phacelia and oilseed rape, maize, sunflower, bean, ryegrass-dandelion pasture, ryegrass-clover pasture, buckwheat, phacelia) and one cropping scenario with 0 % survival probability (cereals, oilseed rape, maize, sunflower, bean, rye-grass, ryegrass-dandelion pasture, ryegrass-clover pasture) for all abundances under our semi-realistic landscape settings. For the presented scenarios we analysed the effects of increasing patch size and increasing foraging distance under hypothetical landscape settings.

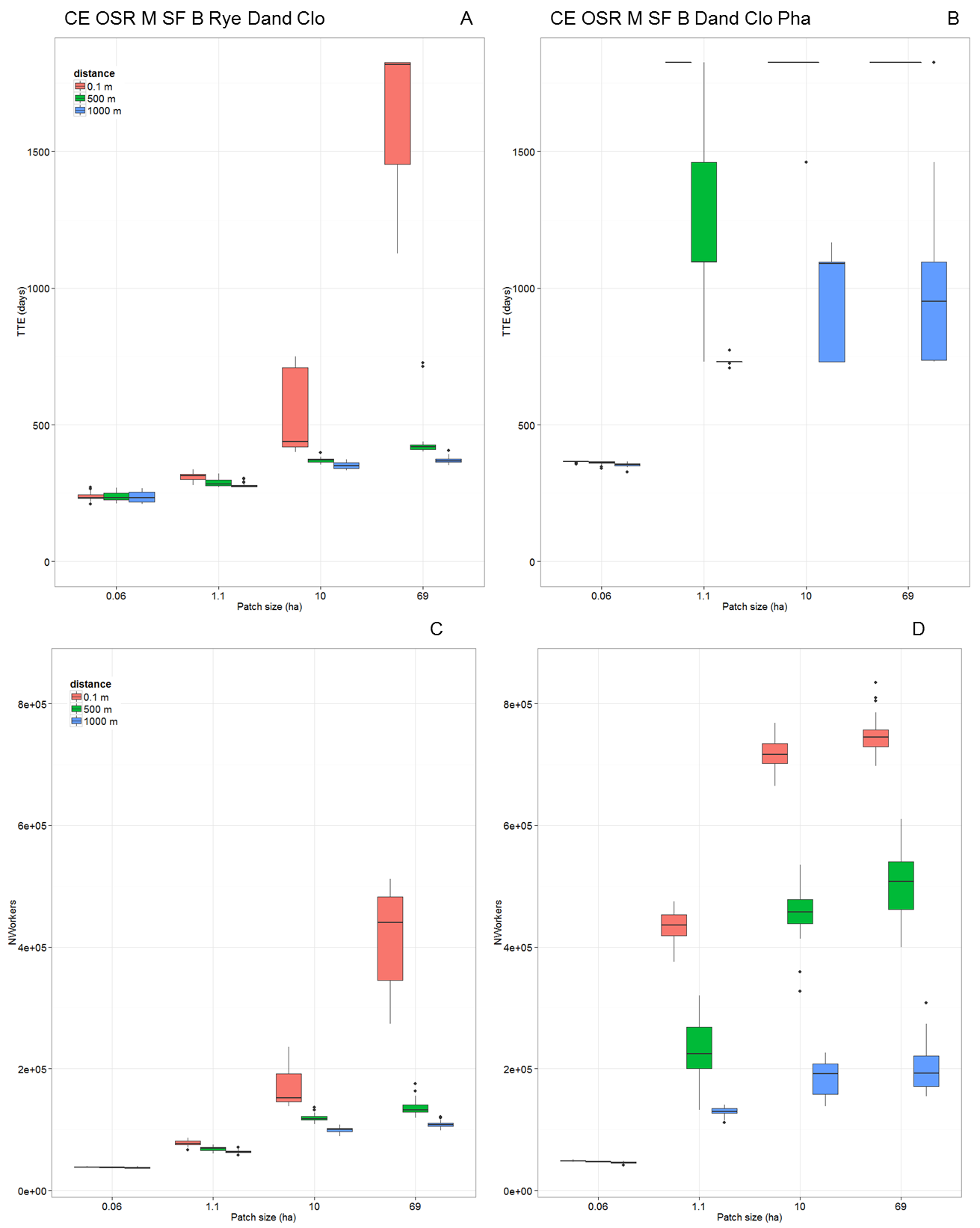

**Figure S31:** Effects of increasing patch size and foraging distance on time to colony extinction (TTE, given in days – upper panels) and colony’s productivity of force of worker bees over simulation time of five years or until the colony became extinct (N_Workers_ - lower panels) for 8 crop cropping systems **(A, C)** cereals, oilseed rape, maize, sunflower, bean, rye-grass, ryegrass-dandelion, ryegrass-clover (CE OSR M SF B Rye Dand Clo) and **(B, D)** cereals, oilseed rape, maize, sunflower, bean, ryegrass-dandelion, ryegrass-clover, phacelia (CE OSR M SF B Dand Clo Pha) in a hypothetical landscape (one single patch per crop species). Patch sizes are given in ha (x-axis) and foraging distances are illustrated by colored bars.

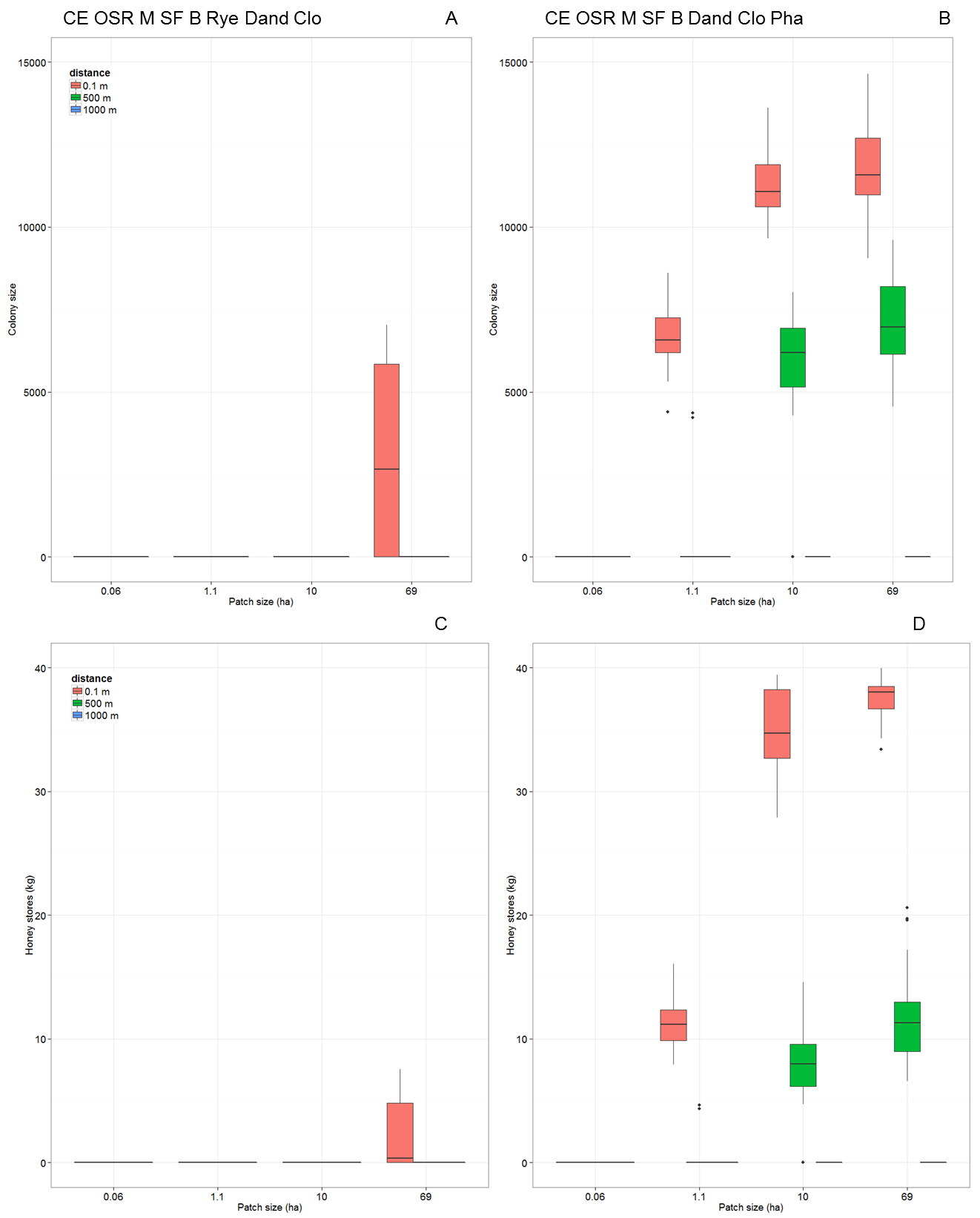

**Figure S32:** Effects of increasing patch size and foraging distance on colony size at the end of the fifth year of simulation time (Colony size - upper panels) and colony’s honey stores at the end of the fifth year of simulation (Honey stores, given in kg - lower panels) for 8 crop cropping systems **(A, C)** cereals, oilseed rape, maize, sunflower, bean, rye-grass, ryegrass-dandelion, ryegrass-clover (CE OSR M SF B Rye Dand Clo) and **(B, D)** cereals, oilseed rape, maize, sunflower, bean, ryegrass-dandelion, ryegrass-clover, phacelia (CE OSR M SF B Dand Clo Pha) in a hypothetical landscape (one single patch per crop species). Patch sizes are given in ha (x-axis) and foraging distances are illustrated by colored bars.

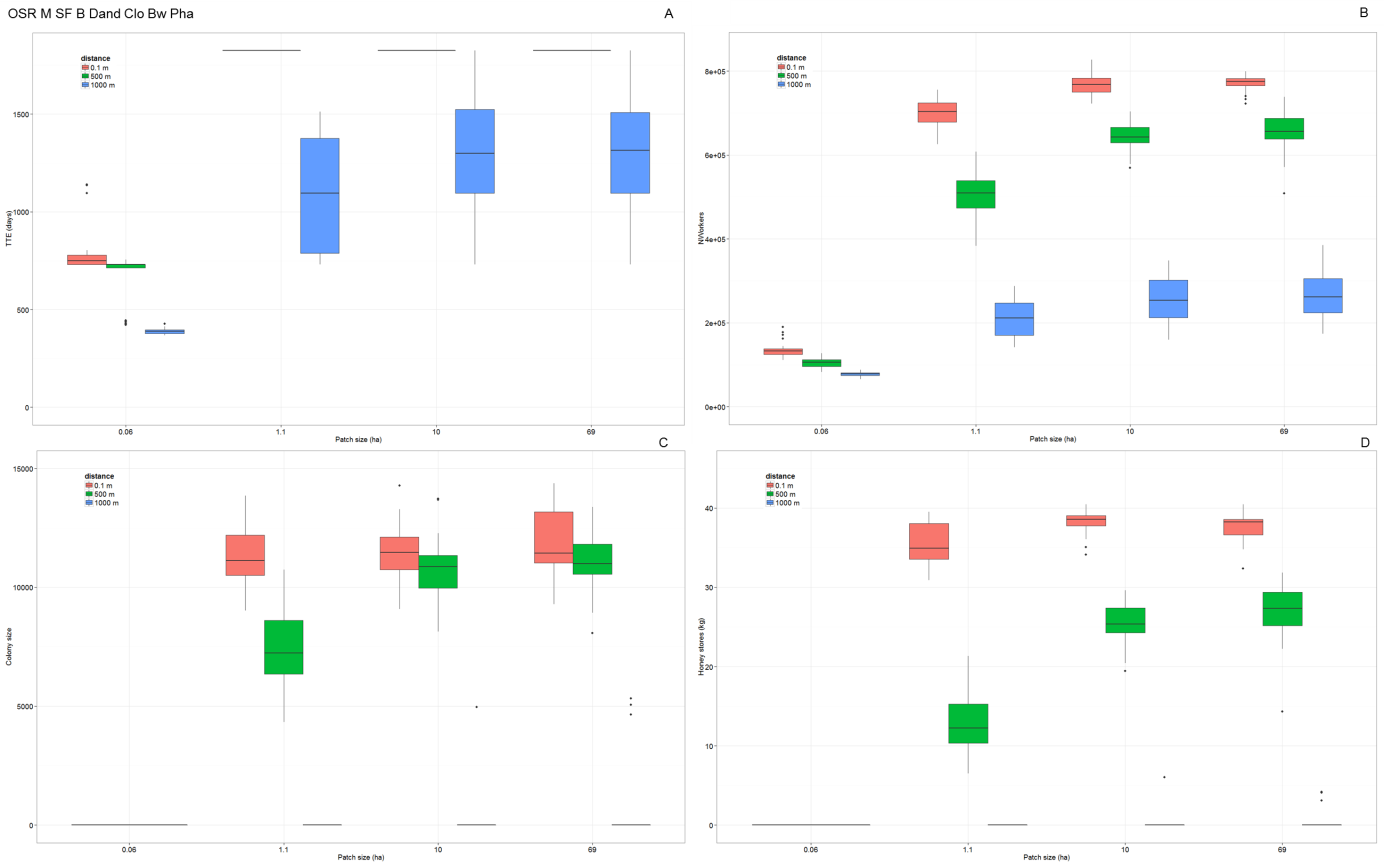

**Figure S33:** Effects of increasing patch size and foraging distance on **(A)** time to colony extinction (TTE, given in days), **(B)** colony’s productivity of force of worker bees over simulation time of five years or until the colony became extinct (N_Workers_), **(C)** colony size at the end of the fifth year of simulation (Colony size) and **(D)** honey stores at the end of the fifth year of simulation (honey stores, given in kg) for the 8 crop cropping systems oilseed rape, maize, sunflower, bean, ryegrass-dandelion, ryegrass-clover, buckwheat, phacelia (OSR M SF B Dand Clo Bw Pha) in a hypothetical landscape (one single patch per crop species). Patch sizes are given in ha (x-axis) and foraging distances are illustrated by colored bars.

**Table S22:** Effects of increasing patch size and distance on colony performance in hypothetical landscapes for cropping systems of eight crop species (one single patch per crop species). For all patch sizes and distances of 8 crop species cropping systems survival probability (SurvProb), time to colony extinction (TTE, given in days), colony size (at the end of the fifth year of simulation, only for surviving colonies), honey stores (at the end of the fifth year, only for surviving colonies) and number of produced worker bees over simulation time of five years or until colony became extinct (N_Workers_) are presented.

| Crop composition | Patch size (ha) | Distance (m) | SurvProb | TTE (days) | Colony size | Honey (kg) | N_Workers_ |
| --- | --- | --- | --- | --- | --- | --- | --- |
| CE OSR M SF B Dand Clo Pha | 69 | 0.1 | 1 | 1825 | 11736 | 37.7 | 748113 |
|  | 69 | 500 | 1 | 1825 | 7089 | 11.6 | 510463 |
|  | 69 | 1000 | 0 | 970.5 | 0 | 0 | 199255 |
|  | 10 | 0.1 | 1 | 1825 | 11254 | 35 | 716534 |
|  | 10 | 500 | 0.93 | 1812.8 | 6250 | 8.5 | 456311 |
|  | 10 | 1000 | 0 | 934.6 | 0 | 0 | 184254 |
|  | 1.1 | 0.1 | 1 | 1825 | 6644 | 11.2 | 437615 |
|  | 1.1 | 500 | 0.067 | 1291 | 4292 | 4.5 | 231782 |
|  | 1.1 | 1000 | 0 | 730.4 | 0 | 0 | 129541 |
|  | 0.06 | 0.1 | 0 | 364 | 0 | 0 | 48825 |
|  | 0.06 | 500 | 0 | 360.2 | 0 | 0 | 47266 |
|  | 0.06 | 1000 | 0 | 353.9 | 0 | 0 | 45791 |
| CE OSR M SF B Rye Dand Clo | 69 | 0.1 | 0.5 | 1595.8 | 5906 | 4.5 | 419126 |
|  | 69 | 500 | 0 | 438.2 | 0 | 0 | 136117 |
|  | 69 | 1000 | 0 | 370.3 | 0 | 0 | 108108 |
|  | 10 | 0.1 | 0 | 533 | 0 | 0 | 169961 |
|  | 10 | 500 | 0 | 369.8 | 0 | 0 | 119211 |
|  | 10 | 1000 | 0 | 351 | 0 | 0 | 99511 |
|  | 1.1 | 0.1 | 0 | 309.3 | 0 | 0 | 77598 |
|  | 1.1 | 500 | 0 | 289.8 | 0 | 0 | 68602 |
|  | 1.1 | 1000 | 0 | 278.3 | 0 | 0 | 63525 |
|  | 0.06 | 0.1 | 0 | 238.5 | 0 | 0 | 38221 |
|  | 0.06 | 500 | 0 | 237.9 | 0 | 0 | 37611 |
|  | 0.06 | 1000 | 0 | 237.6 | 0 | 0 | 37035 |
| OSR M SF B Dand Clo Bw Pha | 69 | 0.1 | 1 | 1825 | 12001 | 37.6 | 771891 |
|  | 69 | 500 | 1 | 1825 | 11169 | 27 | 657064 |
|  | 69 | 1000 | 0.1 | 1354.2 | 5004 | 3.8 | 270947 |
|  | 10 | 0.1 | 1 | 1825 | 11473 | 38.3 | 766970 |
|  | 10 | 500 | 1 | 1825 | 10763 | 25.5 | 645986 |
|  | 10 | 1000 | 0.033 | 1303.9 | 4970 | 6 | 252886 |
|  | 1.1 | 0.1 | 1 | 1825 | 11248 | 35.7 | 700976 |
|  | 1.1 | 500 | 1 | 1825 | 7471 | 13 | 508174 |
|  | 1.1 | 1000 | 0 | 1086.4 | 0 | 0 | 212152 |
|  | 0.06 | 0.1 | 0 | 798.4 | 0 | 0 | 135530 |
|  | 0.06 | 500 | 0 | 659.8 | 0 | 0 | 103933 |
|  | 0.06 | 1000 | 0 | 389.2 | 0 | 0 | 78158 |

**Details on landscape settings of the exemplary discontinuous and continuous forage supply scenarios**

Table S23:

| **Crop species** | **# patches** | **Range patch size (ha)** | **Mean patch size (ha)** | **# patches within patch size class** | | |
| --- | --- | --- | --- | --- | --- | --- |
|  |  |  |  | **<5 ha** | **5 ha – 10 ha** | **>10 ha** |
| ***4 crops – discontinuous forage supply:*** CE OSR M SF (each 25 % abundance on agricultural area) | | | | | | |
| CE | 42 | 1.1 – 34.8 | 9.5 | 17 | 11 | 14 |
| OSR | 43 | 1.2 – 39.5 | 11.4 | 10 | 17 | 16 |
| M | 43 | 1.5 - 69 | 10.4 | 10 | 12 | 21 |
| SF | 43 | 1.9 – 59.7 | 11.9 | 10 | 14 | 19 |
| ***8 crops – discontinuos forage supply:*** CE OSR M SF B Rye Dand Clo (each 12.5 % abundance on agricultural area) | | | | | | |
| CE + Rye | 45 | 1.2 – 59.7 | 10.7 | 15 | 13 | 17 |
| OSR | 21 | 1.7 – 34.8 | 9.1 | 7 | 6 | 8 |
| M | 21 | 1.1 – 39.2 | 10.8 | 5 | 8 | 8 |
| SF | 21 | 2.1 - 69 | 12.6 | 5 | 5 | 11 |
| B | 21 | 2.6 – 39.5 | 11.4 | 4 | 7 | 10 |
| Dand | 21 | 1.3 – 39.5 | 10.9 | 5 | 9 | 7 |
| Clo | 21 | 1.1 – 31.3 | 10.3 | 6 | 6 | 9 |
| ***4 crops – continuous forage supply:*** 89% SF Dand Clo 1% Pha (Dand and Clo with 5 % abundance on agricultural area) | | | | | | |
| SF | 151 | 1.1 - 69 | 10.7 | 44 | 46 | 61 |
| Dand | 9 | 2.4 – 18.1 | 11.1 | 1 | 3 | 5 |
| Clo | 9 | 1.7 – 17.4 | 9.1 | 2 | 4 | 3 |
| Pha | 2 | 7.4 – 39.4 | 24.3 | 0 | 1 | 1 |
| ***8 crops – continuous forage supply:* 69%** CE OSR M SF B Dand Clo 1% Pha ( all others 5 % abundance on agricultural area) | | | | | | |
| CE | 115 | 1.1 – 59.7 | 10.2 | 32 | 36 | 47 |
| OSR | 9 | 6.3 – 26.6 | 11.8 | 0 | 4 | 5 |
| M | 9 | 1.8 – 12.8 | 6.3 | 5 | 2 | 2 |
| SF | 9 | 1.2 – 39.2 | 10.4 | 4 | 2 | 3 |
| B | 9 | 2.1 – 39.5 | 14.2 | 1 | 2 | 6 |
| Dand | 9 | 1.8 – 39.5 | 11.3 | 3 | 4 | 2 |
| Clo | 9 | 1.8 - 69 | 16.1 | 1 | 4 | 4 |
| Pha | 2 | 3 – 39.4 | 21.2 | 1 | 0 | 1 |

**Simulations of semi-natural habitats under semi-realistic landscape settings**

We ran simulations for testing effects of semi-natural habitats under semi-realistic landscape settings for two exemplary cropping systems of low diversity (4 crops – cereals, oilseed rape, maize and sunflower) and high diversity (8 crops – cereals, oilseed rape, maize, sunflower, bean, rye-grass, dandelion-ryegrass pasture and clover-ryegrass pasture) resulting in colony extinction within in one (low diversity example) or two years (high diversity example) due to temporal gaps in continuous supply with nectar and pollen. To these semi-realistic cropping systems we assigned semi-natural habitats as one single forage patch of 1 ha size located in 1000 m or 2000 m flight distance to the honeybee hive. We assumed constant nectar and pollen flow from mid-march onwards. We varied quantitative nectar and pollen rewards from 1 l up to 3 l nectar per day and 100 g up to 500 g pollen per day. To represent early spring foraging on days when weather conditions are suitable for foraging flights, we placed a forage patch providing low daily amounts of nectar (1 l) and pollen (500 g) from January until end-March at 1000 m from the hive, which allows the colony to survive the early spring period (see above). We ran simulations for 5 years and for 30 replicates per scenario, and analysed colony performance in terms of survival probability (SurvProb), time to colony extinction (TTE), colony’s productivity of force of worker bees over simulation time or until the colony became extinct (N_Workers_), colony size and honey stores at the end of the fifth simulation year (only for surviving colonies).

**Table S24:** Design of simulations for analyzing effects of semi-natural habitats under semi-realistic landscape settings.

| Crop composition | Semi-natural habitats | | |
| --- | --- | --- | --- |
|  | Distance (m) | Nectar provision (l per day) | Pollen provision (g per day) |
| CE OSR M SF | 1000  2000 | 1  2  3  3  3 | 500  500  100  300  500 |
| CE OSR M SF B Rye Dand Clo | 1000  2000 | 1  2  3  3  3 | 500  500  100  300  500 |

Our results indicated that in a cropping system of low crop diversity with frequent, prolonged gaps in nectar and pollen availability between mass-flowering (4 crops: cereals, oilseed rape, maize, sunflower) as shown in Fig. 5, even the introduction of semi-natural habitats providing 3 l nectar and 500 g pollen per day throughout the year and at intermediate flight distance of 1000 m resulted in 60% survival and very weak adult worker populations (see Fig. S7.20). For 2000 m flight distance to the semi-natural habitat none of 30 modelled colonies survived the five years of simulation time (see Fig. S7.20). For the cropping system of high diversity with one last-seasonal forage gap in late-summer (8 crops: cereals, oilseed rape, maize, sunflower, bean, rye-grass pasture, rye-grass dandelion infested pasture, trefoil-grass pasture as shown in Fig. 5), the introduction of semi-natural habitats providing 3 l nectar and 100 up to 500 g pollen per day and at intermediate flight distance of 1000 m was sufficient to keep modelled colonies in well-situated foraging conditions (100% survival; N_Workers_ ≥ 500000), but if the semi-natural habitat patch is further away from the hive (2000 m) none of the modelled colonies survived the simulation time (see Fig. S7.20).

### S8 Current limitations

There are a number of ways in which our presented modelling framework could be improved to better simulate colony response to forage availability in agricultural landscape in a more realistic way. The most obvious is that so far the agricultural systems that we considered are simplified in their crop composition, rotation and abundance of floral resources, and in the representation of semi-natural habitats. Data about flowering phenology and nectar and pollen rewards for different crop species and plant species of semi-natural habitats are scarce and hard to distil from literature and vegetation data bases (Baude et al. 2016). We therefore focused on a selection of the most important crops, pasture types, and minor and cover crops and we assumed constant crop species-specific nectar and pollen content over the blooming period. Due to the high intraspecific variation in reported data, we used values which are likely to be linked with good weather throughout the year, because the weather data we used ("Rothamsted (2011)") represent largely beneficial conditions. Nevertheless, applied nectar data for crop cultivation are in the range of those reported by Baude et al. (2016).

BEEHAVE neglects differences in pollen qualities (e.g. amino acids) among different species. In reality honeybees rely on a high diversity of floral resources to meet their pollen requirements especially in spring. Higher pollen diversity enhances immunity to diseases and tolerance to pesticides (Di Pasquale et al. 2013) and ensures the health and sustainability of the colony. This cannot currently be reflected by the model.

A further important limitation is that we do not consider competition with other honeybee colonies and wild pollinators. Implementing realistic densities and distributions of colonies or apiaries will considerably increase the complexity of an already complex modelling framework, but this challenge needs to be tackled. Regarding competition with wild pollinators, currently available data, except for common bumblebee and solitary bee species, are scarce and need to be reflected via aggregated and conservative assumptions. The fact that only a small proportion of scenarios considered in our study allows colonies to persist over longer times indicates that forage gaps generated by the cropping system might be buffered by floral resources that we did not consider. Moreover, it is unknown as to how well honeybee colonies would survive in typical agricultural landscapes without apicultural management.

An important purpose of modelling is to better understand the relative importance of different factors, which can steer future data collection efforts and experiments to areas where so far too little data exist. This presented framework combining the landscape generator NePoFarm and the honeybee model BEEHAVE enables us to translate the spatiotemporal distribution of floral resources into the performance of a single honeybee colony. To reduce uncertainty, data on the spatiotemporal resources in crop and semi-natural habitats need to be compiled and collected in a systematic way.

In our analysis, we deliberately ignored other stressors such as pathogens, certain beekeeping practices, and pesticide effects. It is therefore important to keep in mind that the predictions of colony performance presented here are relative, not absolute. Certain cropping systems reduce the frequency, timing and duration of temporary gaps in nectar and pollen supply affecting colony’s life stages and tasks and enable higher viability and better performance than others, and this enables the colonies to cope better with additional stress.
